# Integrative genomics identifies candidate genes underlying trypanotolerance in hybrid African cattle

**DOI:** 10.1101/2025.07.30.667693

**Authors:** Gillian P. McHugo, James A. Ward, Said Ismael Ng’ang’a, Laurent A.F. Frantz, John A. Browne, Michael Salter-Townshend, Grace M. O’Gorman, Kieran G. Meade, Emmeline W. Hill, Thomas J. Hall, David E. MacHugh

## Abstract

Integrative genomics combines data from different ‘omic sources to link genotypes and phenotypes with the aim of unravelling biological networks and pathways that undergird complex traits, particularly with respect to disease. In this respect, integrative genomics using population and functional genomic data can be employed to understand evolutionary processes that have shaped adaptation to infectious diseases in domestic cattle. This approach can be particularly informative for African cattle, which exhibit a complex mosaic of *Bos taurus* (taurine) and *Bos indicus* (indicine) genomic ancestry. Some African taurine populations have an important evolutionary adaptation known as trypanotolerance, a genetically determined tolerance of infection by trypanosome parasites (*Trypanosoma* spp.) that cause African animal trypanosomiasis (AAT) disease. AAT is one of the largest constraints to livestock production in sub-Saharan Africa and causes a financial burden of approximately $4.5 billion annually. In this study we identified putative candidate genes underlying trypanotolerance through the integration of local ancestry inference (LAI) from genome-wide SNP data for multiple trypanotolerant and trypanosusceptible hybrid cattle populations with RNA-seq and expression microarray transcriptomics data from multiple tissues collected across time course trypanosome infection experiments. These candidate genes included *AGO2*, *CBL*, *CNOT1*, *EDN1*, *IL1B*, *NFKB1*, *RIPK1*, and *TRAF2*. Functional analysis of the gene set outputs from this work highlighted GO terms associated with the immune system (including the major histocompatibility complex – MHC) and cell signalling processes. These results signpost future work to elucidate the cellular networks and pathways that drive trypanotolerance.

**Author Summary:** Integrative genomics combines different types of data to identify links between genes and traits, particularly with respect to disease. In this respect, integrative genomics can be used to understand the admixture and adaptation to infectious diseases that have shaped the genomes of domestic cattle. This is particularly noticeable in the case of African cattle, which form a complex mosaic of *Bos taurus* (taurine) and *Bos indicus* (indicine) ancestry. Some African taurine populations exhibit an evolutionary adaptation known as trypanotolerance, a genetically determined tolerance of infection by trypanosome parasites (*Trypanosoma* spp.) that cause African animal trypanosomiasis (AAT) disease. AAT is one of the largest constraints to livestock production in sub-Saharan Africa and causes a financial burden of approximately $4.5 billion annually. In this study we identify potential candidate genes underlying trypanotolerance through integration of subchromosomal genomic ancestry data from multiple trypanotolerant and trypanosusceptible hybrid cattle populations with gene expression data from multiple tissues collected across time course trypanosome infection experiments.

## Introduction

Integrative genomics, an important strategy for obtaining novel scientific insights from several different but complementary omics technologies, has become increasingly cost-effective as generation of genome-scale data has become more economical (Gamazon *et al*. 2013; Koestler *et al*. 2014). An integrative genomics approach can be used to link phenotypic information with multiple types of biomolecular data with the aim of unravelling the biological networks and pathways that underpin complex traits in multicellular organisms (Subramanian *et al*. 2020; Porcu *et al*. 2021). In this regard, an integrative genomics approach encompassing multi-omics analyses and network biology has been acknowledged in recent years as a powerful strategy for understanding the vertebrate immune system and immune responses to pathogens and parasites (Urbanski *et al*. 2019; Hao *et al*. 2020; Wong *et al*. 2021; Wang *et al*. 2023). This is because each individual omic technology is unable to capture the full biological complexity of host-pathogen interaction for many infectious diseases and the integration of multiple omics sources can provide more detailed and comprehensive insights (Karczewski & Snyder 2018; Cano-Gamez & Trynka 2020).

Integrative and functional population genomics techniques have shown that the immune system of modern humans has been shaped by adaptation to pathogens and parasites, regional human ancestry contributions, gene flow from archaic hominins such as Neanderthals and Denisovans, and the emergence of agriculture, sedentarism, and complex societies (Karlsson *et al*. 2014; Quintana-Murci 2019; Randolph *et al*. 2024). For example, European and African ancestry components in modern human populations have differing effects on the activity of innate immune pathways associated with responses to viral infections (Randolph *et al*. 2021). Also, analyses of paleogenomic and GWAS data have shown that the likelihood of avoiding severe COVID-19 caused by infection with SARS-CoV-2 is influenced positively and negatively by Neanderthal haplotypes at human chromosomes 12 (HSA12) and 3 (HSA3), respectively (Zeberg & Paabo 2020, 2021). In addition, genetic adaptation to pathogens since the development of farming, particularly during the last three thousand years after the Bronze Age, has contributed to inflammatory disease risk in modern European populations (Kerner *et al*. 2023).

Domestic cattle can be categorized into *Bos taurus* (taurine)*, Bos indicus* (indicine), and various grades of hybrid taurine-indicine breeds or populations (Bovine HapMap Consortium *et al*. 2009; Decker *et al*. 2014). The divergence between the taurine and indicine lineages substantially predates the development of animal agriculture with high-resolution genome-scale analyses providing estimates of at least 300 kya for the split between the groups (Chen *et al*. 2018; Wang *et al*. 2018; Wu *et al*. 2018; Hou *et al*. 2024). These diverse ancestries, subsequent admixture, and selection for resistance to infectious diseases caused by viruses, bacteria, and parasites have shaped the evolution of the domestic cattle genome in Africa. Examples include rinderpest disease caused by rinderpest virus (RPV) (Flori *et al*. 2014), tuberculosis caused by *Mycobacterium bovis* (Kassahun *et al*. 2015; Callaby *et al*. 2020), and most notably trypanosomiasis due to bloodstream parasites of the *Trypanosoma* genus (Bahbahani *et al*. 2018; Tijjani *et al*. 2019).

Patterns of genomic variation in African cattle form clines of *B. indicus* and *B. taurus* ancestry primarily across the east-west axis of continent (Hanotte *et al*. 2002). However, at a more granular regional level, nomadic pastoralism, livestock trade, and organised crossbreeding has resulted in a rich diversity of taurine, indicine, and admixed populations (MacHugh *et al*. 1997; Freeman *et al*. 2004; Kim *et al*. 2017a; Kim *et al*. 2020). Importantly, retention in West and Central Africa of substantial African taurine ancestry is largely due to a genetically determined tolerance of the trypanosomes that cause trypanosomiasis, a major impediment to sub-Saharan livestock production (Steverding 2008; Yaro *et al*. 2016). These trypanotolerant cattle exhibit a greater ability to control parasitaemia and anaemia, which makes them more productive in areas infested with trypanosomes than *B. indicus* cattle or other *B. taurus* breeds (Murray *et al*. 1984; Dayo *et al*. 2012; Yaro *et al*. 2016). These trypanotolerant breeds are therefore an important genetic resource as they are uniquely adapted to livestock production in the humid tropical and semitropical regions of Africa (Dayo *et al*. 2012; Food and Agriculture Organization 2023). The genes and genomic regulatory elements (GREs) containing sequence variations that contribute to the polygenic trypanotolerance trait remain poorly understood, although some candidate genes have been proposed (Noyes *et al*. 2011; Alvarez *et al*. 2016; Yaro *et al*. 2016). Better knowledge of the genes and genomic variants responsible for trypanotolerance could inform genome-enabled breeding or genome editing programmes to increase the productivity of livestock populations in sub-Saharan Africa (Yaro *et al*. 2016).

In this study we investigated the genomic architecture of the trypanotolerance trait through the integration of results of local ancestry inference (LAI) using genome-wide SNP data for multiple trypanotolerant and trypanosusceptible hybrid cattle populations with gene expression data from some of the same animals in the form of both RNA-seq and microarray data from time course trypanosome infection experiments.

## Materials and methods

### Data Sources

#### SNP data

As previously described (McHugo *et al*. 2025a), Illumina^®^ BovineHD 777K BeadChip SNP data sets were generated for 39 African cattle (23 Somba, 8 N’Dama and 8 Boran). The N’Dama and Boran animals were obtained from DNA samples collected during a trypanosome infection time course experiment performed in 2003 (O’Gorman *et al*. 2006; Meade *et al*. 2009; O’Gorman *et al*. 2009). These SNP data sets were merged with Illumina^®^ BovineHD 777K BeadChip genome-wide high-density SNP data sets that were assembled from published studies and databases (Sempere *et al*. 2015; Bahbahani *et al*. 2017; Upadhyay *et al*. 2017; Verdugo *et al*. 2019; Barbato *et al*. 2020; Ward *et al*. 2022; Wragg *et al*. 2022), which resulted, after filters were applied, in a total sample set of 750 animals. The high-density BovineHD 777K SNP data set was then downsampled to the subset of SNPs in common with the Illumina^®^ Bovine SNP50 BeadChip.

Low-density genome-wide SNP array data sets (Illumina^®^ Bovine SNP50 BeadChip) were also obtained for 39 cattle that were part of a similar trypanosomiasis time course infection study using RNA-seq data (Berthier *et al*. 2015; Peylhard *et al*. 2023). Several of the populations examined in this study were not included in the initial data set used for the previous local ancestry analysis (McHugo *et al*. 2025a); therefore, additional low-density genome-wide SNP array data sets (Illumina^®^ Bovine SNP50 BeadChip) were obtained from published studies via the Web-Interfaced Next Generation Database (WIDDE) resource for cattle and sheep SNP array genotype data (Gautier *et al*. 2009; Decker *et al*. 2014; Flori *et al*. 2014; Sempere *et al*. 2015). In total, 26 different populations were represented (**Table 1**): three European *B. taurus* populations (Holstein-Friesian, Angus, and Jersey); four African *B. taurus* populations (Muturu, Lagune, Guinean N’Dama, and Baoulé); three *B. indicus* populations (Tharparkar, Gir, and Nelore); five residually admixed European populations (Romagnola, Chianina, Marchigiana, Maremmana, and Alentejana); five trypanotolerant African hybrid populations (hybrid N’Dama, Borgou, Somba, Keteku, and Sheko); and six trypanosusceptible African hybrid populations (Fulani Zebu, Ankole, Nganda, East African Shorthorn Zebu, Karamojong, and Boran).

**Table 1.**
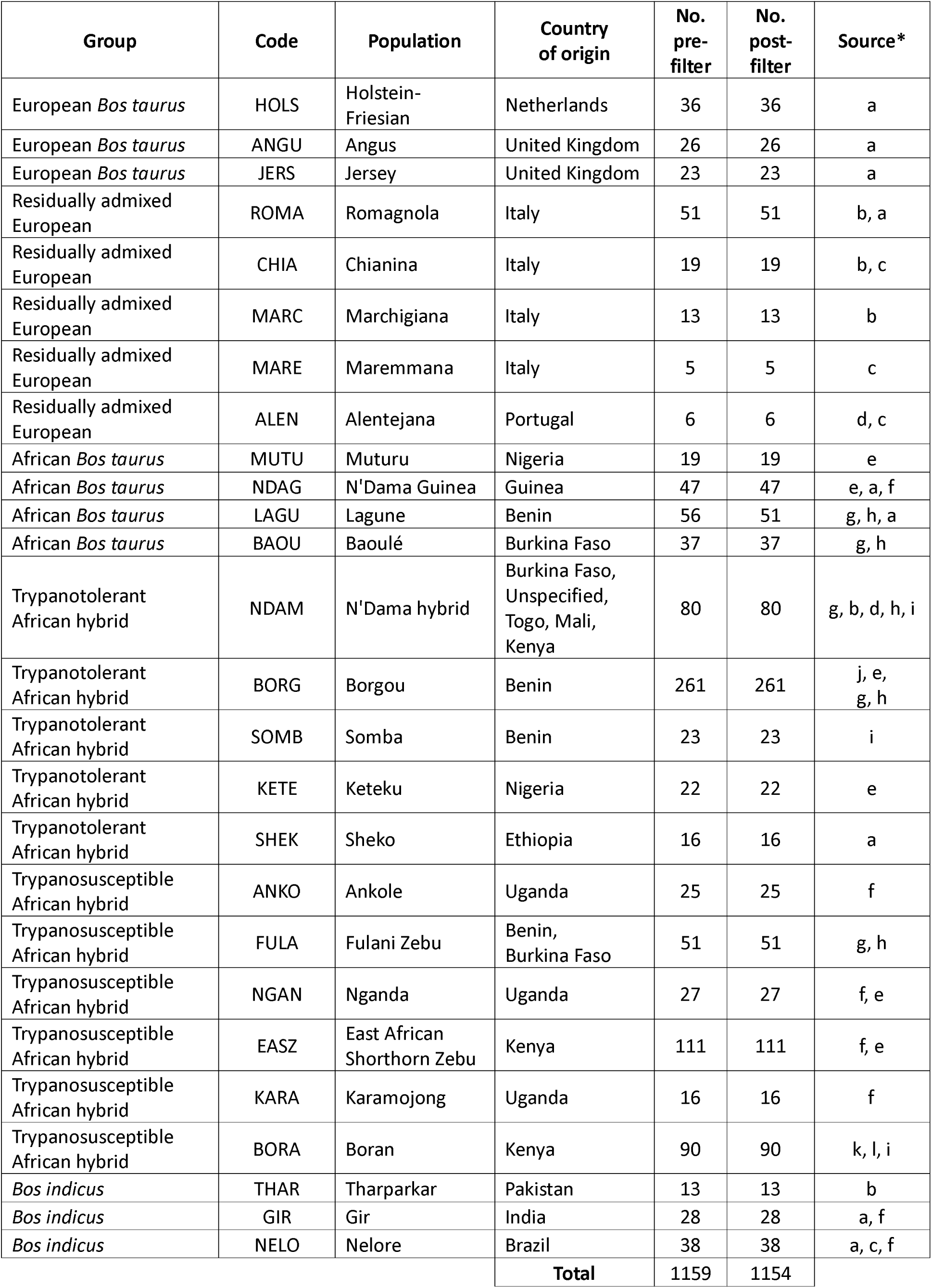
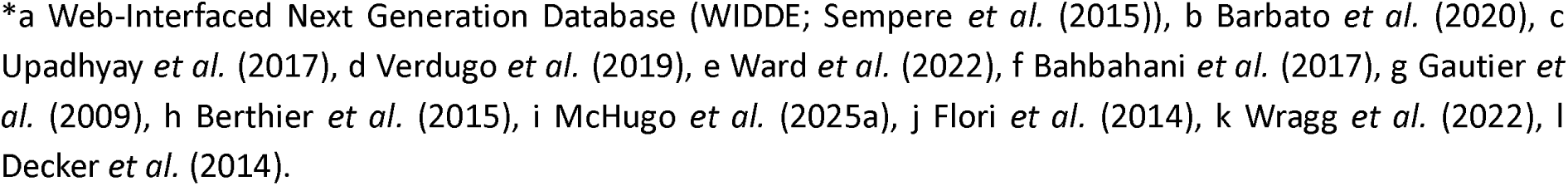
Group, population code, population name, country of origin, number of samples pre- and post-filtering, and sources of SNP data.

| Group | Code | Population | Country of origin | No. pre-filter | No. post-filter | Source* |
| --- | --- | --- | --- | --- | --- | --- |
| European *Bos taurus* | HOLS | Holstein-Friesian | Netherlands | 36 | 36 | a |
| European *Bos taurus* | ANGU | Angus | United Kingdom | 26 | 26 | a |
| European *Bos taurus* | JERS | Jersey | United Kingdom | 23 | 23 | a |
| Residually admixed European | ROMA | Romagnola | Italy | 51 | 51 | b, a |
| Residually admixed European | CHIA | Chianina | Italy | 19 | 19 | b, c |
| Residually admixed European | MARC | Marchigiana | Italy | 13 | 13 | b |
| Residually admixed European | MARE | Maremmiana | Italy | 5 | 5 | c |
| Residually admixed European | ALEN | Alentejana | Portugal | 6 | 6 | d, c |
| African *Bos taurus* | MUTU | Muturu | Nigeria | 19 | 19 | e |
| African *Bos taurus* | NDAG | N'Dama Guinea | Guinea | 47 | 47 | e, a, f |
| African *Bos taurus* | LAGU | Lagune | Benin | 56 | 51 | g, h, a |
| African *Bos taurus* | BAOU | Baoulé | Burkina Faso | 37 | 37 | g, h |
| Trypanotolerant African hybrid | NDAM | N'Dama hybrid | Burkina Faso, Unspecified, Togo, Mali, Kenya | 80 | 80 | g, b, d, h, i |
| Trypanotolerant African hybrid | BORG | Borgou | Benin | 261 | 261 | j, e, g, h |
| Trypanotolerant African hybrid | SOMB | Somba | Benin | 23 | 23 | i |
| Trypanotolerant African hybrid | KETE | Keteku | Nigeria | 22 | 22 | e |
| Trypanotolerant African hybrid | SHEK | Sheko | Ethiopia | 16 | 16 | a |
| Trypanosusceptible African hybrid | ANKO | Ankole | Uganda | 25 | 25 | f |
| Trypanosusceptible African hybrid | FULA | Fulani Zebu | Benin, Burkina Faso | 51 | 51 | g, h |
| Trypanosusceptible African hybrid | NGAN | Nganda | Uganda | 27 | 27 | f, e |
| Trypanosusceptible African hybrid | EASZ | East African Shorthorn Zebu | Kenya | 111 | 111 | f, e |
| Trypanosusceptible African hybrid | KARA | Karamojong | Uganda | 16 | 16 | f |
| Trypanosusceptible African hybrid | BORA | Boran | Kenya | 90 | 90 | k, l, i |
| *Bos indicus* | THAR | Tharparkar | Pakistan | 13 | 13 | b |
| *Bos indicus* | GIR | Gir | India | 28 | 28 | a, f |
| *Bos indicus* | NELO | Nelore | Brazil | 38 | 38 | a, c, f |
| **Total** |  |  |  | 1159 | 1154 |  |

\*a Web-Interfaced Next Generation Database (WIDDE; Sempere *et al.* (2015)), b Barbato *et al.* (2020), c Upadhyay *et al.* (2017), d Verdugo *et al.* (2019), e Ward *et al.* (2022), f Bahbahani *et al.* (2017), g Gautier *et* *al.* (2009), h Berthier *et al.* (2015), i McHugo *et al.* (2025a), j Flori *et al.* (2014), k Wragg *et al.* (2022), l Decker *et al.* (2014).

N’Dama cattle are the most well studied trypanotolerant population and control parasitaemia and anaemia significantly better than trypanosusceptible cattle and have increased trypanotolerance compared to hybrid populations (Murray *et al*. 1984; Paling *et al*. 1991; Achukwi *et al*. 1997; O’Gorman *et al*. 2006; Meade *et al*. 2009; O’Gorman *et al*. 2009; Noyes *et al*. 2011; Berthier *et al*. 2015). Lagune and Baoulé cattle were found to exhibit similar levels of trypanotolerance to the N’Dama population (Berthier *et al*. 2015). The endangered Muturu population is not as well characterised as other trypanotolerant populations; however, minimal gene flow was identified between the N’Dama and Muturu populations, illustrating their genetic separation and the importance of their conservation as a trypanotolerant population (Ibeagha-Awemu & Erhardt 2005; Tijjani *et al*. 2019). While the trypanotolerant African hybrid populations analysed in this study are classed as trypanotolerant, their response to trypanosome infection is intermediate to that of the trypanotolerant populations with higher levels of African *B. taurus* ancestry and trypanosusceptible populations (Berthier *et al*. 2015; Food and Agriculture Organization 2023). This means that they exhibit a lower ability to control both parasitaemia and anaemia during trypanosome infection than other trypanotolerant populations, likely because they are hybrid populations and the degree of trypanotolerance has been show to correlate with levels of African *B. taurus* ancestry (Berthier *et al*. 2015). The trypanosusceptible African hybrid populations analysed in this study show no evidence of trypanotolerance despite their long history of exposure to trypanosome parasites (O’Gorman *et al*. 2006; Meade *et al*. 2009; O’Gorman *et al*. 2009; Noyes *et al*. 2011; Berthier *et al*. 2015; Food and Agriculture Organization 2023). Similarly, the *B. indicus* and European *B. taurus* populations have no documented trypanotolerance despite their introduction into trypanosome infested areas (Berthier *et al*. 2015; Food and Agriculture Organization 2023). Finally, the residually admixed European populations also have no documented evidence of trypanotolerance and, although trypanosomiasis in cattle is rare in Europe, the low level of African *B. taurus* ancestry in the breeds means that trypanotolerance is unlikely as the trait is known to correlate with increased African *B. taurus* ancestry (Berthier *et al*. 2015; Barbato *et al*. 2020; Food and Agriculture Organization 2023).

#### Gene expression data

As previously described (McHugo *et al*. 2025b), Affymetrix^®^ GeneChip^®^ Bovine Genome Array data sets were obtained from published studies with parallel trypanosome infection time course experimental designs (O’Gorman *et al*. 2006; Meade *et al*. 2009; O’Gorman *et al*. 2009; Noyes *et al*. 2011) in which trypanotolerant African hybrid N’Dama and trypanosusceptible African hybrid Boran cattle were experimentally infected with the *Trypanosoma congolense* clone IL1180 (Geigy & Kauffmann 1973; Nantulya *et al*. 1984) delivered via the bites of infected tsetse flies (*Glossina morsitans morsitans*) (Akol & Murray 1982; Dwinger *et al*. 1987). Samples were collected before infection and at various days post-infection (dpi) and included peripheral blood mononuclear cells (PBMC) that were isolated from blood (BL), liver (LI), lymph node (LN), and spleen (SP) samples. This resulted in a total of 220 samples from 50 animals (25 trypanotolerant N’Dama and 25 trypanosusceptible Boran) collected across 12 time points and four tissues before filtering (**Table 2**). **Fig 1** illustrates the experimental design and study workflow. The computer code required to repeat and reproduce the analyses is available at doi.org/10.5281/zenodo.11517978.

**Fig 1.**
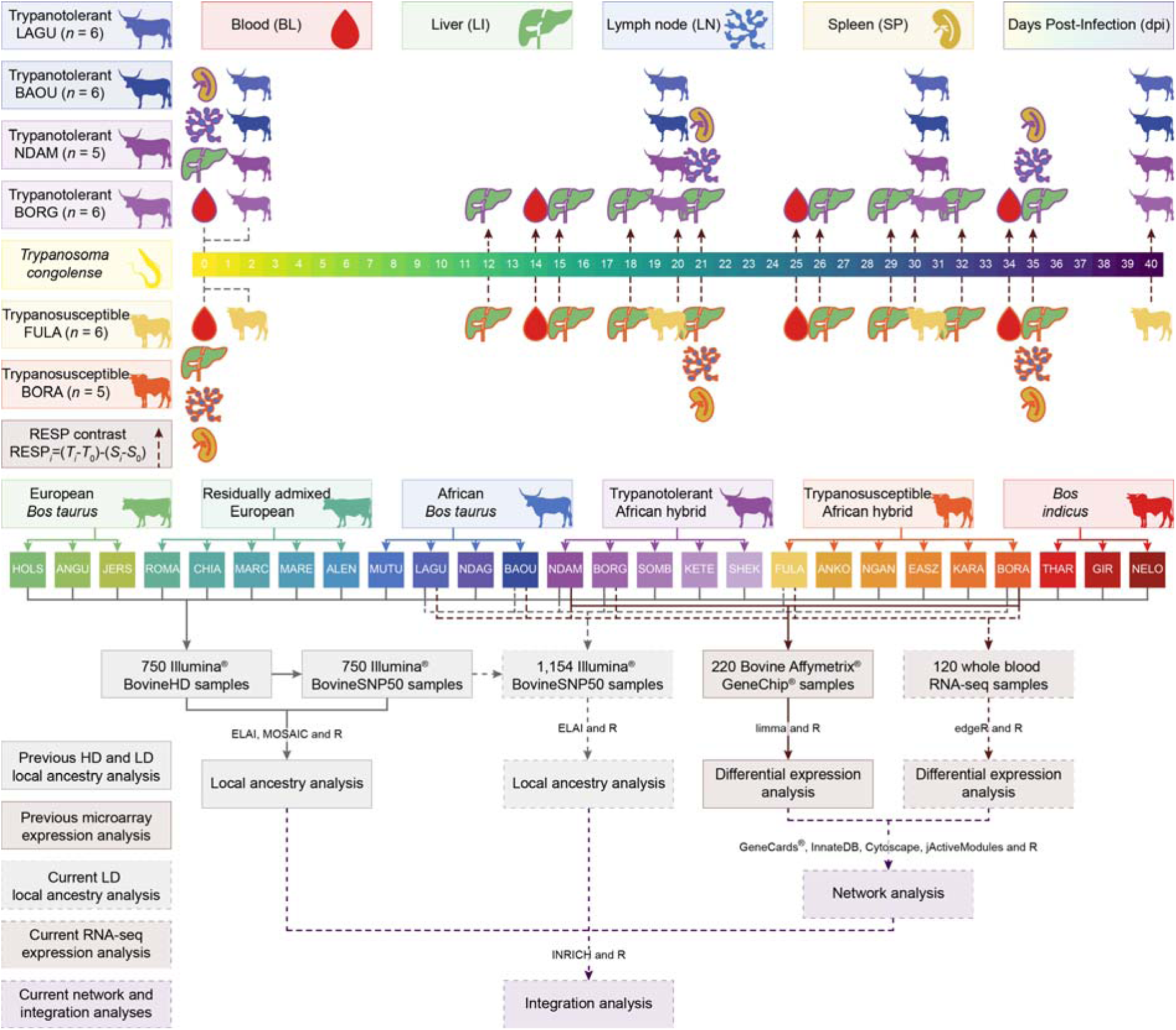
Diagram showing the experimental design and study workflow. Trypanosome image by Matus Valach and cattle images by Tracy A. Heath, T. Michael Keesey, and Steven Traver via <u>phylopic.org</u> and tissue images via <u>healthicons.org</u>. Colours from the khroma (v. 1.10.0) (Frerebeau 2023) and viridis (v. 0.6.3) (Garnier et al. 2023) R packages.

**Table 2.** Days post-infection (dpi), population code, tissue, data type, number of samples pre- and post-filtering, and sources of gene expression data.

| dpi | Population | Tissue | Data type | No. pre-filtering | No. post-filtering | Source* |
| --- | --- | --- | --- | --- | --- | --- |
| 0 | NDAM | BL | Microarray | 5 | 5 | a |
| 0 | BORA | BL | Microarray | 5 | 5 | a |
| 0 | NDAM | LI | Microarray | 20 | 19 | b |
| 0 | BORA | LI | Microarray | 20 | 18 | b |
| 0 | NDAM | LN | Microarray | 5 | 5 | b |
| 0 | BORA | LN | Microarray | 5 | 4 | b |
| 0 | NDAM | SP | Microarray | 5 | 5 | b |
| 0 | BORA | SP | Microarray | 5 | 4 | b |
| 0 | LAGU | WB | RNA-seq | 6 | 6 | c |
| 0 | BAOU | WB | RNA-seq | 6 | 5 | c |
| 0 | NDAM | WB | RNA-seq | 6 | 6 | c |
| 0 | BORG | WB | RNA-seq | 6 | 5 | c |
| 0 | FULA | WB | RNA-seq | 6 | 5 | c |
| 12 | NDAM | LI | Microarray | 5 | 5 | b |
| 12 | BORA | LI | Microarray | 5 | 5 | b |
| 14 | NDAM | BL | Microarray | 5 | 5 | a |
| 14 | BORA | BL | Microarray | 5 | 5 | a |
| 15 | NDAM | LI | Microarray | 5 | 5 | b |
| 15 | BORA | LI | Microarray | 5 | 5 | b |
| 18 | NDAM | LI | Microarray | 5 | 5 | b |
| 18 | BORA | LI | Microarray | 5 | 5 | b |
| 20 | LAGU | WB | RNA-seq | 6 | 6 | c |
| 20 | BAOU | WB | RNA-seq | 6 | 5 | c |
| 20 | NDAM | WB | RNA-seq | 6 | 6 | c |
| 20 | BORG | WB | RNA-seq | 6 | 5 | c |
| 20 | FULA | WB | RNA-seq | 6 | 5 | c |
| 21 | NDAM | LI | Microarray | 5 | 5 | b |
| 21 | BORA | LI | Microarray | 5 | 5 | b |
| 21 | NDAM | LN | Microarray | 5 | 4 | b |
| 21 | BORA | LN | Microarray | 5 | 5 | b |
| 21 | NDAM | SP | Microarray | 5 | 5 | b |
| 21 | BORA | SP | Microarray | 5 | 5 | b |
| 25 | NDAM | BL | Microarray | 5 | 5 | a |
| 25 | BORA | BL | Microarray | 5 | 5 | a |
| 26 | NDAM | LI | Microarray | 5 | 3 | b |
| 26 | BORA | LI | Microarray | 5 | 5 | b |
| 29 | NDAM | LI | Microarray | 5 | 5 | b |
| 29 | BORA | LI | Microarray | 5 | 5 | b |
| 30 | LAGU | WB | RNA-seq | 6 | 6 | c |
| 30 | BAOU | WB | RNA-seq | 6 | 5 | c |
| 30 | NDAM | WB | RNA-seq | 6 | 6 | c |
| 30 | BORG | WB | RNA-seq | 6 | 5 | c |
| 30 | FULA | WB | RNA-seq | 6 | 5 | c |
| 32 | NDAM | LI | Microarray | 5 | 5 | b |
| 32 | BORA | LI | Microarray | 5 | 5 | b |
| 34 | NDAM | BL | Microarray | 5 | 5 | a |
| 34 | BORA | BL | Microarray | 5 | 5 | a |
| 35 | NDAM | LI | Microarray | 5 | 5 | b |
| 35 | BORA | LI | Microarray | 5 | 5 | b |
| 35 | NDAM | LN | Microarray | 5 | 4 | b |
| 35 | BORA | LN | Microarray | 5 | 5 | b |
| 35 | NDAM | SP | Microarray | 5 | 5 | b |
| 35 | BORA | SP | Microarray | 5 | 5 | b |
| 40 | LAGU | WB | RNA-seq | 6 | 6 | c |
| 40 | BAOU | WB | RNA-seq | 6 | 5 | c |
| 40 | NDAM | WB | RNA-seq | 6 | 6 | c |
| 40 | BORG | WB | RNA-seq | 6 | 5 | c |
| 40 | FULA | WB | RNA-seq | 6 | 5 | c |
| **Total** |  |  | Microarray | 220 | 211 |  |
|  |  |  | RNA-seq | 120 | 108 |  |

\*a McHugo *et al.* (2025b), b Noyes *et al.* (2011), c Peylhard *et al.* (2023).

A total of 120 RNA-seq data sets were obtained from a similar trypanosome infection time course experiment in which 30 animals (six trypanotolerant African *B. taurus* Lagune, six trypanotolerant African *B. taurus* Baoulé, six trypanotolerant African hybrid N’Dama, six trypanotolerant African hybrid Borgou, and six trypanosusceptible African hybrid Fulani Zebu), were infected via intravenous inoculation with 10^5^ trypanosomes of the same *T. congolense* IL1180 strain (Geigy & Kauffmann 1973; Nantulya *et al*. 1984; Berthier *et al*. 2015; Peylhard *et al*. 2023). Whole blood (WB) samples were taken before infection and at 20, 30, and 40 dpi and total RNA was extracted for sequencing (Berthier *et al*. 2015; Peylhard *et al*. 2023).

### Analysis of genomic data

#### Preparation of genomic data

The additional SNP samples were converted to binary PLINK files with Illumina^®^ allele coding for the FORWARD strand as required using PLINK (v. 1.90 beta 6.25) (Chang *et al*. 2015) and SNPchiMp (v. 3) (Nicolazzi *et al*. 2015). The Illumina^®^ Bovine SNP50 BeadChip SNP locations were updated from UMD3.1 to the newer bovine genome assembly ARS-UCD1.2 (Rosen *et al*. 2020) using coordinates from the NAGRP Data Repository (Schnabel 2018) and PLINK. The additional SNP samples were downsampled to the subset of the 46,713 SNPs in common with the BovineHD 777K BeadChip using PLINK with a list of the SNPs from the previously downsampled low-density SNP data set generated using dplyr (v. 1.1.2) (Wickham *et al*. 2023a), and readr (v. 2.1.4) (Wickham *et al*. 2023b) with R (v. 4.3.2) (R Core Team 2023). The samples were then merged with the previously downsampled low-density SNP data set with PLINK. The merged low-density SNP data was filtered as described in our previous study (McHugo *et al*. 2025a). Briefly, this included the removal of samples that had missing call rates exceeding 0.95 in addition to samples duplicated in multiple data sources that had an identity-by-state value ≥ 0.99 with PLINK. The data set was then filtered to retain autosomal SNPs with a minimum call rate of 95% and minor allele frequency (MAF) of at least 5% with PLINK.

#### Population genomic analyses

The results of the population genomic analyses of high- and low-density SNP data sets were obtained from our previous study and the same analyses of the merged low-density SNP data set were performed as described in that study (McHugo *et al*. 2025a). Briefly, an inbreeding analysis was performed with PLINK and principal component analysis (PCA) was performed using smartpca after file conversion with convertf, both part of the EIGENSOFT package (v. 7.1.2) (Patterson *et al*. 2006). Genetic structure analysis was performed using structure_threader (v. 1.3.4) (Pina-Martins *et al*. 2017) with fastStructure (v. 1.0) (Raj *et al*. 2014) with the model complexity or number of populations (*K*) set from 2 to 27. The chooseK function was used to test the outputs to find a range of values of *K* that best accounted for the structure in the data (Raj *et al*. 2014).

#### Local ancestry analysis and functional enrichment of introgressed regions

After conversion of the binary PLINK files into BIMBAM format with PLINK, local ancestry inference (LAI) was performed separately for each bovine autosome in the populations with additional low-density SNP samples added to the previously analysed data set (McHugo *et al*. 2025a) using 30 expectation-maximization (EM) steps, three upper clusters, 15 lower clusters, and 200 mixing generations using the Efficient Local Ancestry Inference (ELAI) software (v. 1.0) (Guan 2014). Based on the previous local ancestry results (McHugo *et al*. 2025a), the donor populations for the LAI were the Angus (European *B. taurus*), Guinean N’Dama (African *B. taurus*), and Gir (*B. indicus*). Mean ancestry scores across the individual hybrid animals and a genome-wide *z*-score for each of the three ancestry components were estimated for each hybrid population.

Functional enrichment of the introgressed regions was performed using gprofiler2 (v. 0.2.2) (Kolberg *et al*. 2023) with R. The background set was the set of genes within genomic intervals of 1 Mb up- and downstream from a SNP in the data set. The query sets were the genes within 1 Mb up and downstream from the SNPs with a *z*-score ≥ 2.0 for each of the ancestries.

### Analysis of gene expression data

#### Differential expression analysis

The results of differential expression analysis of the microarray data were obtained from our previous study (McHugo *et al*. 2025b). Differential expression analysis of the RNA-seq data was performed using edgeR (v. 4.0.16) (Robinson *et al*. 2010) with R. The scripts published with the intermediate tables and raw data from Peylhard and colleagues (Peylhard *et al*. 2023) were modified to analyse response (RESP) contrasts to identify changes in expression over time in the trypanotolerant samples (LAGU, BAOU, NDAM, and BORG) relative to the trypanosusceptible FULA (RESP*_i_* = [*T_i_* – *T*_0_] – [*S_i_* – *S*_0_], where *T* represents the trypanotolerant populations and *S* represents the trypanosusceptible populations) (**Fig 1**) (Noyes *et al*. 2011). This was done to facilitate comparison with the previous analysis of the microarray data (McHugo *et al*. 2025b) and because the original analysis of the RNA-seq data did not include these contrasts (Peylhard *et al*. 2023). As in the analysis conducted by Peylhard and colleagues (Peylhard *et al*. 2023), generalized linear model likelihood ratio tests were used to calculate log_2_ fold change values and Benjamini-Hochberg (B-H) corrected *P*-values (Benjamini & Hochberg 1995; McCarthy *et al*. 2012). Genes with an adjusted *P*-value of ≤ 0.05 (B-H *P*_adj._ ≤ 0.05) were considered to be significantly differentially expressed.

#### Gene interaction network analysis

The GeneCards^®^ database (genecards.org, v. 5.19) (Stelzer *et al*. 2016), which integrates data from almost 200 different biological databases, was searched for genes relating to the search term “trypano*”. The results were exported, filtered to select genes with a relevance score ≥ 1.75 and converted from gene symbols to Ensembl IDs to prepare a computationally manageable amount of genes based on the methodology described by Hall and colleagues (Hall *et al*. 2021) and using dplyr, gprofiler2, readr, and stringr (v. 1.5.0) (Wickham 2023) with R. The resulting file was then used as an input for network analysis to identify interactions among these genes and other genes in the bovine interactome using InnateDB (<u>innatedb.com</u>, v. 5.4) (Breuer *et al*. 2013) with interactions predicted by orthology included. The node IDs of the resulting network were prepared using dplyr, readr, and stringr with R. The network was then imported into Cytoscape (v. 3.8.0) (Shannon *et al*. 2003). The differential gene expression results obtained using the microarray data (McHugo *et al*. 2025b) and the RNA-seq data described above were imported as node tables.

Expression-activated subnetworks or modules within the base network were identified using jActiveModules (v. 3.2.1) (Ideker *et al*. 2002) for the B-H corrected *P*-values for each of the eight final contrasts in the microarray and RNA-seq data sets as the final time point produced the highest number of significantly differentially expressed genes (DEGs) in both data sets. The highest scoring module for each contrast was selected, analysed with Cytoscape and yFiles Layout Algorithms (v. 1.1.3) (Becker & Rojas 2001) was used to remove overlaps. The modules were outputted as network files and images and analysed with R.

#### Functional enrichment analysis of gene expression data

Functional enrichment was performed for the complete set of significant DEGs identified by analysis of the RNA-seq data for the response contrasts and the modules identified for both the microarray and RNA-seq data using gprofiler2 with R. The background set for the significant DEGs was the set of detectable genes identified by Peylhard and colleagues (Peylhard *et al*. 2023), while the background set for the modules was the base network. The analyses were restricted to gene ontology (GO) terms and the driver GO terms were highlighted.

### Integration of genomic local ancestry inference results and gene expression data

The results of the LAI (McHugo *et al*. 2025a) and expression analyses (McHugo *et al*. 2025b), including the new results described in this study, were integrated with INRICH (INterval enRICHment analysis, v. 1.1) (Lee *et al*. 2012). A reference gene file of all bovine genes was prepared using biomaRt (v. 2.58.2) (Durinck *et al*. 2009) and readr with R. Reference SNP files were prepared from the bim files for the associated data sets using dplyr and readr with R. A target gene set file containing the genes in the first module identified for each of the eight final contrasts in the microarray and RNA-seq data sets converted to Ensembl IDs was prepared using dplyr, gprofiler2, readr, stringr, and tidyr (v. 1.3.0) (Wickham *et al*. 2023c) with R. Associated genomic interval files were prepared to include regions 1 Mb up- and downstream from SNPs with a *z*-score ≥ 2.0 for each ancestry component of the LAI results (McHugo *et al*. 2025a), excluding the results from the low-density SNP data set analysed with MOSAIC due to the low resolution obtained. This procedure involved preparing files with *z*-scores converted to *P*-values using dplyr, readr, and stringr with R. These data were used as input for linkage disequilibrium (LD) clumping of SNPs 1 Mb up- and downstream from those with converted *P*-values ≤ 0.02275 (equivalent to *z*-scores ≥ 2.0) using the associated SNP data files with PLINK. The resulting clumped files were then converted to interval files for use with INRICH with the intervals extending 1 Mb up- and downstream from the central SNP in each clump using dplyr, readr, and stringr with R. Interval enrichment analyses were then performed for the target gene set file containing the modules and the interval files from the LAI results with the reference gene file and appropriate reference SNP files using INRICH with a target size filter of 2500 to include all the modules and specifying non-human data.

The results of the analyses were visualised using ComplexUpset (v. 1.3.3) (Krassowski 2020), dplyr, ggh4x (v. 0.2.4) (van den Brand 2023), ggplot2 (v. 3.4.2) (Wickham 2009), ggrepel (v. 0.9.3) (Slowikowski 2023), ggtext (v. 0.1.2) (Wilke & Wiernik 2022), magick (v. 2.8.1) (Ooms 2023) with ImageMagick (v. 6.9.12.96) (ImageMagick Studio LLC 2023), magrittr (v. 2.0.3) (Bache & Wickham 2022), parallel (v. 4.3.2) (R Core Team 2023), patchwork (v. 1.1.2) (Pedersen 2023), purrr (v. 1.0.1) (Wickham & Henry 2023), readr, rlang (v. 1.1.1) (Henry & Wickham 2024), scales (v. 1.2.1) (Wickham *et al*. 2023d), stringr, tibble (v. 3.2.1) (Müller & Wickham 2023), tidyr with R. Colours were generated from khroma (v. 1.10.0) (Frerebeau 2023) and viridis (v. 0.6.3) (Garnier *et al*. 2023).

## Results

### Population genomics results reiterate known evolutionary relationships

#### Post-filtering SNP data

After filtering for missing genotypes and identity-by-state (**Fig S1**) there were 1154 animals in the merged low-density data set (**Table 1**). The inbreeding results did not indicate additional filtering was required (**Fig S2**). Filtering for autosomal SNPs with a minimum call rate of 95% and MAF of at least 5% retained 29,869 SNPs with a total genotyping rate of 99.47%.

#### Principal component analysis recapitulates the biogeography of domestic cattle

The first principal component (PC1) explained 43.82% of the total variation for PC1–10 in the data for the SNP data and separated the *B. taurus* and *B. indicus* lineages (**Fig 2**, **S3**). The second principal component (PC2) explained a further 27.69% of the total variation for PC1–10 and separated the European *B. taurus* and African *B. taurus* groups (**Fig 2**). The hybrid and residually admixed animals emerged between the reference populations with the residually admixed European animals clustering close to the European *B. taurus* populations and the African hybrid animals mostly dispersed between the African *B. taurus* and *B. indicus* populations (**Fig 2**). The trypanotolerant African hybrid animals are closest to the African *B. taurus* populations, while the trypanosusceptible African hybrid animals are closest to the *B. indicus* populations (**Fig 2**).

**Fig 2.**
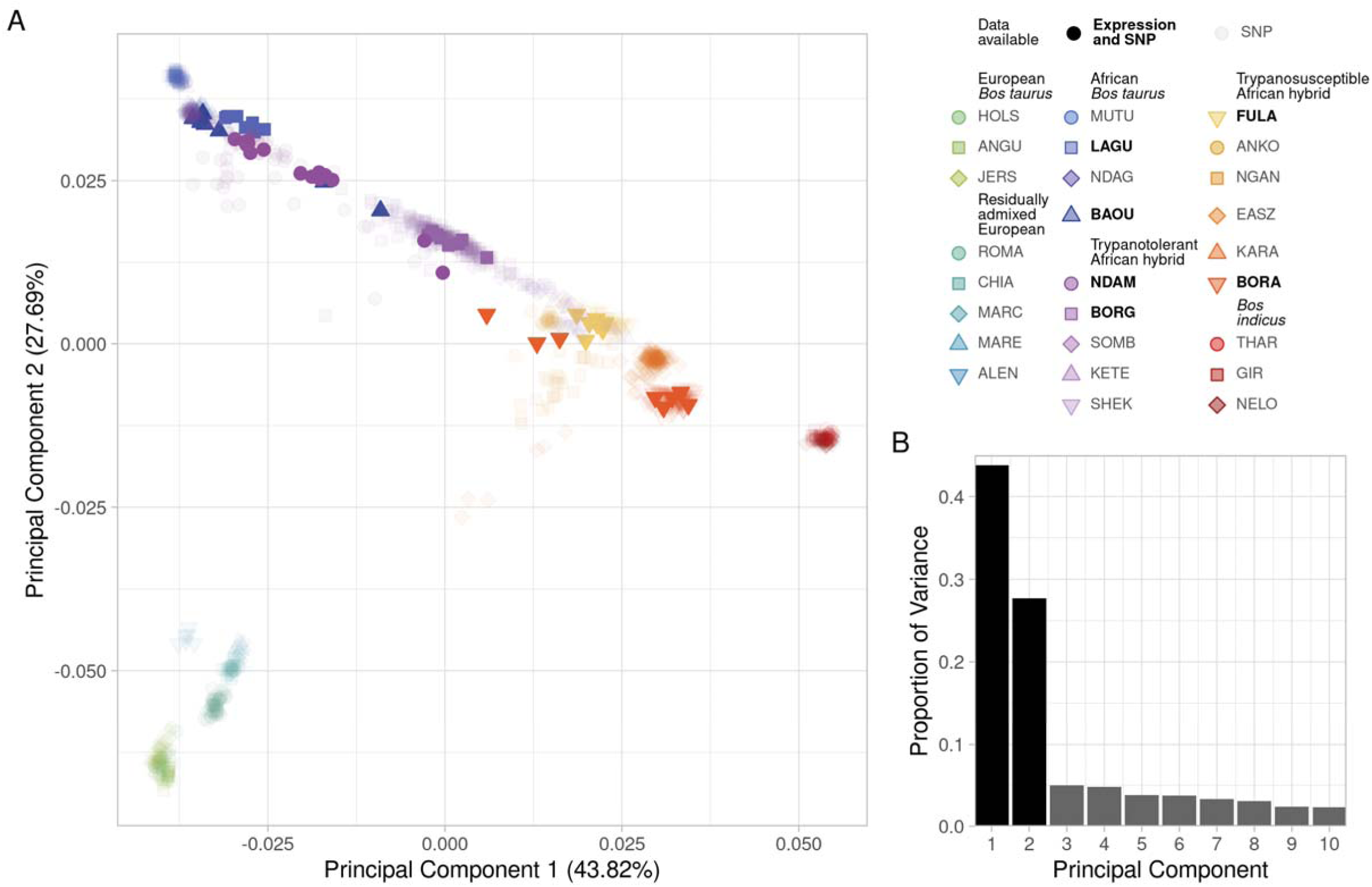
**A.** Principal component analysis of low-density SNP data for the cattle samples coloured according to population showing the first two principa components and **B.** bar chart of proportion of variance of the top ten principal components. The transparency indicates the availability of gene expression data for the sample

#### Genetic structure analysis highlights the multiple ancestries of African cattle

The genetic structure results for the number of assumed populations (*K*) set to three partitioned the European and African *B. taurus* and *B. indicus* ancestry in the data set (**Fig S6–S9**). The model complexity that maximizes the marginal likelihood was 17 and the model components used to explain the structure in the data was 21.

#### Local ancestry inference revealed peaks across admixed African cattle populations

Mean LAI values were estimated for each of the three ancestry components (European *B. taurus*, African *B. taurus*, and *B. indicus*) for the six populations with gene expression data available (LAGU, BAOU, NDAM, BORG, FULA, and BORA) (**Fig 3**). Genome-wide *z*-scores for the mean LAI results were used to identify SNPs with *z*-scores ≥ 2.0 for each population (**Table S2**). There were no SNPs that passed the *z* ≥ 2.0 threshold for the African *B. taurus* ancestry component in the African *B. taurus* populations (LAGU and BAOU), while the trypanotolerant (NDAM and BORG) and trypanosusceptible (FULA and BORA) African hybrid populations had the lowest number of SNPs passing the *z* ≥ 2.0 threshold for the African *B. taurus* and *B. indicus* ancestry components, respectively (**Table S2**). The European *B. taurus* ancestry components had the highest number of SNPs passing the *z* ≥ 2.0 threshold in all six populations (**Table S2**)

**Fig 3.**
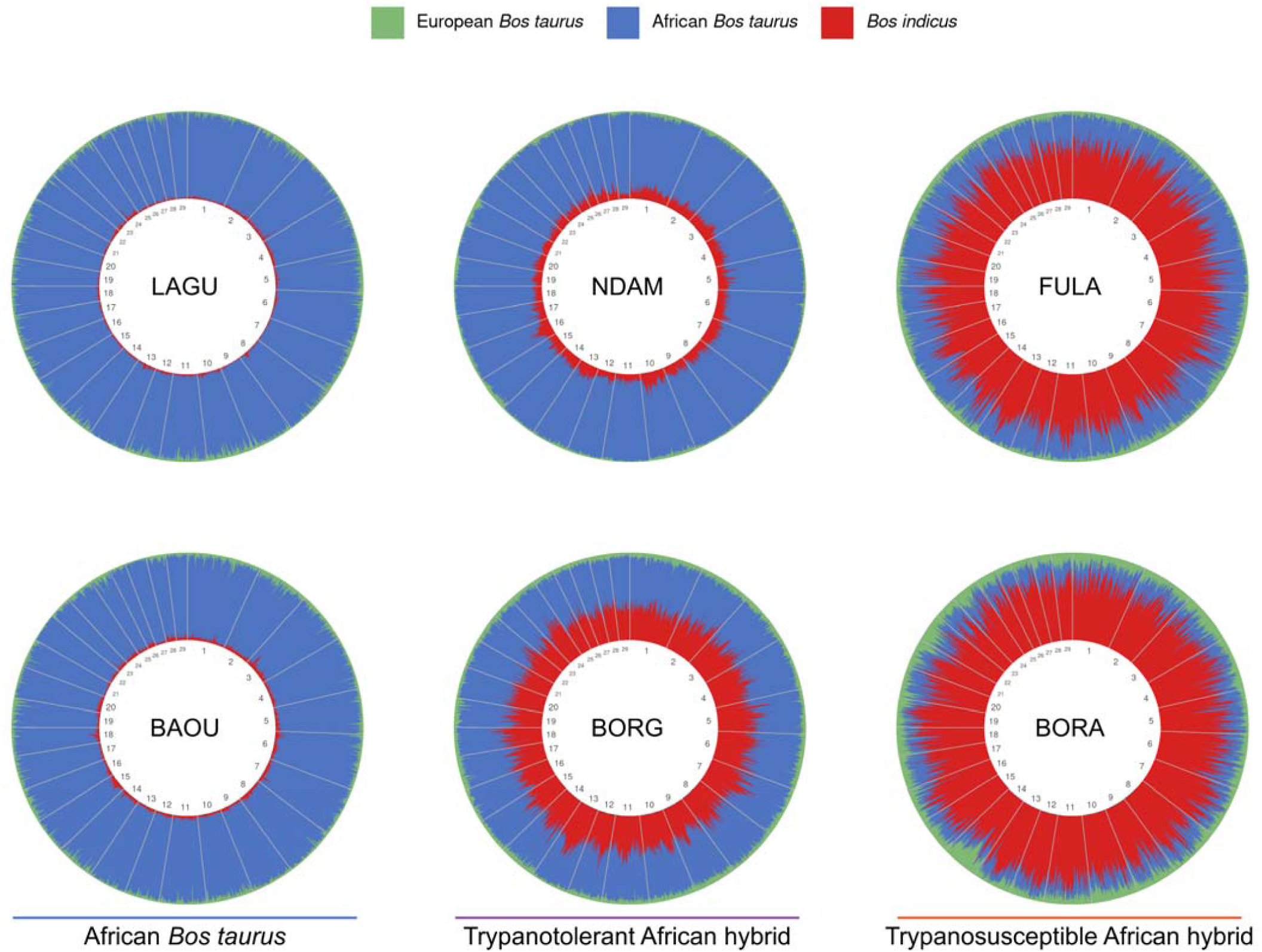
Local ancestry plots showing mean European B. taurus, African B. taurus, and B. indicus ancestry components using low-density SNP data for the six populations with gene expression data available across all autosomes. Each vertical line on the round genome plots represents a SNP and is coloured according to the ancestry results.

#### Introgressed genomic regions showed enrichment of biologically relevant functional categories

The proportions of the numbers of genes found within 1 Mb up- and downstream from each SNP with a *z*-score ≥ 2.0 are similar to those of the numbers of SNPs found for each ancestry component in the populations studied (**Table S1**, **S2**). The top driver GO terms enriched for the European *B. taurus* genes in the African *B. taurus* populations that had gene expression data available (LAGU, BAOU) included those relating to alcohol dehydrogenase activity (*GO:0006069 ethanol oxidation* and *GO:0004024 alcohol dehydrogenase activity, zinc-dependent*) and aspartic-type endopeptidase activity (*GO:0004190 aspartic-type endopeptidase activity*). The top driver GO terms enriched for the *B. indicus* genes include those related to catabolic processes and catalytic activity (*GO:0030574 collagen catabolic process*, *GO:0006032 chitin catabolic process*, and *GO:0004190 aspartic-type endopeptidase activity*); cell signalling (*GO:0051606 detection of stimulus*, *GO:0007186 protein-coupled receptor signalling pathway*, and *GO:0004930 G protein-coupled receptor activity*); haemoglobin complex (*GO:0005833 haemoglobin complex*); and olfactory receptor activity (*GO:0004984 olfactory receptor activity*) (**Fig S10**). As there were no African *B. taurus* SNPs that passed the *z* ≥ 2 threshold there were no African *B. taurus* genes for functional enrichment in these populations (**Table S1**, **S2**).

The trypanotolerant African hybrid populations that had gene expression data available (NDAM and BORG) also had top driver GO terms enriched for the European *B. taurus* genes which related to alcohol dehydrogenase activity (*GO:0006069 ethanol oxidation* and *GO:0004024 alcohol dehydrogenase activity, zinc-dependent*) as well as chemotaxis (*GO:0050918 positive chemotaxis*, *GO:0031731 CCR6 chemokine receptor binding*, and *GO:0042056 chemoattractant activity*); spliceosomal complex assembly (*GO:0000348 mRNA branch site recognition*, *GO:0005686 U2 snRNP*, and *GO:0045131 pre-mRNA branch point binding*); cellular component organisation (*GO:0043933 protein-containing complex organisation* and *GO:0043229 intracellular organelle*); and collagen catabolic process (*GO:0030574 collagen catabolic process*) (**Fig S11**).

The top driver GO terms enriched for the *B. indicus* genes include those related to the skin and keratin (*GO:0031424 keratinisation, GO:0045109 intermediate filament organisation*, *GO:0045095 keratin filament*, *GO:0030280 structural constituent of skin epidermis*, and *GO:0045103 intermediate filament-based process*); L-amino acid transmembrane transport (*GO:0097638 L-arginine import across plasma membrane*, *GO:1903352 L-ornithine transmembrane transport*, and *GO:0000064 L-ornithine transmembrane transporter activity*); oxidoreductase activity (*GO:0004499 N,N-dimethylaniline monooxygenase activity*, *GO:0047822 hypotaurine dehydrogenase activity*, *GO:0047023 androsterone dehydrogenase activity*, and *GO:0047086 ketosteroid monooxygenase activity*); and aminoglycoside antibiotic metabolic process (*GO:0030647 aminoglycoside antibiotic metabolic process*) (**Fig S11**). As with the African *B. taurus* populations, there were too few African *B. taurus* genes for these populations for functional enrichment (**Table S1**, **S2**).

The trypanosusceptible African hybrid populations with gene expression data (FULA and BORA) had driver GO terms enriched for African *B. taurus* genes that related to the immune system, particularly the major histocompatibility complex (MHC) (*GO:0002396 MHC protein complex assembly*, *GO:0042613 MHC class II protein complex*, *GO:0023023 MHC protein complex binding*, *GO:0050829 defense response to Gram-negative bacterium*, *GO:0007156 homophilic cell adhesion via plasma membrane adhesion molecules*, *GO:0050870 positive regulation of T cell activation*, *GO:0050830 defense response to Gram-positive bacterium*, *GO:0002250 adaptive immune response*, and *GO:0042605 peptide antigen binding*) (**Fig S12**). Other driver GO terms include those related to lysozyme activity (*GO:0003796 lysozyme activity* and *GO:0031902 late endosome membrane*); cell signalling (*GO:0007186 G protein-coupled receptor signalling pathway*); and olfactory receptor activity (*GO:0004984 olfactory receptor activity*) (**Fig S12**). There were no GO terms enriched for European *B. taurus* genes and only one driver GO term enriched for *B. indicus* genes for these populations (*GO:0007156 homophilic cell adhesion via plasma membrane adhesion molecules*) (**Fig S12**).

### Gene expression analysis results and the immunobiology of trypanosomiasis

#### Differentially expressed genes included immune genes

The differential expression analysis of the RNA-seq data found very few significant DEGs with only 45 genes, including 43 unique genes, across all the response contrasts (**Fig S13**, **Table S3**). The majority of these were for the LAGU population at 40 days post-infection which had 25 significantly DEGs (**Fig S13**, **S14**, **Table S3**). The 40 dpi time point was also the only time point that had significant DEGs for all four populations (**Fig S13–S17**, **Table S3**). The duplicated genes among the significant DEGs included *NEIL2*, which had significantly decreased expression in the LAGU, NDAM, and BORG samples at 40 dpi (**Table S3**). In addition, *LZTS3* had significantly decreased expression in the NDAM samples at 30 and 40 dpi while *SLC11A1* had significantly increased expression in the NDAM samples at 20 and 30 dpi (**Table S3**).

#### Network analysis identified functionally relevant modules

The search of the GeneCards^®^ database (Stelzer *et al*. 2016) for genes relating to the term “trypano*” generated a list of 1,036 genes. The application of a filter to select genes with a relevance score ≥ 1.75 left 417 genes with valid bovine Ensembl IDs. The gene interaction network (GIN) generated using InnateDB (Breuer *et al*. 2013) with this list of genes contained 5,666 genes (nodes) and 15,651 interactions (edges) (**Fig S18**). Modules were identified using jActiveModules within this base network for each of the final response contrasts for the microarray and RNA-seq data (**Fig 4**, **S19–S26**). The smallest module identified was for the microarray blood samples at 34 days post-infection (MICRO BL 34) with 474 genes and 1,166 interactions while the largest module identified was for the microarray spleen samples at 35 days post-infection (MICRO SP 35) with 2,446 genes and 6,326 interactions (**Fig 5**). The MICRO SP 35 module also had the highest number of unique genes (**Fig 5**, **S27**). There were eight genes with valid gene symbols that were present in all eight modules, these were *AGO2* (argonaute RISC catalytic component 2), *CBL* (Cbl proto-oncogene), *CNOT1* (CCR4-NOT transcription complex subunit 1), *EDN1* (endothelin 1), *IL1B* (interleukin 1 beta), *NFKB1* (nuclear factor kappa B subunit 1), *RIPK1* (receptor interacting serine/threonine kinase 1), and *TRAF2* (TNF receptor associated factor 2) (**Fig 5**, **Table S4**).

**Fig 4.**
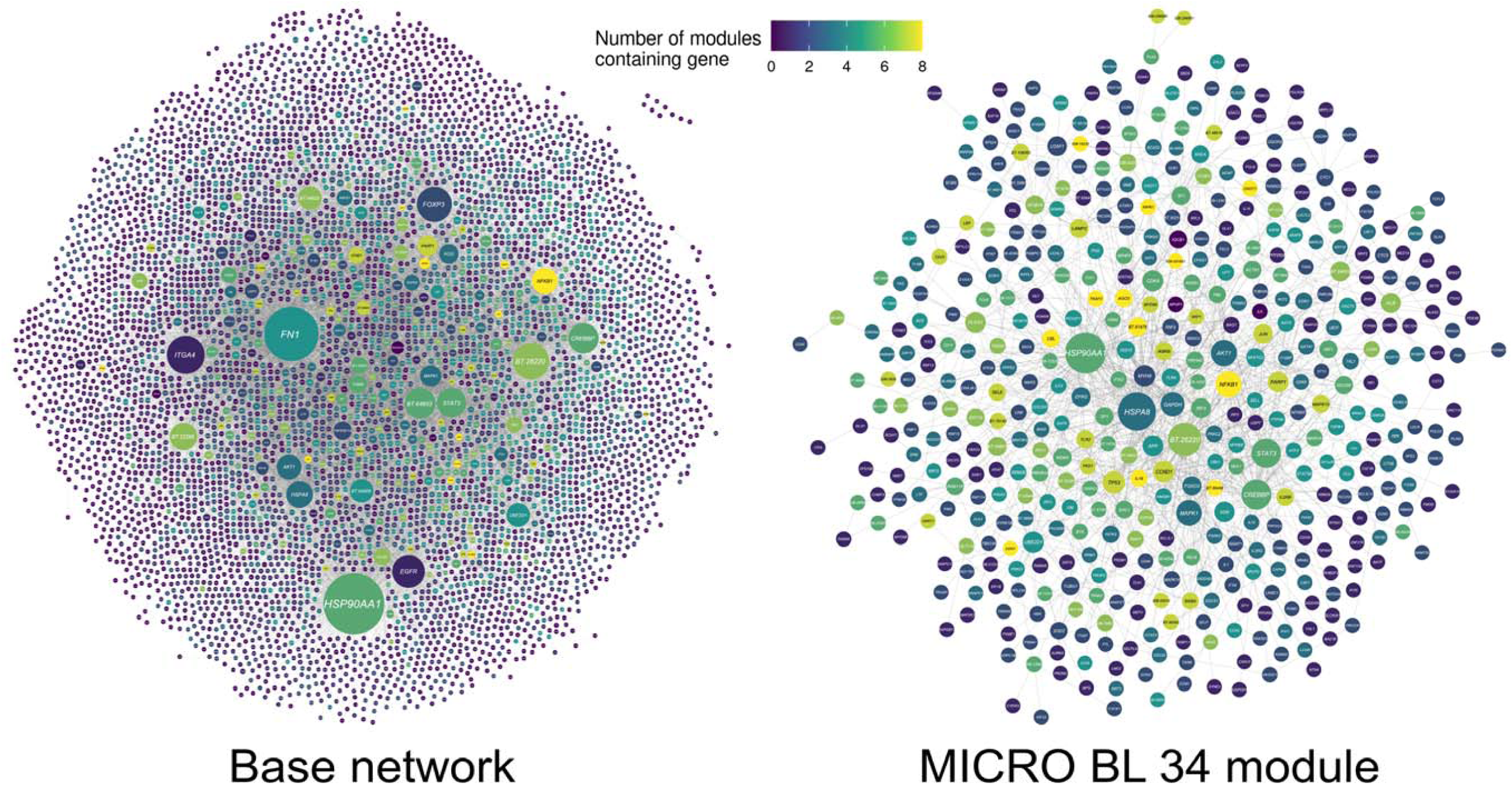
Base network generated using InnateDB with the top results of a search of the GeneCards^®^ for genes relating to the term “trypano*” and functional module identified using jActiveModules and the differential expression results for the MICRO BL 34 contrast. Each node in the networks represents a gene coloured according to the number of functional modules that contain that gene. The edges connecting the nodes represent gene interactions and the nodes are sized according to their number of interactions or degree. The base network contains 5,666 genes (nodes) and 15,651 interactions (edges) while the MICRO BL 34 module contains 474 genes and 1,166 interactions

**Fig 5.**
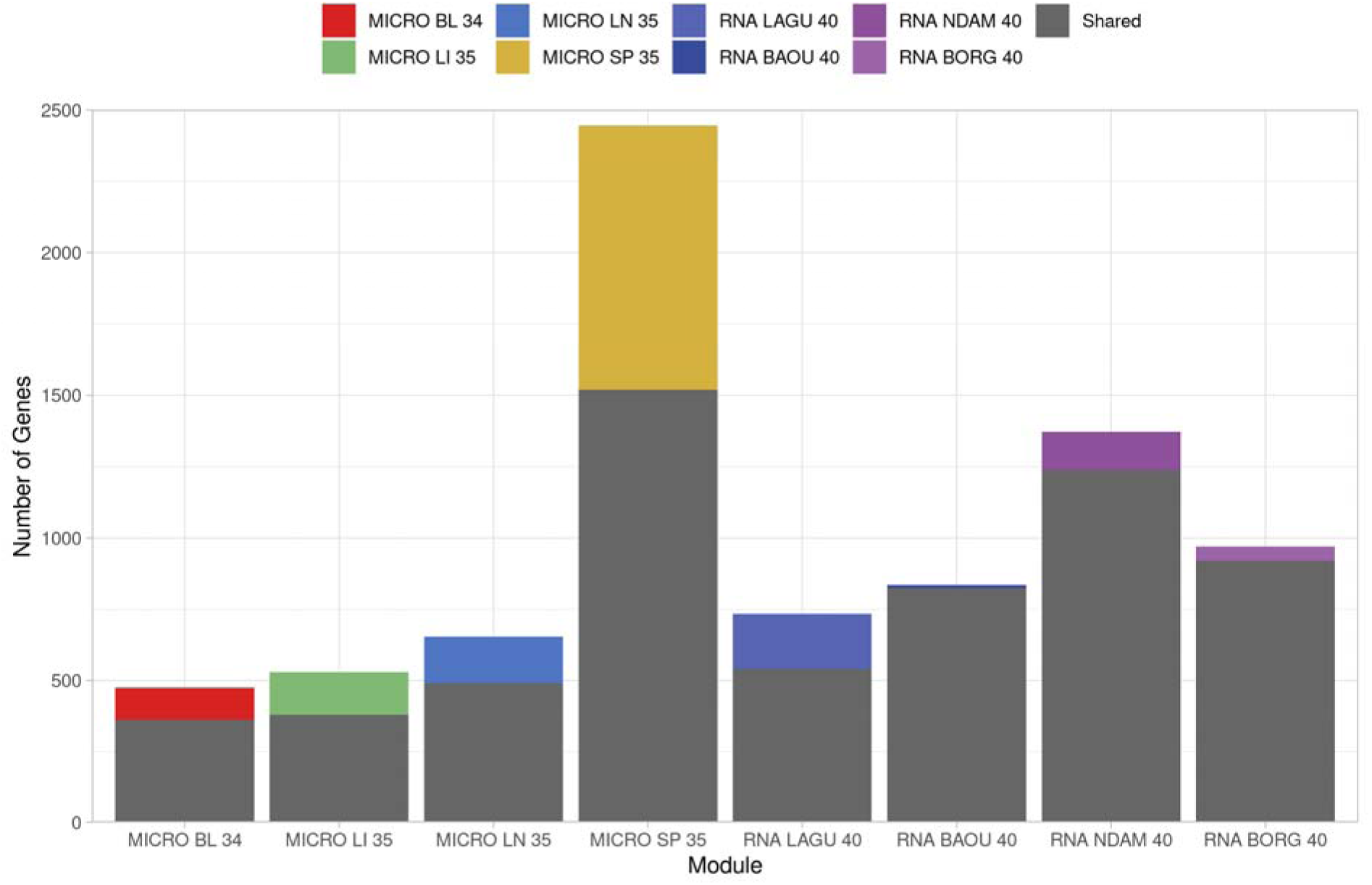
Bar chart showing the number of genes in the functional modules identified using jActiveModules and the differential expression results for each of the final response contrasts for the microarray and RNA-seq data. The colours indicate the number of genes unique to each module or shared between multiple modules.Functional enrichment of gene expression data highlighted the immune system

There were no GO terms significantly enriched for the combined set of significant DEGs across all the response contrasts in the RNA-seq data set against the background set of detectable genes. However, a single GO term (*GO:0030667 secretory granule membrane*) was significantly enriched when no background set was used (**Fig S28**). The GO terms significantly enriched for the genes in the modules identified for each of the final contrasts in the microarray and RNA-seq data against the background set of the base network are shown in **Fig 6**.

**Fig 6.**
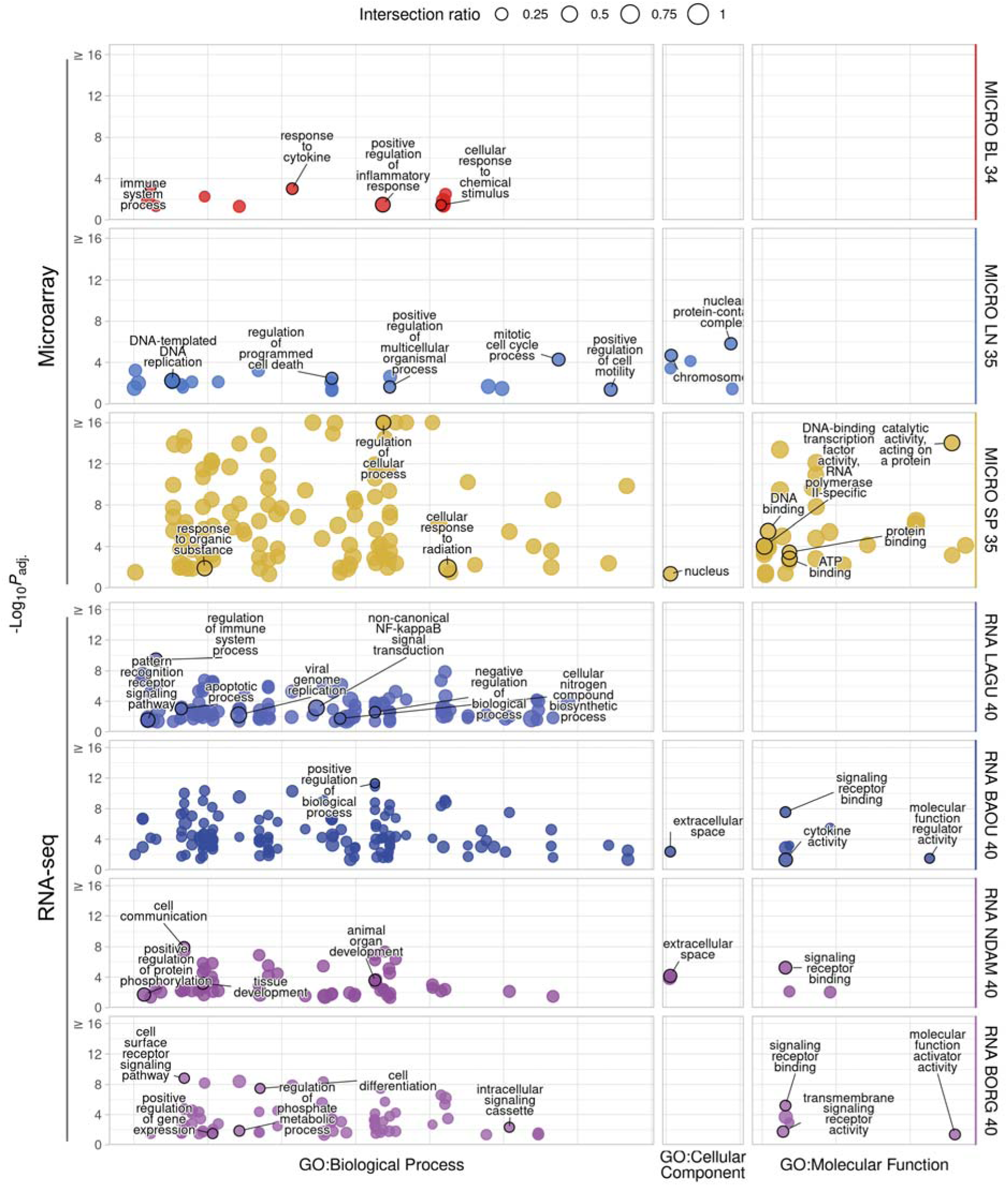
g:Profiler functional enrichment of the genes in the functional modules identified using jActiveModules and the differential expression results for each of the final response contrasts for the microarray and RNA-seq data. Each dot represents a significantly enriched GO term with the size indicating the ratio of the intersection between the term and the introgressed genes. The y-axis shows the -log_10_P_adj._ value up to a maximum of 16 and the panels along the y-axis and colours indicate the module. The panels along the x-axis indicate the source of the term and the position within the panels groups terms from the same GO subtree. The top driver GO terms up to a maximum of ten are indicated with a black outline and label.Results from the integration of genomic and gene expression data showed significant functional overlaps

The driver GO terms significantly enriched for the genes in the MICRO BL 34 module include those related to the immune system (*GO:0002376 immune system process*, *GO:0034097 response to cytokine*, and *GO:0050729 positive regulation of inflammatory response*) and cellular response to chemical stimulus (*GO:0070887 cellular response to chemical stimulus*) (**Fig 6**). The driver GO terms significantly enriched for the genes in the MICRO LN 35 module include those related to the cell cycle and cellular organisation (*GO:1903047 mitotic cell cycle process*, *GO:0043067 regulation of programmed cell death*, *GO:0006261 DNA-templated DNA replication*, *GO:0051240 positive regulation of multicellular organismal process*, *GO:2000147 positive regulation of cell motility*, *GO:0140513 nuclear protein-containing complex*, and *GO:0005694 chromosome*) (**Fig 6**). Although no GO terms were significantly enriched for the genes in the MICRO LI 35 module against the background set of the base network, there were many GO terms significantly enriched when no background set was used (**Fig S29**). In addition, 491 of the 505 GO terms significantly enriched for the MICRO LI 35 module genes were also among the 2,044 GO terms significantly enriched for the genes in the base network (**Fig S30**). The driver GO terms significantly enriched for the genes in the MICRO SP 35 module include those related to cellular responses and binding (*GO:0050794 regulation of cellular process*, *GO:0071478 cellular response to radiation*, *GO:0010033 response to organic substance*, *GO:0005634 nucleus*, *GO:0140096 catalytic activity, acting on a protein*, *GO:0003677 DNA binding*, *GO:0000981 DNA-binding transcription factor activity, RNA polymerase II-specific*, *GO:0005515 protein binding*, and *GO:0005524 ATP binding*) (**Fig 6**).

The driver GO terms significantly enriched for the genes in the RNA LAGU 40 module include those related to the immune system and cell signalling (*GO:0002682 regulation of immune system process*, *GO:0038061 non-canonical NF-kappaB signal transduction*, *GO:0006915 apoptotic process*, *GO:0048519 negative regulation of biological process*, *GO:0019079 viral genome replication*, *GO:0044271 cellular nitrogen compound biosynthetic process*, and *GO:0002221 pattern recognition receptor signalling pathway*) (**Fig 6**). The driver GO terms significantly enriched for the genes in the RNA BAOU 40 module include those related to cell regulation, cell signalling, and cytokine activity (*GO:0048518 positive regulation of biological process*, *GO:0005615 extracellular space*, *GO:0005102 signalling receptor binding*, *GO:0098772 molecular function regulator activity*, and *GO:0005125 cytokine activity*) (**Fig 6**). The driver GO terms significantly enriched for the genes in the RNA NDAM 40 module include those related to cell signalling and tissue and organ development (*GO:0007154 cell communication*, *GO:0048513 animal organ development*, *GO:0009888 tissue development*, *GO:0001934 positive regulation of protein phosphorylation*, *GO:0005615 extracellular space*, and *GO:0005102 signalling receptor binding*) (**Fig 6**). The driver GO terms significantly enriched for the genes in the RNA BORG 40 module include those related to cell signalling and regulation (*GO:0007166 cell surface receptor signalling pathway*, *GO:0030154 cell differentiation*, *GO:0141124 intracellular signalling cassette*, *GO:0019220 regulation of phosphate metabolic process*, *GO:0010628 positive regulation of gene expression*, *GO:0005102 signalling receptor binding*, *GO:0004888 transmembrane signalling receptor activity*, and *GO:0140677 molecular function activator activity*) (**Fig 6**).

A summary of the input data for the integration of the genomic and gene expression data is shown in **Table S5**. This integration generated several significant overlaps between the intervals within 1 Mb up- and downstream from genes with a mean local ancestry *z*-score ≥ 2.0 and the genes contained in the modules identified using the gene expression data (**Table 3**). These overlaps include the *B. indicus* ancestry of the trypanosusceptible FULA population, which was significantly enriched for genes in both the RNA BAOU 40 and RNA BORG 40 modules (**Table 3**). Similarly, the *B. indicus* ancestry of the trypanosusceptible populations in the original LAI analyses (McHugo *et al*. 2025a) were significantly enriched for the genes in the MICRO LI 35 module (**Table 3**). This was true for all combinations of LAI software and SNP data density examined (**Table 3**) and indicates our integrative methodology can detect genomic loci associated with interpopulation differences in susceptibility to trypanosomiasis. The *B. indicus* ancestry of the selected residually admixed European populations analysed with ELAI using low-density SNP data (McHugo *et al*. 2025a) was significantly enriched for genes in both the MICRO LN 35 and RNA LAGU 40 modules (**Table 3**). The African *B. taurus* ancestry of the selected trypanotolerant African hybrids analysed with ELAI using low-density SNP data (McHugo *et al*. 2025a) was significantly enriched for genes in the MICRO SP 35 module (**Table 3**), again suggesting this approach can help dissect the genetic architecture of trypanotolerance. Finally, the European *B. taurus* ancestry of the trypanotolerant LAGU population was significantly enriched for genes in the MICRO LN 35 module (**Table 3**).

**Table 3.**
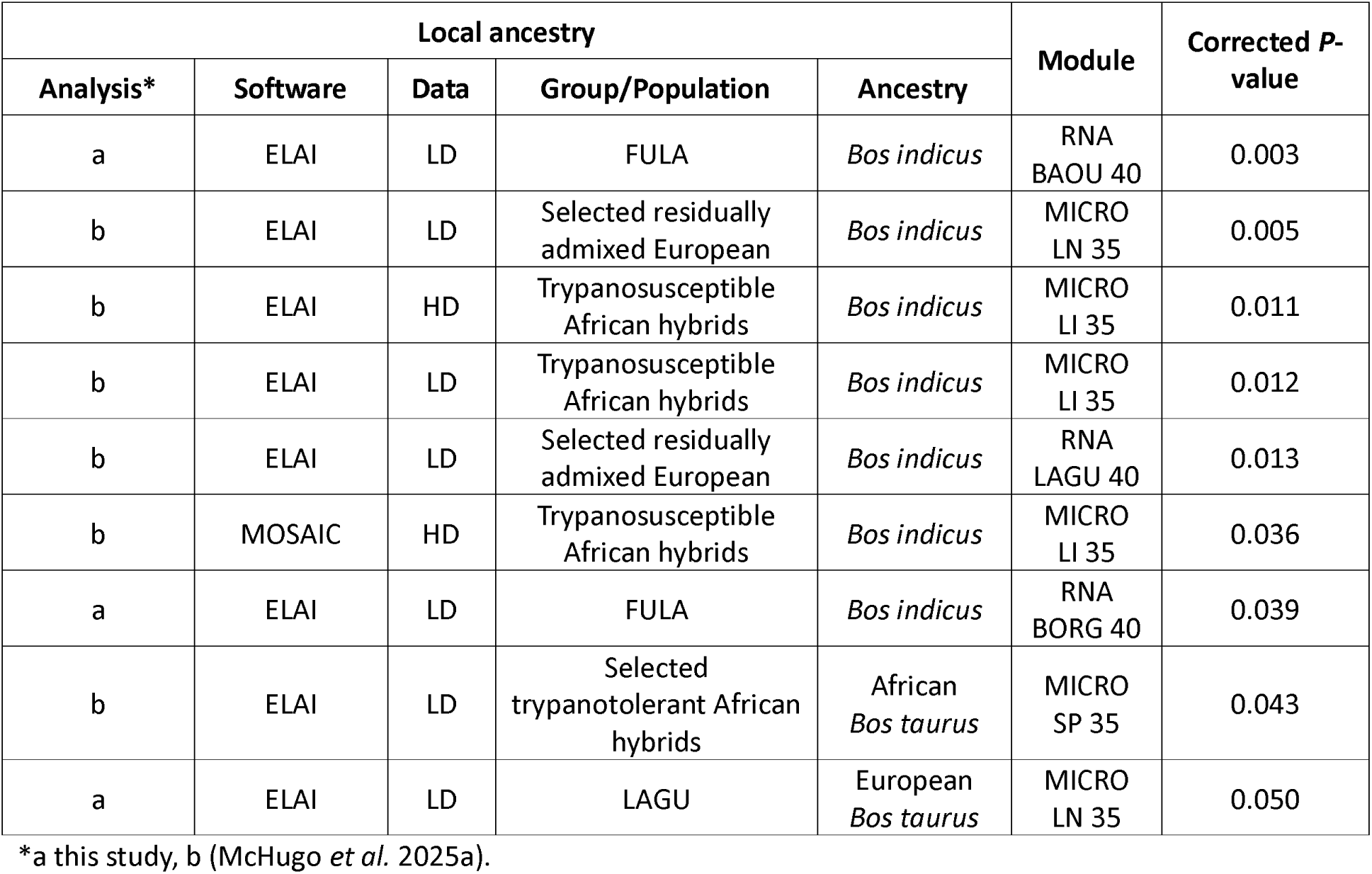
Numbers of SNPs with z-score ≥ 2.0 for mean European B. taurus, African B. taurus and B. indicus ancestry components for the six populations with gene expression data available across all autosomes. The numbers in brackets indicate the percentage of the total number of SNPs in the data set.

## Discussion

### Population genomics analyses confirm the recent evolutionary history of domestic cattle

The results of the population genomics analyses of the SNP data were consistent both with the results from our previous analysis (McHugo *et al*. 2025a) and multiple published studies of hybrid cattle populations (Hanotte *et al*. 2002; Decker *et al*. 2014; Barbato *et al*. 2020; Kim *et al*. 2020; Ward *et al*. 2022). Visualisation of the PCA results by plotting PC1 and PC2 recovered the classic “*Bos* triangle” with the first two PCs explaining a very high proportion of the total variation for PC1–10 within the data (71.51%) and separating the reference European *B. taurus*, African *B. taurus*, and *B. indicus* populations (**Fig 2**, **S3–S5**). Also, as we previously observed (McHugo *et al*. 2025a), the locations of the hybrid populations—nearer to the reference populations they share the most ancestry with—is in agreement with multiple published studies (**Fig 2**) (Gautier *et al*. 2009; Decker *et al*. 2014; Flori *et al*. 2014; Berthier *et al*. 2015; Sempere *et al*. 2015; Bahbahani *et al*. 2017; Upadhyay *et al*. 2017; Verdugo *et al*. 2019; Barbato *et al*. 2020; Ward *et al*. 2022; Wragg *et al*. 2022).

The results of the genetic structure analysis for *K* = 3 mirrored those of the PCA with the separation of the African and European *B. taurus* and *B. indicus* populations, which is also consistent with our previous results (McHugo *et al*. 2025a) (**Fig S6–S9**). Again, the hybrid populations clearly showed evidence of global admixture proportions that was in agreement with both their positions on the PCA and previous studies (**Fig 2**) (Gautier *et al*. 2009; Decker *et al*. 2014; Flori *et al*. 2014; Berthier *et al*. 2015; Sempere *et al*. 2015; Bahbahani *et al*. 2017; Upadhyay *et al*. 2017; Verdugo *et al*. 2019; Barbato *et al*. 2020; Ward *et al*. 2022; Wragg *et al*. 2022). As in our previous study (McHugo *et al*. 2025a), the high modelled *K* values required to best explain the variation among the populations indicated that some of the populations are closely related to the point that they may not be genetically distinct discrete populations (Raj *et al*. 2014).

The results of the LAI analysis demonstrated similar patterns of peaks and troughs dispersed across the genome for each population as those found in our previous analysis (McHugo *et al*. 2025a) using ELAI with high- and low-density SNP data and using MOSAIC with high-density SNP data (**Fig 3**). The African *B. taurus* populations (LAGU and BAOU) exhibited similar low peaks of European *B. taurus* and *B. indicus* ancestry while the African trypanotolerant hybrid NDAM population had slightly higher *B. indicus* peaks (**Fig 3**). This is consistent with previous studies of the origins of these populations as well as the results of the population genomics analyses (**Fig 2**, **S3–S9**) (Gautier *et al*. 2009; Berthier *et al*. 2015; Verdugo *et al*. 2019; Barbato *et al*. 2020). The African trypanotolerant hybrid BORG population showed even higher peaks of *B. indicus* ancestry, which was markedly similar to the mean LAI results obtained previously for the selected trypanotolerant hybrid group (McHugo *et al*. 2025a) (**Fig 3**). This was not unexpected as this group included some of the same samples of the BORG population. In a similar manner, both the LAI results of the trypanosusceptible African hybrid populations (FULA and BORA) resembled the mean LAI results for the trypanosusceptible African hybrid group from our previous results (McHugo *et al*. 2025a); again, this group included some of the same BORA samples (**Fig 3**). Consequently, the proportions of the numbers of SNPs passing the genome-wide *z*-score ≥ 2.0 threshold, as well as the numbers of genes within 1 Mb up- and downstream from these SNPs, were similar to our previous results (McHugo *et al*. 2025a) (**Table S1**, **S2**). The lack of SNPs, and therefore genes, that passed the threshold for the African *B. taurus* ancestry component in the African *B. taurus* populations was likely due to the high and relatively uniform proportion of the African *B. taurus* ancestry component across the genome. This would give rise to a situation such that no SNPs could pass the threshold of two standard deviations from the mean (**Fig 3**, **Table S1**, **S2**). Similarly, the lower numbers of SNPs that exceeded the threshold for the African *B. taurus* and *B. indicus* ancestry components for the trypanotolerant and trypanosusceptible African hybrid populations, respectively, was likely due to the higher proportions of the reference population ancestries to which each hybrid group is most closely related (**Fig 3**, **Table S1**, **S2**).

The introgressed genomic regions also showed similar patterns in terms of functional enrichment as our previous LAI results (McHugo *et al*. 2025a) with less, or even no GO terms enriched for the ancestry component most similar to the population (**Fig S10–S12**). Significant driver GO terms enriched for genes near the peaks of the other ancestry contributions included those relating to the MHC and other components of the immune system, which was also in agreement with previous studies of local ancestry in cattle (**Fig S10–S12**) (Chen *et al*. 2020; Buggiotti *et al*. 2021; Guan *et al*. 2022; Li *et al*. 2023). It is notable that immune system functions were well represented in the top functional enrichment categories for the introgressed genomic regions since there are well documented differences among European *B. taurus*, African *B. taurus*, and *B. indicus* cattle populations in terms of susceptibilities to various infectious diseases, including bovine tuberculosis caused by *M. bovis* (Allen *et al*. 2010; Lee *et al*. 2024); East Coast fever and tropical theileriosis caused by *Theileria parva* and *Theileria annulate*, respectively (Bahbahani & Hanotte 2015); and African animal trypanosomiasis (AAT) caused by *Trypanosoma* spp. (Yaro *et al*. 2016). Similarly, it is notable that, as for the previous LAI analysis (McHugo *et al*. 2025a), significant driver GO terms relating to olfaction were identified and genes related to olfaction have been identified by previous local ancestry studies of both cattle and humans and by selection signature studies in cattle (**Fig S10–S12**) (Yang *et al*. 2014; Melo *et al*. 2017; Pan *et al*. 2022; Sun *et al*. 2023). Although this may simply be due to the large number of olfactory receptor genes dispersed throughout the cattle genome (Niimura & Nei 2007; Lee *et al*. 2013), recent studies have suggested that up to 580 olfactory receptors may be expressed by macrophages, immune cells involved in the detection and phagocytosis of pathogens (Orecchioni *et al*. 2022). In addition, macrophages are the host’s first line of defence to mycobacterial infection with evasion and reprogramming of host macrophages being a key component of intracellular mycobacterial infections that cause tuberculosis disease (Hall *et al*. 2024) and variation in olfactory receptor genes were found to be associated with bovine tuberculosis (Ring *et al*. 2019).

Other significant driver GO terms detected in the present study that were common with the previous LAI analysis (McHugo *et al*. 2025a) included those relating to cell signalling, L-amino acid transmembrane transport, oxidoreductase activity, metabolic processes, cellular component organisation, haemoglobin complex and homophilic cell adhesion via plasma membrane adhesion molecules (**Fig S10–S12**). A notable difference to our previous results was the enrichment of GO terms related to the skin and keratin for the genes near peaks of *B. indicus* ancestry for the trypanotolerant hybrid populations (NDAM and BORG) (**Fig S11**). This may be a result of selection for heat tolerance traits such as the “slick” phenotype associated with mutation in the prolactin receptor gene (*PRLR*) in Latin American Criollo cattle, which derive some of their ancestry from African cattle (Porto-Neto *et al*. 2018; Ward *et al*. 2024; Gebeyehu *et al*. 2025).

### Differential expression analysis highlighted immune genes

As with the microarray data (McHugo *et al*. 2025b) and as expected due to their design, the use of response contrasts to identify changes in expression over time in the trypanotolerant population samples (LAGU, BAOU, NDAM, and BORG) relative to the trypanosusceptible FULA in the differential expression analysis of the RNA-seq data found lower numbers of significant DEGs than the direct contrasts between or within the populations (Rue-Albrecht *et al*. 2014; Peylhard *et al*. 2023). However, despite the low numbers, the significant DEGs followed the established pattern also observed for the microarray data and direct RNA-seq contrasts, with increasing numbers of significant DEGs over the course of the infection leading to the highest numbers of significant DEGs observed for the final time point (O’Gorman *et al*. 2009; Peylhard *et al*. 2023). In addition, it is known that trypanotolerant breeds with high levels of *B. taurus* ancestry have enhanced abilities to control anaemia while hybrid animals exhibit intermediate levels of control when compared to susceptible *B. indicus* breeds (Berthier *et al*. 2015). As the LAGU, BAOU, and NDAM populations are known to have higher levels of *B. taurus* ancestry, which has also been shown by the population genomics analyses, it is therefore unsurprising that these populations exhibited more significant DEGs than the more admixed BORG population (Flori *et al*. 2014; Berthier *et al*. 2015; Gautier 2015; Bahbahani *et al*. 2017; Verdugo *et al*. 2019; Barbato *et al*. 2020; Ward *et al*. 2022).

For the RNA-seq results, as with the results of the microarray analysis (McHugo *et al*. 2025b), there were overlaps in the significant DEGs between the contrast types. These include the duplicated genes *NEIL2* (Nei like DNA glycosylase 2), which is involved in the immune system, the tumour suppressor *LZTS3* (leucine zipper tumour suppressor family member 3), and *SLC11A1* (solute carrier family 11 member 1), which encodes a regulator of iron homeostasis in macrophages (Archer *et al*. 2015; Gu *et al*. 2023; Tapryal *et al*. 2023). In particular, variants of *SLC11A1* are associated with susceptibility to infectious diseases, including tuberculosis in cattle (Archer *et al*. 2015; Holder *et al*. 2020). Of the 43 unique genes identified as significantly differentially expressed by the response contrasts only 10 were not identified as significantly differentially expressed by any of the direct contrasts used by Peylhard and colleagues (Peylhard *et al*. 2023). These genes include *ANXA6* (annexin A6), *NT5C2* (5’-nucleotidase, cytosolic II), *PIGR* (polymeric immunoglobulin receptor), and *TET2* (tet methylcytosine dioxygenase 2), which were identified as significantly differentially expressed for various contrasts using a gene expression microarray (McHugo *et al*. 2025b) and are known to play roles in the immune system and haematological disorders (Sphyris & Mani 2011; Mercher *et al*. 2012; Jordheim 2018; Cong *et al*. 2021; Rashidi *et al*. 2023). Notably, anaemia is the main cause of death due trypanosomiasis and the ability to control this anaemia is thought to be critical to trypanotolerance in cattle (Naessens 2006; Stijlemans *et al*. 2015).

Other genes, which were not detectable with microarray data, included *GPATCH2L* (g-patch domain containing 2 like), *NFAT5* (nuclear factor of activated T cells 5), *PICALM* (phosphatidylinositol binding clathrin assembly protein), *PTPN4* (protein tyrosine phosphatase non-receptor type 4), *SLC38A4* (solute carrier family 38 member 4), and *TTBK2* (tau tubulin kinase 2). Interestingly, *TTBK2* has been identified as a candidate gene underlying trypanotolerance in the Sheko cattle breed and mutations in this gene have been hypothesised to be associated with response to the presence of trypanosome parasites in the brain white matter, cerebral fluid, thyroid, and parathyroid glands (Mekonnen *et al*. 2019). In addition, *NFAT5* encodes a member of a group of transcription factors that have been shown to mediate production of cytokines during trypanosome infection (Kayama *et al*. 2009), while *PICALM* interacts with phosphatidylinositol 4-kinases that are thought to be drug targets for human trypanosomiasis (Li *et al*. 2024). The *PTPN4* gene is involved in the immune system (Huai *et al*. 2015) and *GPATCH2L* has been highlighted in a study of haemorrhagic fever (Redwan *et al*. 2019). Finally, *SLC38A4* is related to both *SLC40A1* and *SLC11A1*, which were observed to be differentially expressed in the microarray data (McHugo *et al*. 2025b) and direct RNA-seq contrasts, respectively (Peylhard *et al*. 2023). Finally, the lack of enriched GO terms enriched for the significant DEGs from the RNA-seq data was likely due to the low number of genes.

### Network analysis identified modules of genes involved in immunobiology

The base GIN network for genes associated with trypanosome infection and trypanosomiasis contained a similar number of genes and interactions as previous GINs generated for integrative genomics studies of *M. bovis* infection in cattle by Hall *et al*. (Hall *et al*. 2021; Hall *et al*. 2024). In common with many GINs obtained using transcriptomics data, we observed a scale-free topology for functional modules (Yang 2020). This indicates that most genes within each functional module interact with only one other gene, while a small subset of genes interact with substantially more (Barabasi & Albert 1999; Albert 2005; Hall *et al*. 2024). However, the functional modules identified for each of the final contrasts in the microarray and RNA-seq data in the present study were considerably larger than those identified by Hall *et al*. (**Fig 5**) (Hall *et al*. 2021; Hall *et al*. 2024). When it is unable to adequately detect larger subnetworks, the jActiveModules algorithm can still retrieve subnetworks consisting of a small number of genes, or even single genes; therefore, the large modules identified in the present study indicate that the microarray and RNA-seq data sets are robust (Ideker *et al*. 2002). In addition, modules identified from RNA-seq data tend to be larger than those identified from microarray data (Hatem *et al*. 2012), and this was the case for all the modules apart from that identified using the final spleen microarray contrast. The large number of genes shared between the modules is also unsurprising as they were drawn from the same base network. Also, the significant DEGs identified using the microarray data had a large number of genes in common between the tissues (McHugo *et al*. 2025b). Our results are also in agreement with previous studies that observed high numbers of genes in common between modules identified from a shared base network (Hall *et al*. 2021; Hall *et al*. 2024).

The eight genes shared between all the modules include genes related to the immune system that have already been highlighted by previous studies of response to trypanosome infection in cattle, such as *IL1B* (O’Gorman *et al*. 2006) and *NFKB1* (O’Gorman *et al*. 2006; Peylhard *et al*. 2023) (**Fig 5**, **Table S4**). Other genes have been similarly highlighted by studies of trypanosome infection in mice, including *EDN1* (Corral *et al*. 2013) and *TRAF2* (Santamaría *et al*. 2023), as well as by studies of other bovine infections, such as *CBL* which was highlighted by a study of bovine anaplasmosis caused by *Anaplasma marginale* (Ahlawat *et al*. 2023) and *CNOT1* which was highlighted by a study of Johne’s disease caused by *Mycobacterium avium paratuberculosis* (MAP) (Kleinwort *et al*. 2019). Finally, *RIPK1* is known to play a role in inflammation (Newton 2015) while *AGO2* has been recently found to be involved in regulating immune response (Wang *et al*. 2025).The driver GO terms significantly enriched for the genes in the microarray modules were similar to those found to be enriched for the significant DEGs in the microarray results outputs used to generate the modules (**Fig 6**) (McHugo *et al*. 2025b). Therefore, the presence of driver GO terms related to the immune system, and cytokines in particular, among those significantly enriched for genes in the MICRO BL 34 is expected as these genes were significantly differentially expressed for both the microarray data (McHugo *et al*. 2025b), the RNA-seq data and other published studies (Uzonna *et al*. 1999; O’Gorman *et al*. 2006; O’Gorman *et al*. 2009; Noyes *et al*. 2011; Peylhard *et al*. 2023). In addition, the anaemia caused by trypanosome infection can be considered an immune response, which may be driven by cytokines (Stijlemans *et al*. 2008; Stijlemans *et al*. 2010; Stijlemans *et al*. 2015). The enrichment of driver GO terms related to the cell cycle and cellular organisation for genes in the MICRO LN 35 module was also in agreement with the results of the analysis of the microarray data (Noyes *et al*. 2011) (**Fig 6**). The lack of GO terms significantly enriched for genes in the MICRO LI 35 module against the background set of the base network was likely due to the similarity between the module and the base network, as evidenced by the almost total overlap in significantly enriched GO terms with the base network when no background set was used (**Fig S29**, **S30**). The driver GO terms significantly enriched for the genes in the MICRO SP 35 module included those related to cellular responses and binding (**Fig 6**). This aligns with the small number of GO terms enriched for the small number of significant DEGs in the spleen samples in the previous microarray results (McHugo *et al*. 2025b) and highlights the utility of the jActiveModules method for incorporating genes that are highly interconnected with the significant DEGs but that were otherwise not detected by the differential expression analysis (Hall *et al*. 2021; Hall *et al*. 2024). The power of a network-based approach using jActiveModules was also evident from the modules identified using the RNA-seq data, where fewer significant DEGs were detected compared to the microarray data, meaning that incorporating highly interconnected genes was of greater benefit.

The driver GO terms significantly enriched for the genes in the RNA LAGU 40 module included those related to the immune system and cell signalling, while the driver GO terms significantly enriched for the genes in the RNA BAOU 40 module included those related to cell regulation, cell signalling, and cytokine activity (**Fig 6**). The driver GO terms significantly enriched for the genes in the RNA NDAM 40 module also included those related to cell signalling, in addition to tissue and organ development. The driver GO terms significantly enriched for the genes in the RNA BORG 40 module also included those related to cell signalling and regulation (**Fig 6**). That driver GO terms related to cell signalling and regulation were common among the terms enriched for the genes in the modules is logical as the modules were identified from a base network made of interacting genes that are frequently linked by cell signalling and regulatory pathways.

Similarly, the presence of driver GO terms related to the immune system among those significantly enriched for the genes in the RNA-seq modules is to be expected as the modules were identified using the differential expression results from an infection experiment that identified several immune genes as significantly differentially expressed.

### Integration revealed significant overlap in the results from the genomic and gene expression data

The presence of genes related to the immune system in both the modules identified with the microarray and RNA-seq data and the LAI results may explain the significant enrichment found in the intervals within 1 Mb up- and downstream from genes with a mean local ancestry *z*-score ≥ 2.0 and the genes contained in the modules identified using the gene expression data (**Table 3**). It is notable that immune genes were located in genomic regions around the peaks of the LAI results of hybrid cattle populations and that these regions were also enriched for genes identified by functional modules of response contrasts for the same hybrid populations for both RNA-seq and microarray gene expression data using multiple tissues and populations.

We hypothesised that the genes within 1 Mb of the top African *B. taurus* ancestry peaks in the trypanotolerant populations could underpin the complex trypanotolerance trait, particularly if these regions showed significant enrichment for the genes identified during the differential expression and network analyses. However, there was only one such significant overlap: the African *B. taurus* ancestry of the selected trypanotolerant African hybrids analysed with ELAI using low-density SNP data (McHugo *et al*. 2025a) was significantly enriched for genes in the MICRO SP 35 module (**Table 3**). These genes may therefore be considered as candidate genes for trypanotolerance. Although it must also be noted that the MICRO SP 35 module was the largest and the number of genes contained in this module may have increased the probability of overlap with the local ancestry peaks, despite the *P*-value correction to account for the number of genes examined (Lee *et al*. 2012).

Interestingly, the European *B. taurus* ancestry of the trypanotolerant LAGU population was enriched for genes in the MICRO LN 35 module (*P*_adj._ = 0.05; **Table 3**). While it is possible that these genes may also be candidate genes for trypanotolerance as they have been found within peaks of *B. taurus* ancestry, which can also be considered troughs of trypanosusceptible *B. indicus* ancestry, this is unlikely. The pure reference European and African *B. taurus* populations were clearly separated by the various population genomics analyses and it is likely that that the LAI algorithms could distinguish between the trypanosusceptible European *B. taurus* and trypanotolerant African *B. taurus* ancestry. In a similar manner, the peaks of *B. indicus* ancestry in the selected residually admixed European populations analysed with ELAI using low-density SNP data (McHugo *et al*. 2025a) were significantly enriched for genes in both the MICRO LN 35 and RNA LAGU 40 modules (**Table 3**). A possible explanation for this is that the immune genes in these modules overlapped with those in the regions around the local ancestry peaks of *B. indicus* ancestry as these regions were enriched for GO terms relating to the MHC. This makes sense as increased diversity in the MHC region is associated with disease resistance and MHC genes are under balancing selection in cattle (Ellis 2004; Codner *et al*. 2012; Ellis & Hammond 2014).

It is also notable that the peaks of *B. indicus* ancestry in several trypanosusceptible populations were significantly enriched for genes in the functional modules. These overlaps include the trypanosusceptible FULA population, which was significantly enriched for genes in both the RNA BAOU 40 and RNA BORG 40 modules (**Table 3**). Similarly, the peaks of *B. indicus* ancestry in the trypanosusceptible populations in the original LAI analyses (McHugo *et al*. 2025a) were significantly enriched for the genes in the MICRO LI 35 module for all combinations of LAI software and SNP data density examined (**Table 3**). It is also possible that the genes in these additional functional modules can be considered as candidate genes for trypanotolerance since they are found in the peaks of trypanosusceptible *B. indicus* ancestry in multiple trypanosusceptible hybrid populations. This implies that these regions of increased trypanosusceptible *B. indicus* introgression in multiple trypanosusceptible hybrid populations were significantly enriched for genes identified as part of downstream network analyses of differential expression analyses comparing the response of trypanosusceptible and trypanotolerant cattle to trypanosome infection using multiple populations, tissues and data types. Therefore, because these genomic regions were found to be significantly enriched for multiple functional modules, it suggests that the genes shared by multiple modules may be prioritised as candidate genes for trypanotolerance. The similarity of these genomic regions is highlighted by the fact that the driver GO term *GO:0007156 homophilic cell adhesion via plasma membrane adhesion molecules* and the upstream GO term *GO:0098742 cell-cell adhesion via plasma-membrane adhesion molecules* were enriched for the genes located in the regions around the peaks of *B. indicus* ancestry in the FULA population and the trypanosusceptible African hybrid populations in the LAI analysis using high-density SNP data with MOSAIC and low-density SNP data with ELAI (McHugo *et al*. 2025a). The *GO:0098742* term was also found in another study to be enriched for genes in introgressed African taurine genomic regions in hybrid African cattle populations (Friedrich *et al*. 2023).

### Limitations of this study

The complex nature of the trypanotolerance trait coupled with the many selective forces in the form of other pathogens acting on the immune systems of hybrid African cattle mean these results must be interpreted with caution. For example, trypanotolerant breeds are also less susceptible to other infectious diseases such as helminthiasis and tick-borne-diseases and genes associated with the immune response have been found to be under selection in West African cattle populations (Gautier *et al*. 2009; Kim *et al*. 2017b). This suggests an alternative hypothesis such that the immune genes found in the peaks of African *B. taurus* ancestry in the trypanotolerant populations may be associated with resistance to another infectious disease. In addition, an alternative hypothesis can be proposed that the genes enriched for *B. indicus* ancestry the trypanosusceptible populations may confer resistance to bovine tuberculosis (bTB), rather than trypanotolerance; *B. indicus* cattle have long been known to have lower susceptibility to bTB compared to *B. taurus* cattle (Liston & Soparkar 1917; Allen *et al*. 2010). *Bos indicus* and *B. taurus* cattle exhibit differing immune responses to infection with *M. bovis*, the causative agent of bTB, and the genes underlying this polygenic disease resistance trait remain unknown (Vordermeier *et al*. 2012; Kumar *et al*. 2023). Notably, integration of data from different experiments involving infectious diseases in cattle, including trypanosomiasis and tuberculosis, found many of the same genes were differentially expressed, suggesting common immune mechanisms in response to these infections (Beiki *et al*. 2016; Beiki *et al*. 2018).

## Conclusion

In conclusion, integration of genomic data in the form of high- and low-density SNP data from a range of trypanotolerant and trypanosusceptible cattle populations with RNA-seq and microarray transcriptomics data from the same populations has provided a new approach for identification of trypanotolerance candidate genes.

## Acknowledgements

We thank Morris Agaba, Olivier Hanotte, Stephen J. Kemp, Daniel G. Bradley, and Stephen V. Gordon for assistance with sample resources and for useful scientific discussion. This research work was funded by Science Foundation Ireland (SFI) under Investigator Programme Awards (grant nos: SFI/01/F.1/B028 and SFI/15/IA/3154). JAW was supported by the Centre for Research Training in Genomics Data Science (grant no. SFI/18/CRT/6214).

## Data Availability Statement

No new data were generated for this study. The computer code required to repeat and reproduce the analyses is available at doi.org/10.5281/zenodo.11517978.

## Ethics Statement

As no new data were generated for this study, no ethical approval was required.

## Supporting information captions

**Table S1.** Numbers of SNPs with z-score ≥ 2.0 for mean European *B. taurus*, African *B. taurus* and *B. indicus* ancestry components for the six populations with gene expression data available across all autosomes. The numbers in brackets indicate the percentage of the total 29,869 SNPs in the data set.

**Table S2.** Numbers of genes within 1 Mb up- and downstream of SNPs with z-score ≥ 2.0 for mean European *B. taurus*, African *B. taurus* and *B. indicus* ancestry components for the six populations with gene expression data available across all autosomes. The numbers in brackets indicate the percentage of the total 34,080 genes in the data set.

**Table S3.** Population, days post-infection and the significantly differentially expressed genes with increased and decreased expression with gene symbols for the response contrasts.

**Table S4.** Gene symbol and modules containing each of the 243 genes with valid gene symbols that were found in four or more of the eight functional modules.

**Table S5.** Numbers of intervals within 1 Mb up- and downstream of SNPs with z-score ≥ 2.0 for mean European *B. taurus*, African *B. taurus* and *B. indicus* ancestry components for the six populations with gene expression data available across all autosomes and groups of populations from the original local ancestry analysis.

**Fig S1.** Heatmap of mean identity by state values for SNP data in European, African, and Asian cattle populations.

**Fig S2.** Tukey box plots showing the distribution of inbreeding values (*F*) for SNP data for each population of European, African, and Asian cattle. Outliers are indicated with a black outline.

**Fig S3.** A. Principal component analysis (PCA) of SNP data for cattle coloured according to population showing the first two principal components and B. bar chart of proportion of variance of the top ten principal components.

**Fig S4. A.** Principal component analysis (PCA) of the high-density SNP data for the cattle samples from McHugo and colleagues (McHugo et al. 2025a) coloured according to population showing the first two principal components and **B.** bar chart of proportion of variance of the top ten principal components. The transparency indicates the availability of microarray gene expression data for the sample.

**Fig S5. A.** Principal component analysis (PCA) of the low-density SNP data for the cattle samples from McHugo and colleagues (McHugo et al. 2025a) coloured according to population showing the first two principal components and **B.** bar chart of proportion of variance of the top ten principal components. The transparency indicates the availability of microarray gene expression data for the sample.

**Fig S6.** Hierarchical clustering of the SNP data for European, African, and Asian cattle populations. Results are shown for an assumed value of the number of ancestral populations K = 3. The transparency indicates the availability of gene expression data for the sample.

**Fig S7.** Hierarchical clustering of the SNP data for European, African, and Asian cattle populations. Results are shown for an assumed value of the number of ancestral populations K = 3.

**Fig S8.** Hierarchical clustering of the high-density SNP data from McHugo and colleagues (McHugo et al. 2025a) for European, African, and Asian cattle populations. Results are shown for an assumed value of the number of ancestral populations K = 3. The transparency indicates the availability of microarray gene expression data for the sample.

**Fig S9.** Hierarchical clustering of the low-density SNP data from McHugo and colleagues (McHugo et al. 2025a) for European, African, and Asian cattle populations. Results are shown for an assumed value of the number of ancestral populations K = 3. The transparency indicates the availability of microarray gene expression data for the sample.

**Fig S10.** g:Profiler functional enrichment of introgressed regions in the African *B. taurus* cattle populations with gene expression data available according to local ancestry analysis of SNP data. Each dot represents a significantly enriched GO term with the size indicating the ratio of the intersection between the term and the introgressed genes. The *y*-axis shows the -log_10_*P*_adj._ value up to a maximum of 16 and the panels along the *y*-axis and colours indicate the ancestry component. The panels along the *x*-axis indicate the source of the term and the position within the panels groups terms from the same GO subtree. The top driver GO terms up to a maximum of ten are indicated with a black outline and label.

**Fig S11.** g:Profiler functional enrichment of introgressed regions in the trypanotolerant African hybrid cattle populations with gene expression data available according to local ancestry analysis of SNP data. Each dot represents a significantly enriched GO term with the size indicating the ratio of the intersection between the term and the introgressed genes. The *y*-axis shows the -log_10_*P*_adj._ value up to a maximum of 16 and the panels along the *y*-axis and colours indicate the ancestry component. The panels along the *x*-axis indicate the source of the term and the position within the panels groups terms from the same GO subtree. The top driver GO terms up to a maximum of ten are indicated with a black outline and label.

**Fig S12.** g:Profiler functional enrichment of introgressed regions in the trypanosusceptible African hybrid cattle populations with gene expression data available according to local ancestry analysis of SNP data. Each dot represents a significantly enriched GO term with the size indicating the ratio of the intersection between the term and the introgressed genes. The *y*-axis shows the -log_10_*P*_adj._ value up to a maximum of 16 and the panels along the *y*-axis and colours indicate the ancestry component. The panels along the *x*-axis indicate the source of the term and the position within the panels groups terms from the same GO subtree. The top driver GO terms up to a maximum of ten are indicated with a black outline and label.

**Fig S13.** Bar chart showing the numbers of significantly differentially expressed genes for the response contrasts of the RNA-seq data. The extent of the bar above and below 0 on the *y*-axis indicates the numbers of significantly differentially expressed genes with increased and decreased expression respectively. The position on the *x*-axis indicates the number of days post-infection and the colour and shapes within the bars represent the population.

**Fig S14.** Volcano plot showing the results of the response contrast for the RNA-seq data from the LAGU population at 40 days post-infection. Each dot represents a gene with the position on the *x*- and *y*-axes indicating the log_2_ fold change and -log_10_*P*_adj._, respectively. Genes above the horizontal dashed line are significantly differentially expressed with the colours representing the change in expression. The top 10 most significant genes for increased and decreased expression with gene symbols are labelled.

**Fig S15.** Volcano plot showing the results of the response contrast for the RNA-seq data from the BAOU population at 40 days post-infection. Each dot represents a gene with the position on the *x*- and *y*-axes indicating the log_2_ fold change and -log_10_*P*_adj._, respectively. Genes above the horizontal dashed line are significantly differentially expressed with the colours representing the change in expression. The top 10 most significant genes for increased and decreased expression with gene symbols are labelled.

**Fig S16.** Volcano plot showing the results of the response contrast for the RNA-seq data from the NDAM population at 40 days post-infection. Each dot represents a gene with the position on the *x*- and *y*-axes indicating the log_2_ fold change and -log_10_*P*_adj._, respectively. Genes above the horizontal dashed line are significantly differentially expressed with the colours representing the change in expression. The top 10 most significant genes for increased and decreased expression with gene symbols are labelled.

**Fig S17.** Volcano plot showing the results of the response contrast for the RNA-seq data from the BORG population at 40 days post-infection. Each dot represents a gene with the position on the *x*- and *y*-axes indicating the log_2_ fold change and -log_10_*P*_adj._, respectively. Genes above the horizontal dashed line are significantly differentially expressed with the colours representing the change in expression. The top 10 most significant genes for increased and decreased expression with gene symbols are labelled.

**Fig S18.** Base network generated using InnateDB with the top results of a search of the GeneCards^®^ for genes relating to the term “trypano*”. Each node in the network represents a gene while the edges connecting the nodes represent gene interactions. The nodes are sized according to their degree or number of interactions.

**Fig S19.** Functional module identified using jActiveModules and differential expression results for the MICRO BL 34 contrast. Each node in the network represents a gene coloured according to expression with significant differential expression indicated by the outline. The edges connecting the nodes represent gene interactions and the nodes are sized according to their number of interactions or degree.

**Fig S20.** Functional module identified using jActiveModules and differential expression results for the MICRO LI 35 contrast. Each node in the network represents a gene coloured according to expression with significant differential expression indicated by the outline. The edges connecting the nodes represent gene interactions and the nodes are sized according to their number of interactions or degree.

**Fig S21.** Functional module identified using jActiveModules and differential expression results for the MICRO LN 35 contrast. Each node in the network represents a gene coloured according to expression with significant differential expression indicated by the outline. The edges connecting the nodes represent gene interactions and the nodes are sized according to their number of interactions or degree.

**Fig S22.** Functional module identified using jActiveModules and differential expression results for the MICRO SP 35 contrast. Each node in the network represents a gene coloured according to expression with significant differential expression indicated by the outline. The edges connecting the nodes represent gene interactions and the nodes are sized according to their number of interactions or degree.

**Fig S23.** Functional module identified using jActiveModules and differential expression results for the RNA LAGU 40 contrast. Each node in the network represents a gene coloured according to expression with significant differential expression indicated by the outline. The edges connecting the nodes represent gene interactions and the nodes are sized according to their number of interactions or degree.

**Fig S24.** Functional module identified using jActiveModules and differential expression results for the RNA BAOU 40 contrast. Each node in the network represents a gene coloured according to expression with significant differential expression indicated by the outline. The edges connecting the nodes represent gene interactions and the nodes are sized according to their number of interactions or degree.

**Fig S25.** Functional module identified using jActiveModules and differential expression results for the RNA NDAM 40 contrast. Each node in the network represents a gene coloured according to expression with significant differential expression indicated by the outline. The edges connecting the nodes represent gene interactions and the nodes are sized according to their number of interactions or degree.

**Fig S26.** Functional module identified using jActiveModules and differential expression results for the RNA BORG 40 contrast. Each node in the network represents a gene coloured according to expression with significant differential expression indicated by the outline. The edges connecting the nodes represent gene interactions and the nodes are sized according to their number of interactions or degree.

**Fig S27.** Upset plot showing the top 20 intersections between the genes in the functional modules identified using jActiveModules and differential expression results for each of the final response contrasts for the microarray and RNA-seq data. The horizontal bars indicate the total number of genes in each module while the vertical bars indicate the number of genes in common between the modules annotated with black dots connected by lines in the intersection matrix. The background colour of the stripes in the intersection matrix and colour of the horizontal and vertical bars represent the module. Black bars indicate an overlap between different modules.

**Fig S28.** g:Profiler functional enrichment of significantly differentially expressed genes in the RNA-seq response contrasts with no background data set specified. Each dot represents a significantly enriched GO term with the size indicating the ratio of the intersection between the term and the introgressed genes. The *y*-axis shows the -log_10_*P*_adj._ value up to a maximum of 16 and the panels along the *y*-axis and colours indicate the module. The panels along the *x*-axis indicate the source of the term and the position within the panels groups terms from the same GO subtree. The top driver GO terms up to a maximum of ten are indicated with a black outline and label.

**Fig S29.** g:Profiler functional enrichment of the genes in the MICRO LI 35 functional module with no background data set specified. Each dot represents a significantly enriched GO term with the size indicating the ratio of the intersection between the term and the introgressed genes. The *y*-axis shows the -log_10_*P*_adj._ value up to a maximum of 16 and the panels along the *y*-axis and colours indicate the module. The panels along the *x*-axis indicate the source of the term and the position within the panels groups terms from the same GO subtree. The top driver GO terms up to a maximum of ten are indicated with a black outline and label.

**Fig S30.** g:Profiler functional enrichment of the genes in the base network with no background data set specified. Each dot represents a significantly enriched GO term with the size indicating the ratio of the intersection between the term and the introgressed genes. The *y*-axis shows the - log_10_*P*_adj._ value up to a maximum of 16 and the panels along the *y*-axis and colours indicate the module. The panels along the *x*-axis indicate the source of the term and the position within the panels groups terms from the same GO subtree. The top driver GO terms up to a maximum of ten are indicated with a black outline and label.

