## Supporting information (supplemental tables and figures) for "Integrative genomics identifies candidate genes underlying trypanotolerance in hybrid African cattle"

**Short title:** Integrative genomics highlights candidate trypanotolerance genes

**Authors and Affiliations:** Gillian P. McHugo^1^, James A. Ward^1^, Said Ismael Ng’ang’a^2,3^, Laurent A.F. Frantz^2,3^, John A. Browne^1^, Michael Salter-Townshend^4^, Grace M. O’Gorman^5^, Kieran G. Meade^1,6,7^, Emmeline W. Hill^1^, Thomas J. Hall^1^, David E. MacHugh^1,6,7*^

^1^ UCD School of Agriculture and Food Science, University College Dublin, Belfield, Dublin, Ireland.

^2^ Palaeogenomics Group, Department of Veterinary Sciences, Ludwig Maximilian University, Munich, Germany.

^3^ School of Biological and Behavioural Sciences, Queen Mary University of London, London, UK.

^4^ UCD School of Mathematics and Statistics, University College Dublin, Dublin, Ireland.

^5^ UK Agri-Tech Centre, Innovation Centre, York Science Park, York, UK.

^6^ UCD Conway Institute of Biomolecular and Biomedical Research, University College Dublin, Dublin, Ireland.

^7^ UCD One Health Centre, University College Dublin, Dublin, Ireland.

* Corresponding author

**Table S1.** Numbers of SNPs with *z*-score ≥ 2.0 for mean European *B. taurus*, African *B. taurus* and *B. indicus* ancestry components for the six populations with gene expression data available across all autosomes. The numbers in brackets indicate the percentage of the total 29,869 SNPs in the data set.

| **Group** | **Population** | **European**  ***Bos taurus* ancestry** | **African**  ***Bos taurus* ancestry** | ***Bos indicus* ancestry** |
| --- | --- | --- | --- | --- |
| African *Bos taurus* | LAGU | 1,348 (4.51%) | 0 (0%) | 1,257 (4.21%) |
| African *Bos taurus* | BAOU | 1,359 (4.55%) | 0 (0%) | 1,190 (3.98%) |
| Trypanotolerant African hybrid | NDAM | 1,407 (4.71%) | 112 (0.37%) | 1,148 (3.84%) |
| Trypanotolerant African hybrid | BORG | 1,193 (3.99%) | 455 (1.52%) | 958 (3.21%) |
| Trypanosusceptible African hybrid | FULA | 1,251 (4.19%) | 754 (2.52%) | 654 (2.19%) |
| Trypanosusceptible African hybrid | BORA | 1,282 (4.29%) | 1,125 (3.77%) | 300(1.00%) |

**Table S2.** Numbers of genes within 1 Mb up- and downstream of SNPs with *z*-score ≥ 2.0 for mean European *B. taurus*, African *B. taurus* and *B. indicus* ancestry components for the six populations with gene expression data available across all autosomes. The numbers in brackets indicate the percentage of the total 34,080 genes in the data set.

| **Group** | **Population** | **European**  ***Bos taurus* ancestry** | **African**  ***Bos taurus* ancestry** | ***Bos indicus* ancestry** |
| --- | --- | --- | --- | --- |
| African *Bos taurus* | LAGU | 845 (2.48%) | 0 (0%) | 1,076 (3.16%) |
| African *Bos taurus* | BAOU | 947 (2.78%) | 0 (0%) | 1,060 (3.11%) |
| Trypanotolerant African hybrid | NDAM | 1,006 (2.95%) | 45 (0.13%) | 842 (2.47%) |
| Trypanotolerant African hybrid | BORG | 561 (1.65%) | 82 (0.24%) | 1,232 (3.62%) |
| Trypanosusceptible African hybrid | FULA | 943 (2.77%) | 171 (0.50%) | 516 (1.51%) |
| Trypanosusceptible African hybrid | BORA | 796 (2.34%) | 569 (1.67%) | 129 (0.38%) |

**Table S3.** Population, days post-infection (dpi) and the significantly differentially expressed genes with increased and decreased expression with gene symbols for the response contrasts.

| **Population** | **dpi** | **Significantly differentially expressed genes with**  **increased expression** |
| --- | --- | --- |
| NDAM | 20 | *PDK4, SLC11A1, CPT1A, DCTN5* |
| BAOU | 30 | *SRXN1, GNG11* |
| NDAM | 30 | *SLC11A1* |
| LAGU | 40 | *IL12RB2, SLC25A36, NFAT5, E2F3, PHF20L1, PTPN4, ZC3HAV1, PIGR, PICALM, TTBK2, RAB11FIP2, MYSM1, PDPR, LOC100366216, SLC38A4, ZBTB37, CD55, GPATCH2L, NT5C2, TET2* |
| NDAM | 40 | *FKBP5* |
| **Population** | **dpi** | **Significantly differentially expressed genes with decreased expression** |
| NDAM | 30 | *SNX9, LZTS3* |
| LAGU | 40 | *OXER1, NEIL2, FADS1, NAB2, ANXA6* |
| BAOU | 40 | *SPARC* |
| NDAM | 40 | *GATA1, IGF2, STK36, PRKCG, NEIL2, SYNGR3, PDLIM1, LZTS3* |
| BORG | 40 | *NEIL2* |

**Table S4.** Gene symbol and modules containing each of the 243 genes with valid gene symbols that were found in four or more of the eight functional modules.

| **Symbol** | **Modules** |
| --- | --- |
| *AGO2* | MICRO BL 34, MICRO LI 35, MICRO LN 35, MICRO SP 35, RNA LAGU 40, RNA BAOU 40, RNA NDAM 40, RNA BORG 40 |
| *CBL* | MICRO BL 34, MICRO LI 35, MICRO LN 35, MICRO SP 35, RNA LAGU 40, RNA BAOU 40, RNA NDAM 40, RNA BORG 40 |
| *CNOT1* | MICRO BL 34, MICRO LI 35, MICRO LN 35, MICRO SP 35, RNA LAGU 40, RNA BAOU 40, RNA NDAM 40, RNA BORG 40 |
| *EDN1* | MICRO BL 34, MICRO LI 35, MICRO LN 35, MICRO SP 35, RNA LAGU 40, RNA BAOU 40, RNA NDAM 40, RNA BORG 40 |
| *IL1B* | MICRO BL 34, MICRO LI 35, MICRO LN 35, MICRO SP 35, RNA LAGU 40, RNA BAOU 40, RNA NDAM 40, RNA BORG 40 |
| *NFKB1* | MICRO BL 34, MICRO LI 35, MICRO LN 35, MICRO SP 35, RNA LAGU 40, RNA BAOU 40, RNA NDAM 40, RNA BORG 40 |
| *RIPK1* | MICRO BL 34, MICRO LI 35, MICRO LN 35, MICRO SP 35, RNA LAGU 40, RNA BAOU 40, RNA NDAM 40, RNA BORG 40 |
| *TRAF2* | MICRO BL 34, MICRO LI 35, MICRO LN 35, MICRO SP 35, RNA LAGU 40, RNA BAOU 40, RNA NDAM 40, RNA BORG 40 |
| *BIKBA* | MICRO BL 34, MICRO LI 35, MICRO LN 35, MICRO SP 35, RNA BAOU 40, RNA NDAM 40, RNA BORG 40 |
| *CAV3* | MICRO BL 34, MICRO LI 35, MICRO LN 35, MICRO SP 35, RNA BAOU 40, RNA NDAM 40, RNA BORG 40 |
| *CCND1* | MICRO BL 34, MICRO LI 35, MICRO LN 35, MICRO SP 35, RNA BAOU 40, RNA NDAM 40, RNA BORG 40 |
| *CFTR* | MICRO LI 35, MICRO LN 35, MICRO SP 35, RNA LAGU 40, RNA BAOU 40, RNA NDAM 40, RNA BORG 40 |
| *CTNNB1* | MICRO LI 35, MICRO LN 35, MICRO SP 35, RNA LAGU 40, RNA BAOU 40, RNA NDAM 40, RNA BORG 40 |
| *HPRT1* | MICRO BL 34, MICRO LI 35, MICRO LN 35, MICRO SP 35, RNA BAOU 40, RNA NDAM 40, RNA BORG 40 |
| *IKBKB* | MICRO BL 34, MICRO LI 35, MICRO LN 35, MICRO SP 35, RNA BAOU 40, RNA NDAM 40, RNA BORG 40 |
| *IL2RB* | MICRO BL 34, MICRO LI 35, MICRO SP 35, RNA LAGU 40, RNA BAOU 40, RNA NDAM 40, RNA BORG 40 |
| *IQCB1* | MICRO BL 34, MICRO LI 35, MICRO LN 35, MICRO SP 35, RNA LAGU 40, RNA NDAM 40, RNA BORG 40 |
| *JUN* | MICRO BL 34, MICRO LI 35, MICRO LN 35, MICRO SP 35, RNA LAGU 40, RNA BAOU 40, RNA BORG 40 |
| *LAMP2* | MICRO BL 34, MICRO LI 35, MICRO LN 35, MICRO SP 35, RNA LAGU 40, RNA BAOU 40, RNA NDAM 40 |
| *LBR* | MICRO BL 34, MICRO LI 35, MICRO SP 35, RNA LAGU 40, RNA BAOU 40, RNA NDAM 40, RNA BORG 40 |
| *MAPK10* | MICRO BL 34, MICRO LI 35, MICRO LN 35, MICRO SP 35, RNA BAOU 40, RNA NDAM 40, RNA BORG 40 |
| *MYD88* | MICRO BL 34, MICRO LI 35, MICRO LN 35, MICRO SP 35, RNA LAGU 40, RNA BAOU 40, RNA BORG 40 |
| *PARP1* | MICRO BL 34, MICRO LI 35, MICRO LN 35, MICRO SP 35, RNA LAGU 40, RNA NDAM 40, RNA BORG 40 |
| *PIK3R1* | MICRO LI 35, MICRO LN 35, MICRO SP 35, RNA LAGU 40, RNA BAOU 40, RNA NDAM 40, RNA BORG 40 |
| *PKD1* | MICRO BL 34, MICRO LI 35, MICRO LN 35, MICRO SP 35, RNA LAGU 40, RNA BAOU 40, RNA NDAM 40 |
| *REL* | MICRO LI 35, MICRO LN 35, MICRO SP 35, RNA LAGU 40, RNA BAOU 40, RNA NDAM 40, RNA BORG 40 |
| *SELE* | MICRO BL 34, MICRO LN 35, MICRO SP 35, RNA LAGU 40, RNA BAOU 40, RNA NDAM 40, RNA BORG 40 |
| *SKP1* | MICRO BL 34, MICRO LI 35, MICRO LN 35, MICRO SP 35, RNA LAGU 40, RNA BAOU 40, RNA BORG 40 |
| *TLR2* | MICRO BL 34, MICRO LI 35, MICRO LN 35, MICRO SP 35, RNA LAGU 40, RNA BAOU 40, RNA NDAM 40 |
| *TP53* | MICRO BL 34, MICRO LN 35, MICRO SP 35, RNA LAGU 40, RNA BAOU 40, RNA NDAM 40, RNA BORG 40 |
| *ALB* | MICRO BL 34, MICRO LI 35, MICRO LN 35, RNA BAOU 40, RNA NDAM 40, RNA BORG 40 |
| *APOE* | MICRO LI 35, MICRO LN 35, MICRO SP 35, RNA BAOU 40, RNA NDAM 40, RNA BORG 40 |
| *CCND2* | MICRO BL 34, MICRO LN 35, MICRO SP 35, RNA LAGU 40, RNA NDAM 40, RNA BORG 40 |
| *CDC37* | MICRO LN 35, MICRO SP 35, RNA LAGU 40, RNA BAOU 40, RNA NDAM 40, RNA BORG 40 |
| *CDKN1A* | MICRO LN 35, MICRO SP 35, RNA LAGU 40, RNA BAOU 40, RNA NDAM 40, RNA BORG 40 |
| *COPS6* | MICRO BL 34, MICRO LI 35, MICRO SP 35, RNA LAGU 40, RNA BAOU 40, RNA NDAM 40 |
| *CTR9* | MICRO LI 35, MICRO LN 35, MICRO SP 35, RNA LAGU 40, RNA BAOU 40, RNA NDAM 40 |
| *ERBB3* | MICRO LI 35, MICRO LN 35, MICRO SP 35, RNA BAOU 40, RNA NDAM 40, RNA BORG 40 |
| *GRIN1* | MICRO BL 34, MICRO LI 35, MICRO LN 35, MICRO SP 35, RNA NDAM 40, RNA BORG 40 |
| *HRMT1L2* | MICRO LI 35, MICRO SP 35, RNA LAGU 40, RNA BAOU 40, RNA NDAM 40, RNA BORG 40 |
| *IL1R2* | MICRO LI 35, MICRO SP 35, RNA LAGU 40, RNA BAOU 40, RNA NDAM 40, RNA BORG 40 |
| *IL2RA* | MICRO BL 34, MICRO SP 35, RNA LAGU 40, RNA BAOU 40, RNA NDAM 40, RNA BORG 40 |
| *IRAK1* | MICRO BL 34, MICRO LI 35, MICRO LN 35, MICRO SP 35, RNA LAGU 40, RNA NDAM 40 |
| *KRT18* | MICRO BL 34, MICRO LI 35, MICRO LN 35, RNA BAOU 40, RNA NDAM 40, RNA BORG 40 |
| *MT-CO2* | MICRO BL 34, MICRO LI 35, MICRO LN 35, MICRO SP 35, RNA NDAM 40, RNA BORG 40 |
| *NEK2* | MICRO BL 34, MICRO LI 35, MICRO LN 35, MICRO SP 35, RNA LAGU 40, RNA NDAM 40 |
| *NTRK2* | MICRO LI 35, MICRO LN 35, MICRO SP 35, RNA BAOU 40, RNA NDAM 40,RNA BORG 40 |
| *PLCG1* | MICRO BL 34, MICRO LI 35, MICRO LN 35, MICRO SP 35, RNA LAGU 40, RNA BORG 40 |
| *POLR2A* | MICRO BL 34, MICRO LI 35, NA, MICRO SP 35, RNA BAOU 40, RNA NDAM 40, RNA BORG 40 |
| *RFC1* | MICRO BL 34, MICRO LN 35, MICRO SP 35, RNA BAOU 40, RNA NDAM 40, RNA BORG 40 |
| *SF3B14* | MICRO BL 34, MICRO LN 35, MICRO SP 35, RNA BAOU 40, RNA NDAM 40, RNA BORG 40 |
| *SIRT1* | MICRO LN 35, MICRO SP 35, RNA LAGU 40, RNA BAOU 40, RNA NDAM 40, RNA BORG 40 |
| *SLC11A1* | MICRO LI 35, MICRO LN 35, MICRO SP 35, RNA LAGU 40, RNA NDAM 40, RNA BORG 40 |
| *TBK1* | MICRO LI 35, MICRO LN 35, MICRO SP 35, RNA LAGU 40, RNA BAOU 40, RNA NDAM 40 |
| *TERF1* | MICRO LI 35, MICRO SP 35, RNA LAGU 40, RNA BAOU 40, RNA NDAM 40, RNA BORG 40 |
| *TOP1* | MICRO LI 35, MICRO LN 35, MICRO SP 35, RNA BAOU 40, RNA NDAM 40, RNA BORG 40 |
| *TRADD* | MICRO BL 34, MICRO LI 35, MICRO LN 35, MICRO SP 35, RNA BAOU 40, RNA NDAM 40 |
| *ACTC1* | MICRO SP 35, RNA LAGU 40, RNA BAOU 40, RNA NDAM 40, RNA BORG 40 |
| *ACTN1* | MICRO BL 34, MICRO LN 35, RNA LAGU 40, RNA NDAM 40, RNA BORG 40 |
| *AR* | MICRO LN 35, MICRO SP 35, RNA BAOU 40, RNA NDAM 40, RNA BORG 40 |
| *ARF1* | MICRO LI 35, MICRO LN 35, RNA LAGU 40, RNA NDAM 40, RNA BORG 40 |
| *ARRB1* | MICRO BL 34, MICRO LI 35, MICRO LN 35, MICRO SP 35, RNA LAGU 40 |
| *ATF2* | MICRO LN 35, RNA LAGU 40, RNA BAOU 40, RNA NDAM 40, RNA BORG 40 |
| *CD14* | MICRO BL 34, MICRO LN 35, MICRO SP 35, RNA LAGU 40, RNA NDAM 40 |
| *CDK4* | MICRO BL 34, MICRO LI 35, MICRO LN 35, MICRO SP 35, RNA NDAM 40 |
| *CDK9* | MICRO LN 35, MICRO SP 35, RNA LAGU 40, RNA NDAM 40, RNA BORG 40 |
| *CHUK* | MICRO LN 35, RNA LAGU 40, RNA BAOU 40, RNA NDAM 40, RNA BORG 40 |
| *CREBBP* | MICRO BL 34, MICRO LI 35, MICRO LN 35, MICRO SP 35, RNA LAGU 40 |
| *CSF3* | MICRO SP 35, RNA LAGU 40, RNA BAOU 40, RNA NDAM 40, RNA BORG 40 |
| *CSNK2A1* | MICRO LI 35, MICRO SP 35, RNA BAOU 40, RNA NDAM 40, RNA BORG 40 |
| *CTTN* | MICRO LI 35, MICRO LN 35, MICRO SP 35, RNA BAOU 40, RNA NDAM 40 |
| *CUL2* | MICRO LI 35, MICRO SP 35, RNA LAGU 40, RNA BAOU 40, RNA BORG 40 |
| *CXCL10* | MICRO LN 35, MICRO SP 35, RNA LAGU 40, RNA BAOU 40, RNA NDAM 40 |
| *DDX58* | MICRO SP 35, RNA LAGU 40, RNA BAOU 40, RNA NDAM 40, RNA BORG 40 |
| *DLG4* | MICRO LI 35, MICRO LN 35, MICRO SP 35, RNA LAGU 40, RNA NDAM 40 |
| *DST* | MICRO LI 35, MICRO SP 35, RNA BAOU 40, RNA NDAM 40, RNA BORG 40 |
| *EEF1B* | MICRO SP 35, RNA LAGU 40, RNA BAOU 40, RNA NDAM 40, RNA BORG 40 |
| *EIF2C1* | MICRO SP 35, RNA LAGU 40, RNA BAOU 40, RNA NDAM 40, RNA BORG 40 |
| *EIF2S1* | MICRO LI 35, MICRO LN 35, MICRO SP 35, RNA LAGU 40, RNA NDAM 40 |
| *FBXW11* | MICRO LN 35, MICRO SP 35, RNA LAGU 40, RNA BAOU 40, RNA NDAM 40 |
| *FLIP* | MICRO SP 35, RNA LAGU 40, RNA BAOU 40, RNA NDAM 40, RNA BORG 40 |
| *GRK5* | MICRO LN 35, MICRO SP 35, RNA LAGU 40, RNA NDAM 40, RNA BORG 40 |
| *HNRNPD* | MICRO BL 34, MICRO LI 35, RNA BAOU 40, RNA NDAM 40, RNA BORG 40 |
| *HSP90AA1* | MICRO BL 34, MICRO LI 35, MICRO LN 35, MICRO SP 35, RNA LAGU 40 |
| *HSPA9* | MICRO LN 35, MICRO SP 35, RNA LAGU 40, RNA NDAM 40, RNA BORG 40 |
| *IL12RB2* | MICRO SP 35, RNA LAGU 40, RNA BAOU 40, RNA NDAM 40, RNA BORG 40 |
| *IL17A* | MICRO SP 35, RNA LAGU 40, RNA BAOU 40, RNA NDAM 40, RNA BORG 40 |
| *IRF1* | MICRO BL 34, MICRO LI 35, MICRO SP 35, RNA BAOU 40, RNA BORG 40 |
| *IRF3* | MICRO BL 34, MICRO SP 35, RNA LAGU 40, RNA NDAM 40, RNA BORG 40 |
| *IRF4* | MICRO LI 35, MICRO LN 35, RNA BAOU 40, RNA NDAM 40, RNA BORG 40 |
| *LRRK2* | MICRO LI 35, MICRO SP 35, RNA BAOU 40, RNA NDAM 40, RNA BORG 40 |
| *LYN* | MICRO LN 35, MICRO SP 35, RNA LAGU 40, RNA BAOU 40, RNA NDAM 40 |
| *MAP3K7* | MICRO LN 35, MICRO SP 35, RNA LAGU 40, RNA BAOU 40, RNA NDAM 40 |
| *MCL1* | MICRO BL 34, MICRO LI 35, RNA BAOU 40, RNA NDAM 40, RNA BORG 40 |
| *MDM2* | MICRO BL 34, MICRO SP 35, RNA LAGU 40, RNA NDAM 40, RNA BORG 40 |
| *METTL21B* | MICRO LI 35, MICRO SP 35, RNA BAOU 40, RNA NDAM 40, RNA BORG 40 |
| *MT-ND4* | MICRO BL 34, MICRO LI 35, MICRO LN 35, MICRO SP 35, RNA LAGU 40 |
| *NCOA6* | MICRO BL 34, MICRO LN 35, MICRO SP 35, RNA LAGU 40, RNA NDAM 40 |
| *NGF* | MICRO LN 35, MICRO SP 35, RNA BAOU 40, RNA NDAM 40, RNA BORG 40 |
| *NPHP1* | MICRO BL 34, MICRO LI 35, MICRO LN 35, MICRO SP 35, RNA BAOU 40 |
| *NPHP4* | MICRO BL 34, MICRO LN 35, MICRO SP 35, RNA BAOU 40, RNA BORG 40 |
| *PLAA* | MICRO BL 34, MICRO LI 35, MICRO LN 35, MICRO SP 35, RNA NDAM 40 |
| *PML* | MICRO BL 34, MICRO SP 35, RNA BAOU 40, RNA NDAM 40, RNA BORG 40 |
| *PPP2R1A* | MICRO BL 34, MICRO LI 35, MICRO LN 35, MICRO SP 35, RNA LAGU 40 |
| *PPP2R4* | MICRO LI 35, MICRO SP 35, RNA BAOU 40, RNA NDAM 40, RNA BORG 40 |
| *PSMD10* | MICRO LI 35, MICRO LN 35, RNA BAOU 40, RNA NDAM 40, RNA BORG 40 |
| *PTK2* | MICRO BL 34, MICRO LI 35, MICRO LN 35, RNA LAGU 40, RNA BORG 40 |
| *PTPN11* | MICRO LI 35, MICRO LN 35, MICRO SP 35, RNA BAOU 40, RNA NDAM 40 |
| *RAB11A* | MICRO BL 34, MICRO LI 35, MICRO LN 35, MICRO SP 35, RNA NDAM 40 |
| *RELB* | MICRO BL 34, MICRO LI 35, MICRO SP 35, RNA BAOU 40, RNA NDAM 40 |
| *RRM2* | MICRO BL 34, MICRO LN 35, MICRO SP 35, RNA LAGU 40, RNA NDAM 40 |
| *SF1* | MICRO BL 34, MICRO LN 35, RNA LAGU 40, RNA NDAM 40, RNA BORG 40 |
| *SF3A2* | MICRO BL 34, MICRO LI 35, RNA BAOU 40, RNA NDAM 40, RNA BORG 40 |
| *SHC1* | MICRO BL 34, MICRO LN 35, RNA BAOU 40, RNA NDAM 40, RNA BORG 40 |
| *SMARCA4* | MICRO BL 34, MICRO LI 35, MICRO LN 35, RNA LAGU 40, RNA BORG 40 |
| *SP1* | MICRO BL 34, MICRO LI 35, MICRO LN 35, MICRO SP 35, RNA BAOU 40 |
| *STAT3* | MICRO BL 34, MICRO SP 35, RNA LAGU 40, RNA BAOU 40, RNA NDAM 40 |
| *SYNCRIP* | MICRO BL 34, MICRO LI 35, MICRO LN 35, RNA BAOU 40, RNA NDAM 40 |
| *TCN2* | MICRO BL 34, MICRO LI 35, MICRO LN 35, MICRO SP 35, RNA LAGU 40 |
| *TCP1* | MICRO BL 34, MICRO LI 35, MICRO SP 35, RNA BAOU 40, RNA NDAM 40 |
| *TRAF6* | MICRO LI 35, MICRO LN 35, MICRO SP 35, RNA NDAM 40, RNA BORG 40 |
| *TRIM21* | MICRO BL 34, MICRO SP 35, RNA LAGU 40, RNA BAOU 40, RNA NDAM 40 |
| *TUBB5* | MICRO SP 35, RNA LAGU 40, RNA BAOU 40, RNA NDAM 40, RNA BORG 40 |
| *TXNRD1* | MICRO BL 34, MICRO LI 35, MICRO LN 35, MICRO SP 35, RNA NDAM 40 |
| *XIAP* | MICRO LN 35, MICRO SP 35, RNA LAGU 40, RNA BAOU 40, RNA NDAM 40 |
| *YWHAE* | MICRO BL 34, MICRO SP 35, RNA BAOU 40, RNA NDAM 40, RNA BORG 40 |
| *A2M* | MICRO LN 35, MICRO SP 35, RNA LAGU 40, RNA BAOU 40 |
| *AHCY* | MICRO LN 35, MICRO SP 35, RNA LAGU 40, RNA NDAM 40 |
| *AKAP8* | MICRO BL 34, MICRO LI 35, MICRO SP 35, RNA NDAM 40 |
| *APAF1* | MICRO LN 35, MICRO SP 35, RNA LAGU 40, RNA NDAM 40 |
| *APLP2* | MICRO BL 34, MICRO LI 35, MICRO LN 35, MICRO SP 35 |
| *APOA2* | MICRO LI 35, RNA BAOU 40, RNA NDAM 40, RNA BORG 40 |
| *APP* | MICRO BL 34, MICRO LN 35, RNA LAGU 40, RNA NDAM 40 |
| *ASPM* | MICRO BL 34, MICRO LN 35, MICRO SP 35, RNA NDAM 40 |
| *ATF3* | MICRO BL 34, MICRO LI 35, MICRO SP 35, RNA NDAM 40 |
| *ATP5J* | MICRO SP 35, RNA BAOU 40, RNA NDAM 40, RNA BORG 40 |
| *ATR* | MICRO LN 35, MICRO SP 35, RNA LAGU 40, RNA NDAM 40 |
| *BCAR3* | MICRO LN 35, MICRO SP 35, RNA LAGU 40, RNA NDAM 40 |
| *BCR* | MICRO BL 34, MICRO LN 35, MICRO SP 35, RNA LAGU 40 |
| *BDNF* | MICRO SP 35, RNA BAOU 40, RNA NDAM 40, RNA BORG 40 |
| *BECN1* | MICRO LI 35, MICRO LN 35, MICRO SP 35, RNA LAGU 40 |
| *BIRC3* | MICRO LN 35, MICRO SP 35, RNA BAOU 40, RNA NDAM 40 |
| *C20ORF18* | MICRO SP 35, RNA BAOU 40, RNA NDAM 40, RNA BORG 40 |
| *C2ORF29* | MICRO SP 35, RNA BAOU 40, RNA NDAM 40, RNA BORG 40 |
| *C3H1orf52* | MICRO BL 34, RNA BAOU 40, RNA NDAM 40, RNA BORG 40 |
| *CCR5* | MICRO BL 34, MICRO LN 35, MICRO SP 35, RNA LAGU 40 |
| *CCRL1* | MICRO SP 35, RNA BAOU 40, RNA NDAM 40, RNA BORG 40 |
| *CD28* | MICRO BL 34, MICRO LN 35, MICRO SP 35, RNA LAGU 40 |
| *CDC25C* | MICRO BL 34, MICRO SP 35, RNA LAGU 40, RNA NDAM 40 |
| *CDC73* | MICRO BL 34, MICRO LI 35, RNA LAGU 40, RNA BAOU 40 |
| *CDK1* | MICRO LN 35, RNA BAOU 40, RNA NDAM 40, RNA BORG 40 |
| *CLU* | MICRO BL 34, MICRO SP 35, RNA LAGU 40, RNA NDAM 40 |
| *CMYA5* | MICRO SP 35, RNA BAOU 40, RNA NDAM 40, RNA BORG 40 |
| *CNOT6* | MICRO LN 35, MICRO SP 35, RNA LAGU 40, RNA BAOU 40 |
| *CNOT7* | MICRO BL 34, MICRO LI 35, MICRO SP 35, RNA BORG 40 |
| *CNTF* | MICRO SP 35, RNA BAOU 40, RNA NDAM 40, RNA BORG 40 |
| *CRYAB* | MICRO LI 35, RNA BAOU 40, RNA NDAM 40, RNA BORG 40 |
| *CSF2* | MICRO SP 35, RNA BAOU 40, RNA NDAM 40, RNA BORG 40 |
| *CSNK1G1* | MICRO SP 35, RNA LAGU 40, RNA BAOU 40, RNA NDAM 40 |
| *CSNK2A2* | MICRO BL 34, MICRO SP 35, RNA LAGU 40, RNA NDAM 40 |
| *CTPS* | MICRO SP 35, RNA BAOU 40, RNA NDAM 40, RNA BORG 40 |
| *CXCR7* | MICRO SP 35, RNA BAOU 40, RNA NDAM 40, RNA BORG 40 |
| *DDIT4* | MICRO LI 35, RNA LAGU 40, RNA NDAM 40, RNA BORG 40 |
| *DHFR* | MICRO BL 34, MICRO LI 35, MICRO LN 35, MICRO SP 35 |
| *DPF1* | MICRO SP 35, RNA BAOU 40, RNA NDAM 40, RNA BORG 40 |
| *DVL2* | MICRO BL 34, MICRO LI 35, MICRO SP 35, RNA NDAM 40 |
| *EIF2AK2* | MICRO LI 35, MICRO SP 35, RNA LAGU 40, RNA NDAM 40 |
| *ELA2* | MICRO SP 35, RNA BAOU 40, RNA NDAM 40, RNA BORG 40 |
| *EP300* | MICRO SP 35, RNA BAOU 40, RNA NDAM 40, RNA BORG 40 |
| *FBXO32* | MICRO LN 35, RNA LAGU 40, RNA BAOU 40, RNA BORG 40 |
| *FBXO5* | MICRO LN 35, MICRO SP 35, RNA BAOU 40, RNA BORG 40 |
| *FKBP5* | MICRO SP 35, RNA LAGU 40, RNA NDAM 40, RNA BORG 40 |
| *FLJ20565* | MICRO SP 35, RNA BAOU 40, RNA NDAM 40, RNA BORG 40 |
| *FN1* | MICRO LN 35, RNA BAOU 40, RNA NDAM 40, RNA BORG 40 |
| *GEMIN2* | MICRO BL 34, MICRO LI 35, MICRO LN 35, MICRO SP 35 |
| *GNB2L1* | MICRO SP 35, RNA BAOU 40, RNA NDAM 40, RNA BORG 40 |
| *GR-A* | MICRO SP 35, RNA BAOU 40, RNA NDAM 40, RNA BORG 40 |
| *GRB2* | MICRO SP 35, RNA LAGU 40, RNA BAOU 40, RNA NDAM 40 |
| *HGS* | MICRO LI 35, MICRO SP 35, RNA LAGU 40, RNA NDAM 40 |
| *HMGA2* | MICRO SP 35, RNA BAOU 40, RNA NDAM 40, RNA BORG 40 |
| *HMGB1* | MICRO BL 34, MICRO LI 35, MICRO LN 35, MICRO SP 35 |
| *HSPA4* | MICRO SP 35, RNA BAOU 40, RNA NDAM 40, RNA BORG 40 |
| *HTT* | MICRO BL 34, MICRO LN 35, MICRO SP 35, RNA BORG 40 |
| *ICT1* | MICRO SP 35, RNA BAOU 40, RNA NDAM 40, RNA BORG 40 |
| *IL10* | MICRO LN 35, MICRO SP 35, RNA LAGU 40, RNA NDAM 40 |
| *IL11* | MICRO SP 35, RNA BAOU 40, RNA NDAM 40, RNA BORG 40 |
| *IL12A* | MICRO SP 35, RNA BAOU 40, RNA NDAM 40, RNA BORG 40 |
| *ILF3* | MICRO BL 34, MICRO LN 35, MICRO SP 35, RNA NDAM 40 |
| *IRF2* | MICRO BL 34, MICRO LI 35, MICRO LN 35, MICRO SP 35 |
| *ISG15* | MICRO BL 34, MICRO LN 35, MICRO SP 35, RNA LAGU 40 |
| *ITGB2* | MICRO LN 35, MICRO SP 35, RNA LAGU 40, RNA NDAM 40 |
| *JAKMIP1* | MICRO SP 35, RNA BAOU 40, RNA NDAM 40, RNA BORG 40 |
| *KAT5* | MICRO BL 34, MICRO LN 35, RNA LAGU 40, RNA BORG 40 |
| *KSR1* | MICRO LN 35, MICRO SP 35, RNA LAGU 40, RNA NDAM 40 |
| *LARS* | MICRO LI 35, MICRO LN 35, MICRO SP 35, RNA NDAM 40 |
| *LCK* | MICRO LN 35, MICRO SP 35, RNA BAOU 40, RNA NDAM 40 |
| *LEO1* | MICRO SP 35, RNA LAGU 40, RNA BAOU 40, RNA NDAM 40 |
| *LIFR* | MICRO SP 35, RNA BAOU 40, RNA NDAM 40, RNA BORG 40 |
| *MAFG* | MICRO BL 34, MICRO LI 35, MICRO LN 35, MICRO SP 35 |
| *MAP3K7IP1* | MICRO BL 34, MICRO SP 35, RNA BAOU 40, RNA NDAM 40 |
| *MAPK6* | MICRO SP 35, RNA LAGU 40, RNA BAOU 40, RNA NDAM 40 |
| *MDH2* | MICRO LI 35, RNA BAOU 40, RNA NDAM 40, RNA BORG 40 |
| *MIB1* | MICRO SP 35, RNA LAGU 40, RNA BAOU 40, RNA NDAM 40 |
| *NANOG* | MICRO SP 35, RNA BAOU 40, RNA NDAM 40, RNA BORG 40 |
| *NANOS2* | MICRO SP 35, RNA BAOU 40, RNA NDAM 40, RNA BORG 40 |
| *NFAT5* | MICRO SP 35, RNA LAGU 40, RNA BAOU 40, RNA BORG 40 |
| *NFATC2* | MICRO BL 34, MICRO LI 35, MICRO LN 35, MICRO SP 35 |
| *NFKBIE* | MICRO BL 34, MICRO SP 35, RNA LAGU 40, RNA BAOU 40 |
| *NFKBIL1* | MICRO BL 34, MICRO LI 35, MICRO LN 35, MICRO SP 35 |
| *NPM1* | MICRO LI 35, MICRO LN 35, MICRO SP 35, RNA BAOU 40 |
| *NR4A1* | MICRO BL 34, MICRO LI 35, MICRO SP 35, RNA NDAM 40 |
| *PCAF* | MICRO SP 35, RNA BAOU 40, RNA NDAM 40, RNA BORG 40 |
| *PGD* | MICRO BL 34, MICRO LI 35, RNA LAGU 40, RNA NDAM 40 |
| *PGHS-2* | MICRO SP 35, RNA BAOU 40, RNA NDAM 40, RNA BORG 40 |
| *PGR* | MICRO SP 35, RNA BAOU 40, RNA NDAM 40, RNA BORG 40 |
| *PIK3R2* | MICRO BL 34, MICRO SP 35, RNA LAGU 40, RNA NDAM 40 |
| *POU2F1* | MICRO BL 34, MICRO LI 35, MICRO LN 35, RNA LAGU 40 |
| *PPP2R1B* | MICRO LI 35, MICRO LN 35, RNA LAGU 40, RNA NDAM 40 |
| *PRDX5* | MICRO BL 34, MICRO LI 35, MICRO LN 35, RNA LAGU 40 |
| *PRKCI* | MICRO BL 34, MICRO LN 35, MICRO SP 35, RNA LAGU 40 |
| *PROS1* | MICRO LN 35, MICRO SP 35, RNA BAOU 40, RNA NDAM 40 |
| *PTPN1* | MICRO LN 35, MICRO SP 35, RNA LAGU 40, RNA NDAM 40 |
| *PTPN6* | MICRO BL 34, MICRO SP 35, RNA BAOU 40, RNA BORG 40 |
| *RAG1* | MICRO SP 35, RNA BAOU 40, RNA NDAM 40, RNA BORG 40 |
| *RAG2* | MICRO SP 35, RNA BAOU 40, RNA NDAM 40, RNA BORG 40 |
| *RGS7* | MICRO SP 35, RNA BAOU 40, RNA NDAM 40, RNA BORG 40 |
| *RQCD1* | MICRO SP 35, RNA BAOU 40, RNA NDAM 40, RNA BORG 40 |
| *RRM2B* | MICRO BL 34, MICRO LN 35, MICRO SP 35, RNA LAGU 40 |
| *SELL* | MICRO BL 34, MICRO LI 35, MICRO SP 35, RNA LAGU 40 |
| *SNCA* | MICRO BL 34, MICRO SP 35, RNA NDAM 40, RNA BORG 40 |
| *SRRM2* | MICRO BL 34, MICRO LI 35, RNA BAOU 40, RNA BORG 40 |
| *STAT5B* | MICRO BL 34, MICRO LI 35, RNA BAOU 40, RNA BORG 40 |
| *TBPL1* | MICRO LI 35, MICRO LN 35, MICRO SP 35, RNA NDAM 40 |
| *TGFB1* | MICRO BL 34, MICRO SP 35, RNA LAGU 40, RNA NDAM 40 |
| *TLR4* | MICRO BL 34, MICRO LI 35, MICRO SP 35, RNA LAGU 40 |
| *TMEM66* | MICRO SP 35, RNA BAOU 40, RNA NDAM 40, RNA BORG 40 |
| *TNF* | MICRO LI 35, MICRO SP 35, RNA LAGU 40, RNA BAOU 40 |
| *TNIP3* | MICRO SP 35, RNA LAGU 40, RNA BAOU 40, RNA NDAM 40 |
| *TOMM70A* | MICRO SP 35, RNA BAOU 40, RNA NDAM 40, RNA BORG 40 |
| *TRAIP* | MICRO SP 35, RNA BAOU 40, RNA NDAM 40, RNA BORG 40 |
| *UBA1* | MICRO BL 34, MICRO LI 35, MICRO LN 35, MICRO SP 35 |
| *UBE2D1* | MICRO BL 34, MICRO LI 35, MICRO SP 35, RNA LAGU 40 |
| *UBE2N* | MICRO LI 35, MICRO LN 35, MICRO SP 35, RNA BORG 40 |
| *UNC5CL* | MICRO SP 35, RNA BAOU 40, RNA NDAM 40, RNA BORG 40 |
| *VDR* | MICRO BL 34, MICRO LN 35, MICRO SP 35, RNA LAGU 40 |
| *VIM* | MICRO BL 34, RNA LAGU 40, RNA BAOU 40, RNA NDAM 40 |
| *VPS4A* | MICRO SP 35, RNA BAOU 40, RNA NDAM 40, RNA BORG 40 |
| *WDR96* | MICRO SP 35, RNA BAOU 40, RNA NDAM 40, RNA BORG 40 |
| *YY1* | MICRO BL 34, MICRO LI 35, MICRO SP 35, RNA BAOU 40 |

**Table S5.** Numbers of intervals within 1 Mb up- and downstream of SNPs with *z*-score ≥ 2.0 for mean European *B. taurus*, African *B. taurus* and *B. indicus* ancestry components for the six populations with gene expression data available across all autosomes and groups of populations from the original local ancestry analysis.

| **Analysis** | **Software** | **Data** | **Group/Population** | **European *Bos taurus* intervals** | **African**  ***Bos taurus* intervals** | ***Bos indicus* intervals** |
| --- | --- | --- | --- | --- | --- | --- |
| Updated | ELAI | LD | LAGU | 1,237 | 0 | 1,169 |
| Updated | ELAI | LD | BAOU | 1,262 | 0 | 1,111 |
| Updated | ELAI | LD | NDAM | 1,302 | 102 | 1,049 |
| Updated | ELAI | LD | BORG | 1,084 | 426 | 864 |
| Updated | ELAI | LD | FULA | 1,150 | 711 | 580 |
| Updated | ELAI | LD | BORA | 1,203 | 1,053 | 260 |
| Original | ELAI | HD | Selected residually admixed European | 0 | 9,296 | 12,898 |
| Original | ELAI | HD | Selected trypanotolerant African hybrids | 10,748 | 3,408 | 6,966 |
| Original | ELAI | HD | Trypanosusceptible African hybrids | 10,273 | 8,930 | 2,871 |
| Original | ELAI | LD | Selected residually admixed European | 132 | 1,220 | 1,306 |
| Original | ELAI | LD | Selected trypanotolerant African hybrids | 1,153 | 582 | 738 |
| Original | ELAI | LD | Trypanosusceptible African hybrids | 1,091 | 1,013 | 398 |
| Original | MOSAIC | HD | Selected residually admixed European | 0 | 6,737 | 12,802 |
| Original | MOSAIC | HD | Selected trypanotolerant African hybrids | 9,924 | 3,457 | 6,774 |
| Original | MOSAIC | HD | Trypanosusceptible African hybrids | 10,634 | 7,352 | 3,415 |

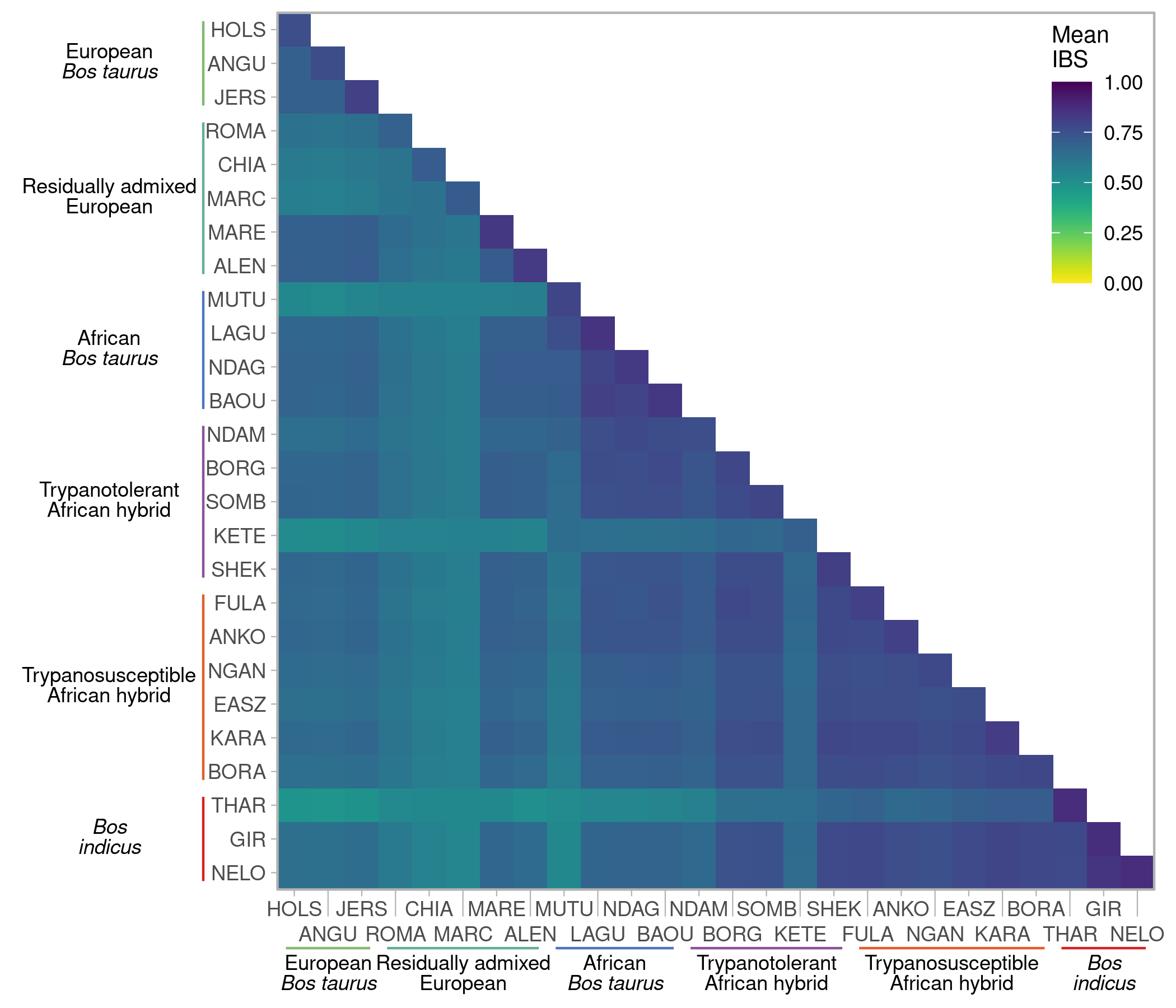

**Fig S1.** Heatmap of mean identity by state values for SNP data in European, African, and Asian cattle populations.

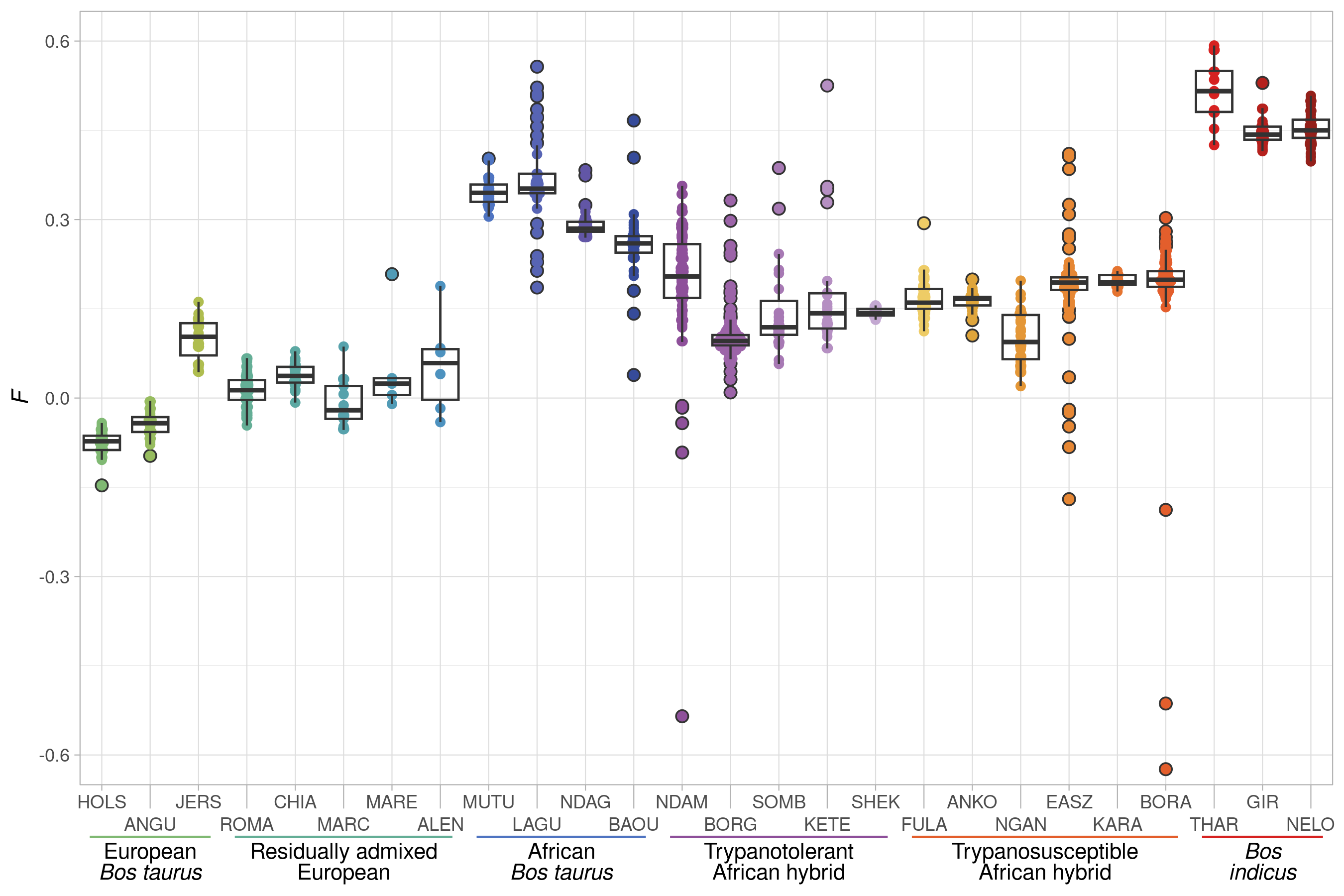

**Fig S2.** Tukey box plots showing the distribution of inbreeding values (*F*) for SNP data for each population of European, African, and Asian cattle. Outliers are indicated with a black outline.

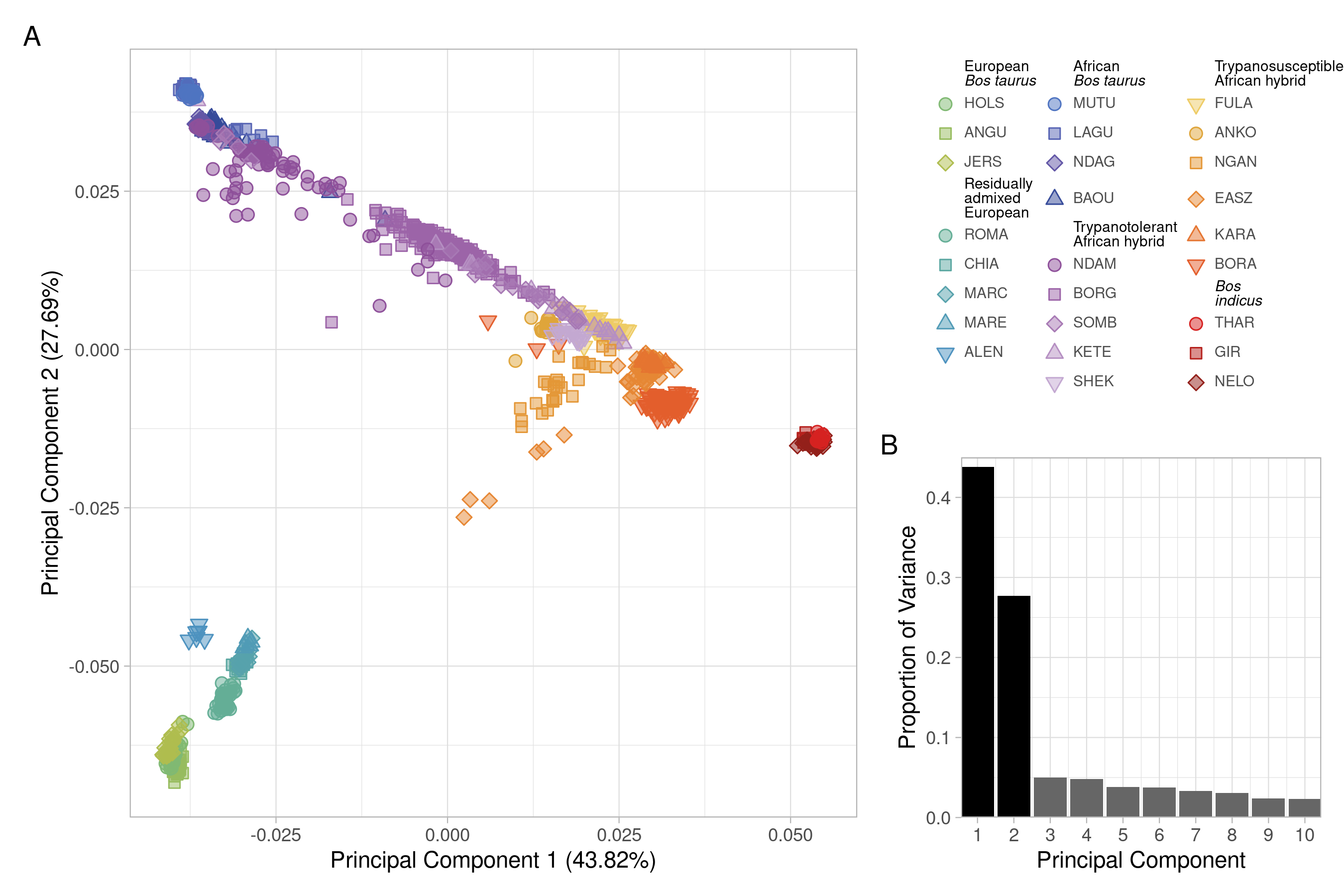

**Fig S3. A.** Principal component analysis (PCA) of SNP data for cattle coloured according to population showing the first two principal components and **B.** bar chart of proportion of variance of the top ten principal components.

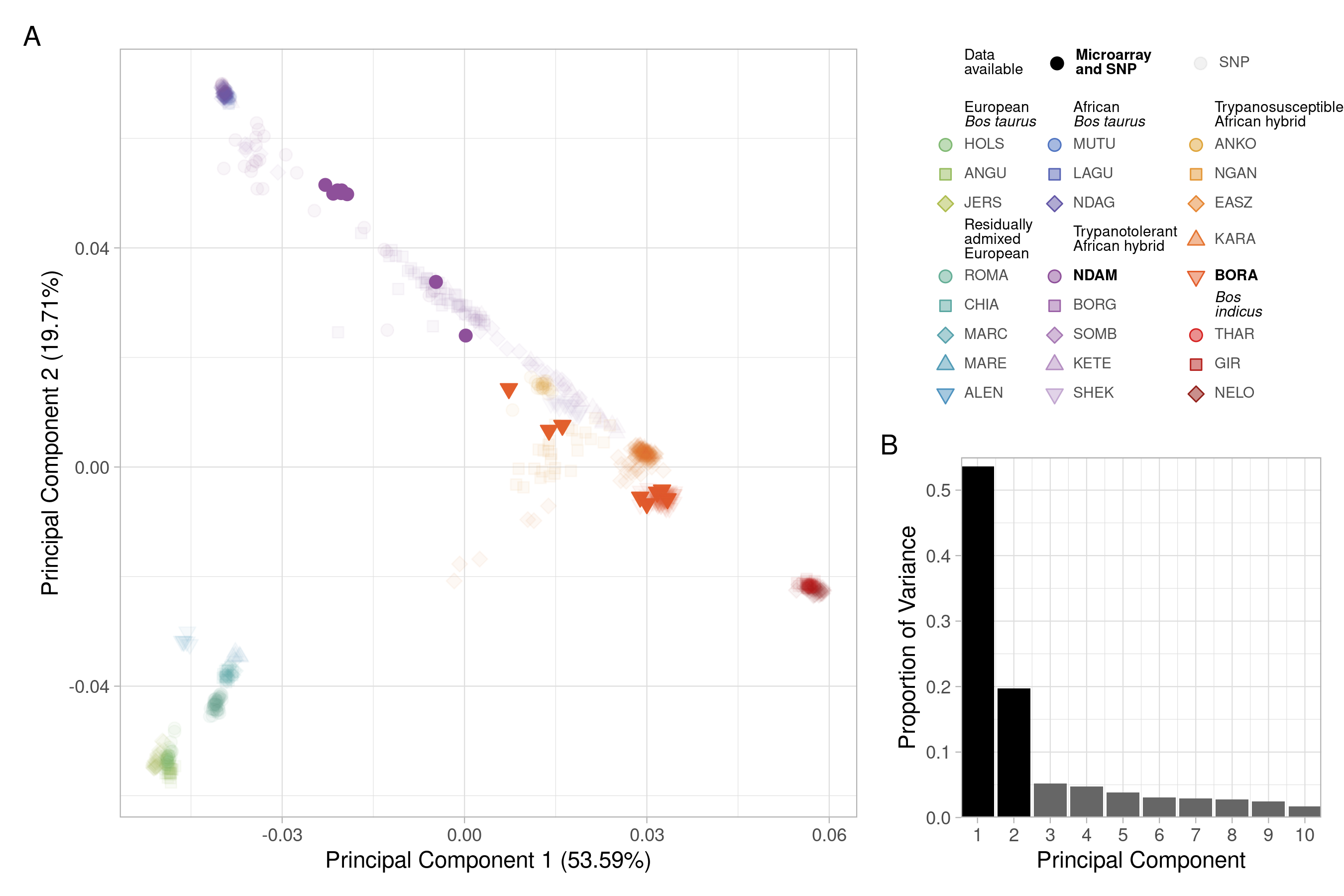

**Fig S4. A.** Principal component analysis (PCA) of the high-density SNP data for the cattle samples from McHugo and colleagues (McHugo *et al.* 2025) coloured according to population showing the first two principal components and **B.** bar chart of proportion of variance of the top ten principal components. The transparency indicates the availability of microarray gene expression data for the sample.

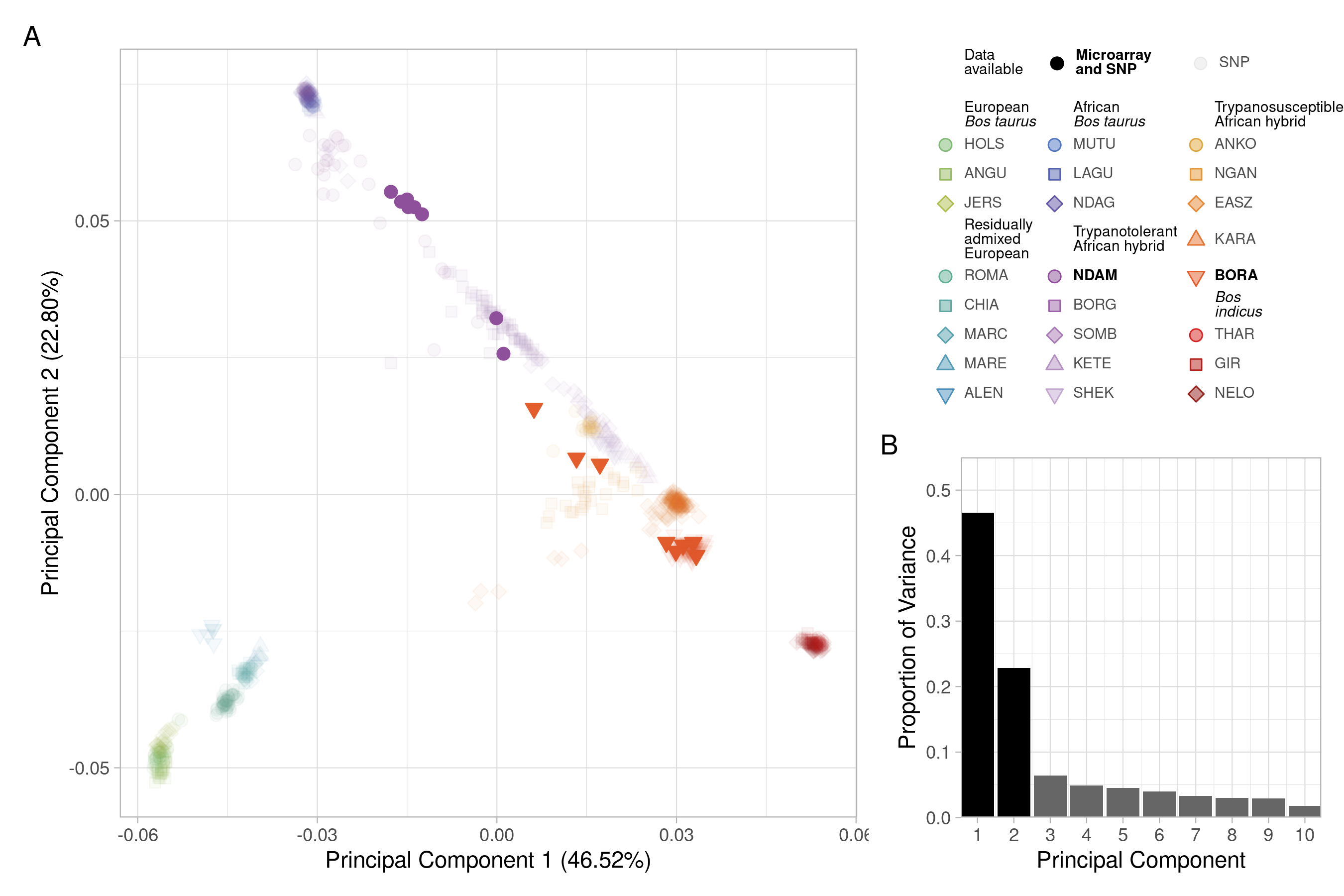

**Fig S5. A.** Principal component analysis (PCA) of the low-density SNP data for the cattle samples from McHugo and colleagues (McHugo *et al.* 2025) coloured according to population showing the first two principal components and **B.** bar chart of proportion of variance of the top ten principal components. The transparency indicates the availability of microarray gene expression data for the sample.

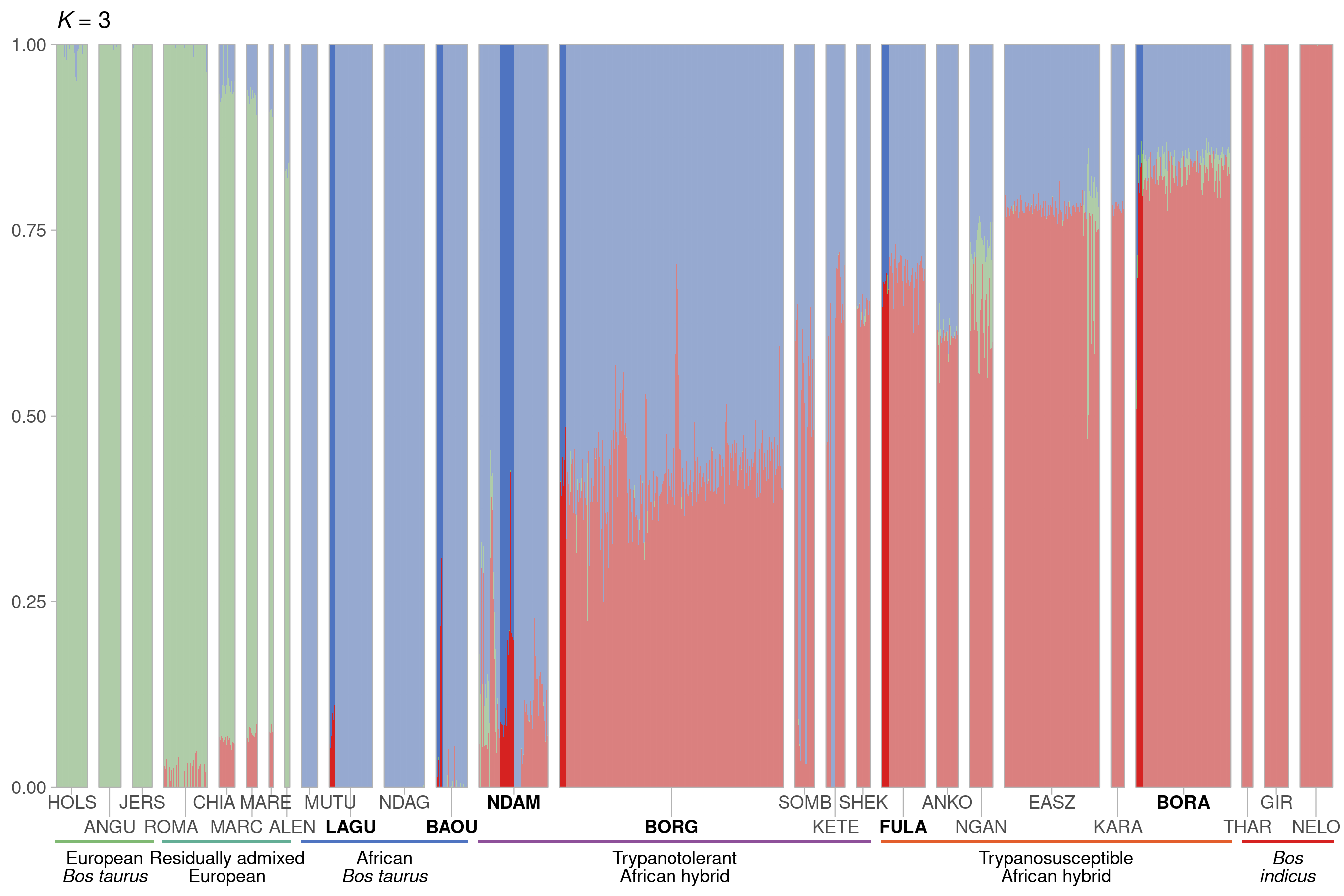

**Fig S6.** Hierarchical clustering of the SNP data for European, African, and Asian cattle populations. Results are shown for an assumed value of the number of ancestral populations *K* = 3. The transparency indicates the availability of gene expression data for the sample.

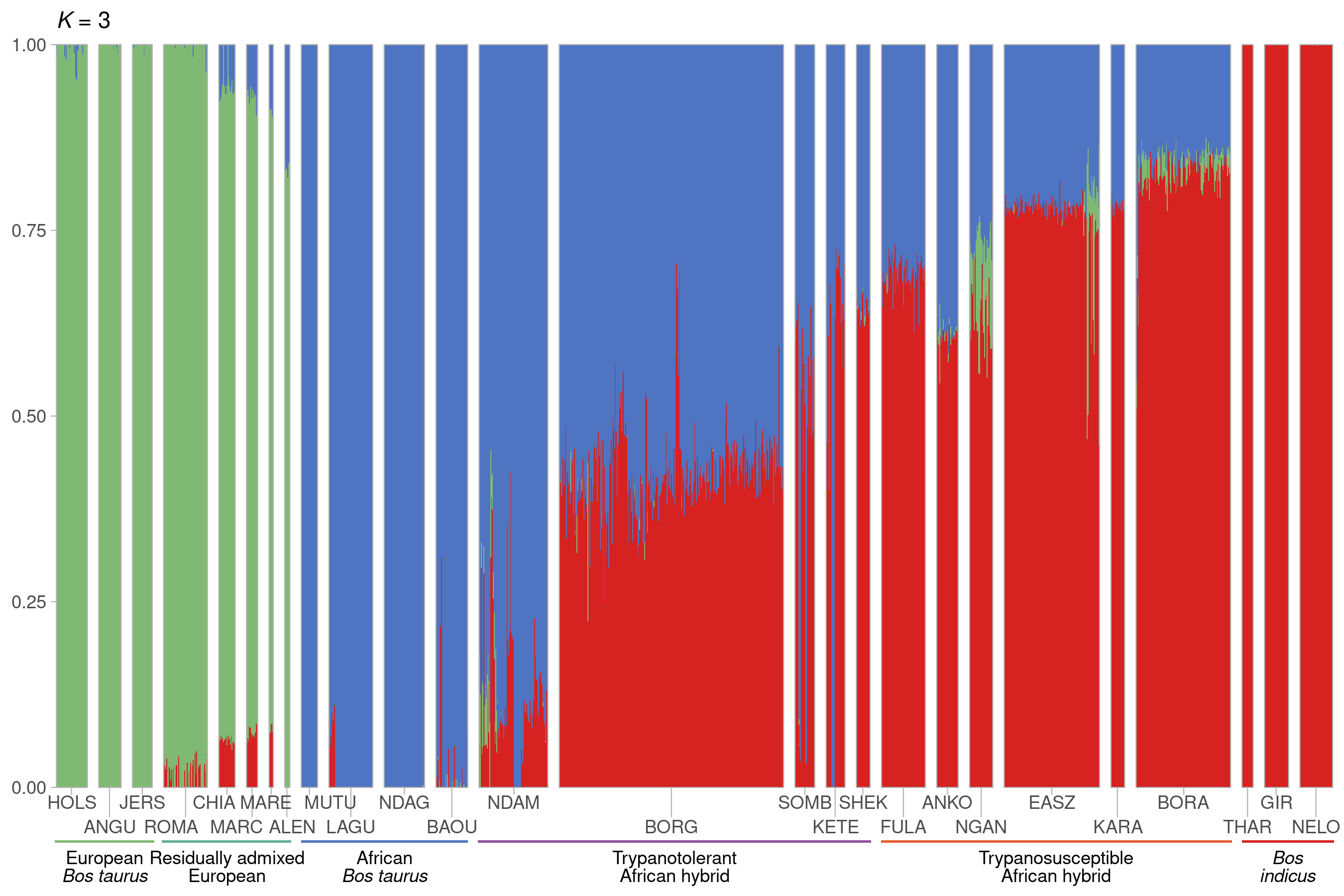

**Fig S7.** Hierarchical clustering of the SNP data for European, African, and Asian cattle populations. Results are shown for an assumed value of the number of ancestral populations *K* = 3.

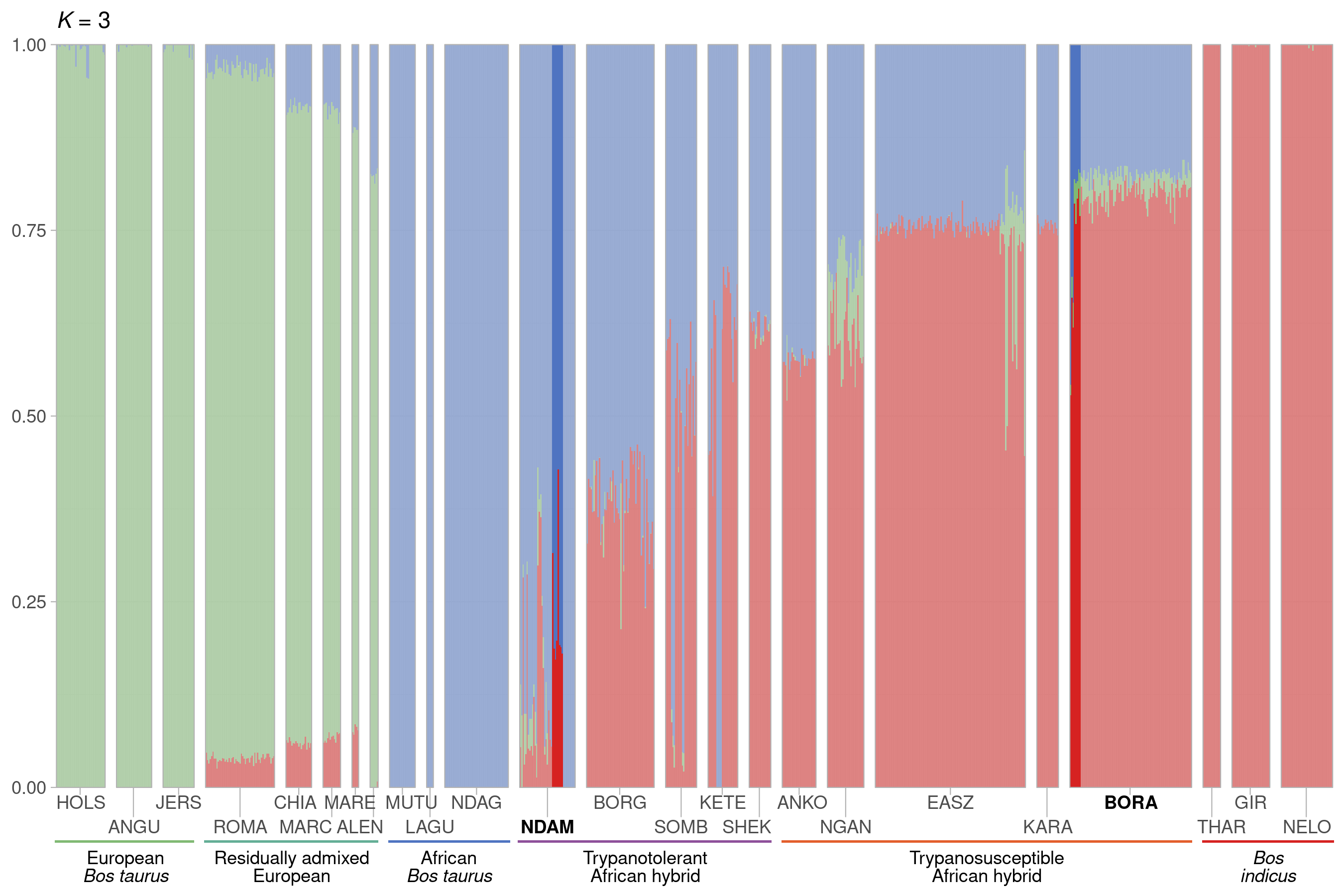

**Fig S8.** Hierarchical clustering of the high-density SNP data from McHugo and colleagues (McHugo *et al.* 2025) for European, African, and Asian cattle populations. Results are shown for an assumed value of the number of ancestral populations *K* = 3. The transparency indicates the availability of microarray gene expression data for the sample.

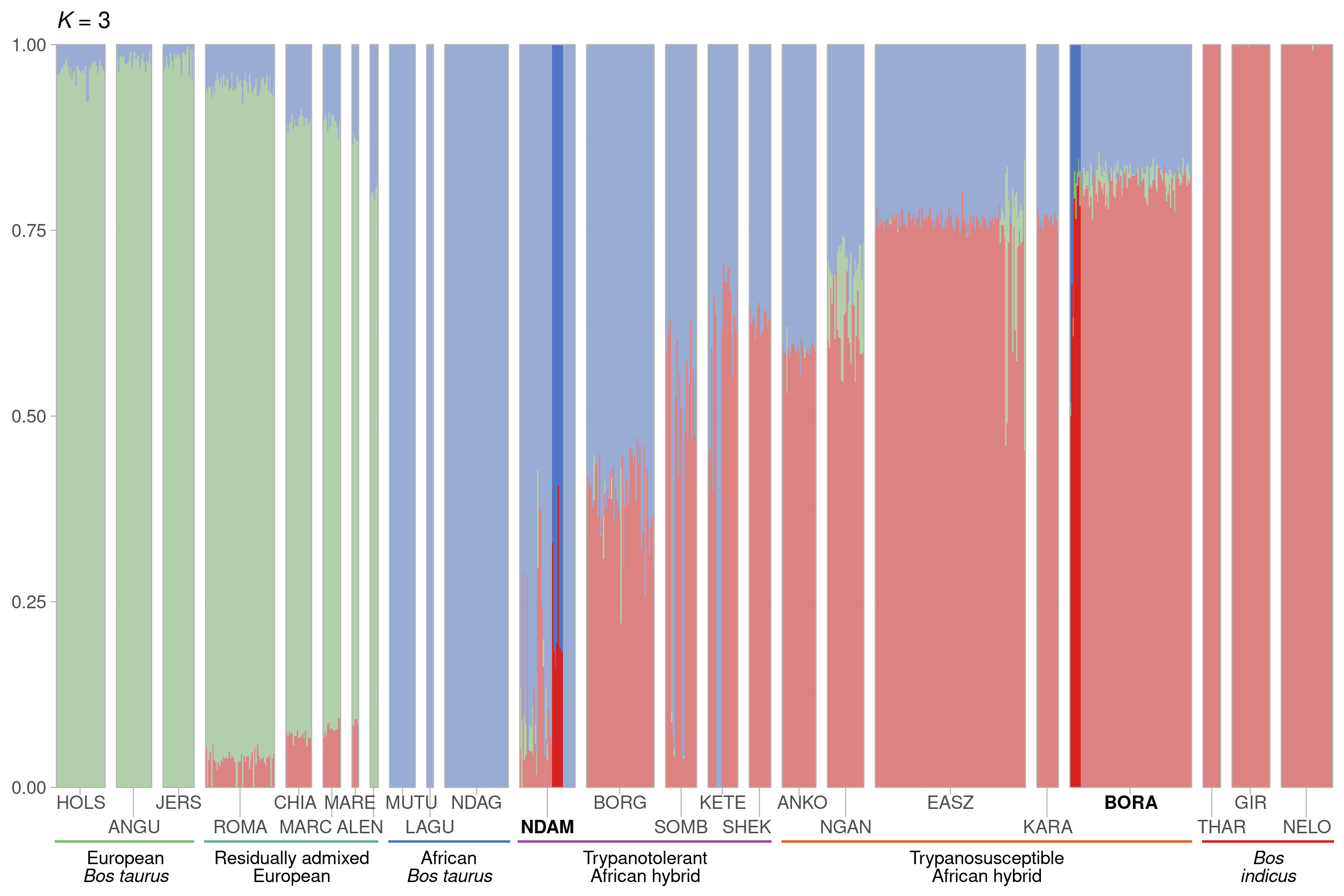

**Fig S9.** Hierarchical clustering of the low-density SNP data from McHugo and colleagues (McHugo *et al.* 2025) for European, African, and Asian cattle populations. Results are shown for an assumed value of the number of ancestral populations *K* = 3. The transparency indicates the availability of microarray gene expression data for the sample.

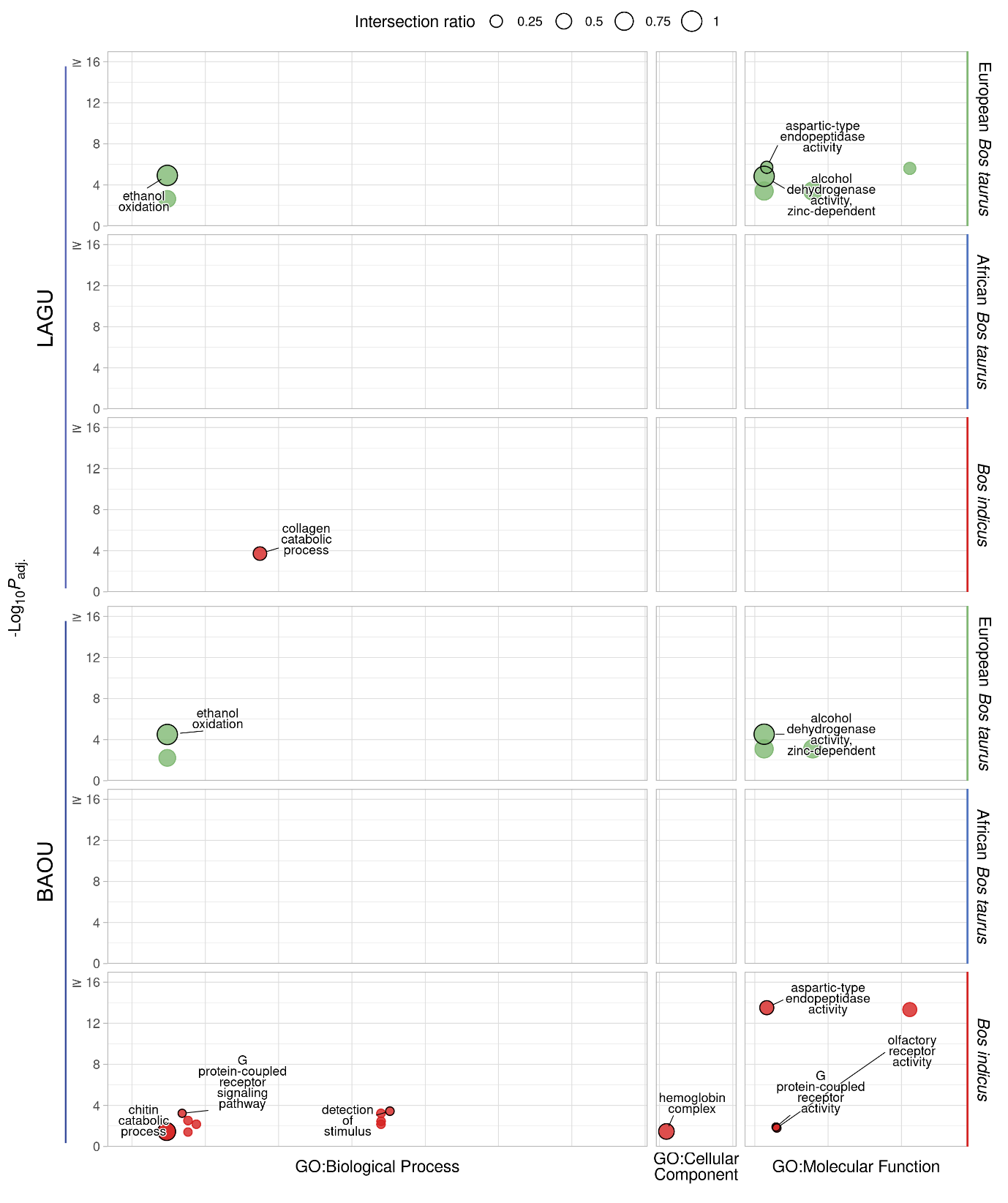

**Fig S10.** g:Profiler functional enrichment of introgressed regions in the African *B. taurus* cattle populations with gene expression data available according to local ancestry analysis of SNP data. Each dot represents a significantly enriched GO term with the size indicating the ratio of the intersection between the term and the introgressed genes. The *y*-axis shows the -log_10_*P*_adj._ value up to a maximum of 16 and the panels along the *y*-axis and colours indicate the ancestry component. The panels along the *x*-axis indicate the source of the term and the position within the panels groups terms from the same GO subtree. The top driver GO terms up to a maximum of ten are indicated with a black outline and label.

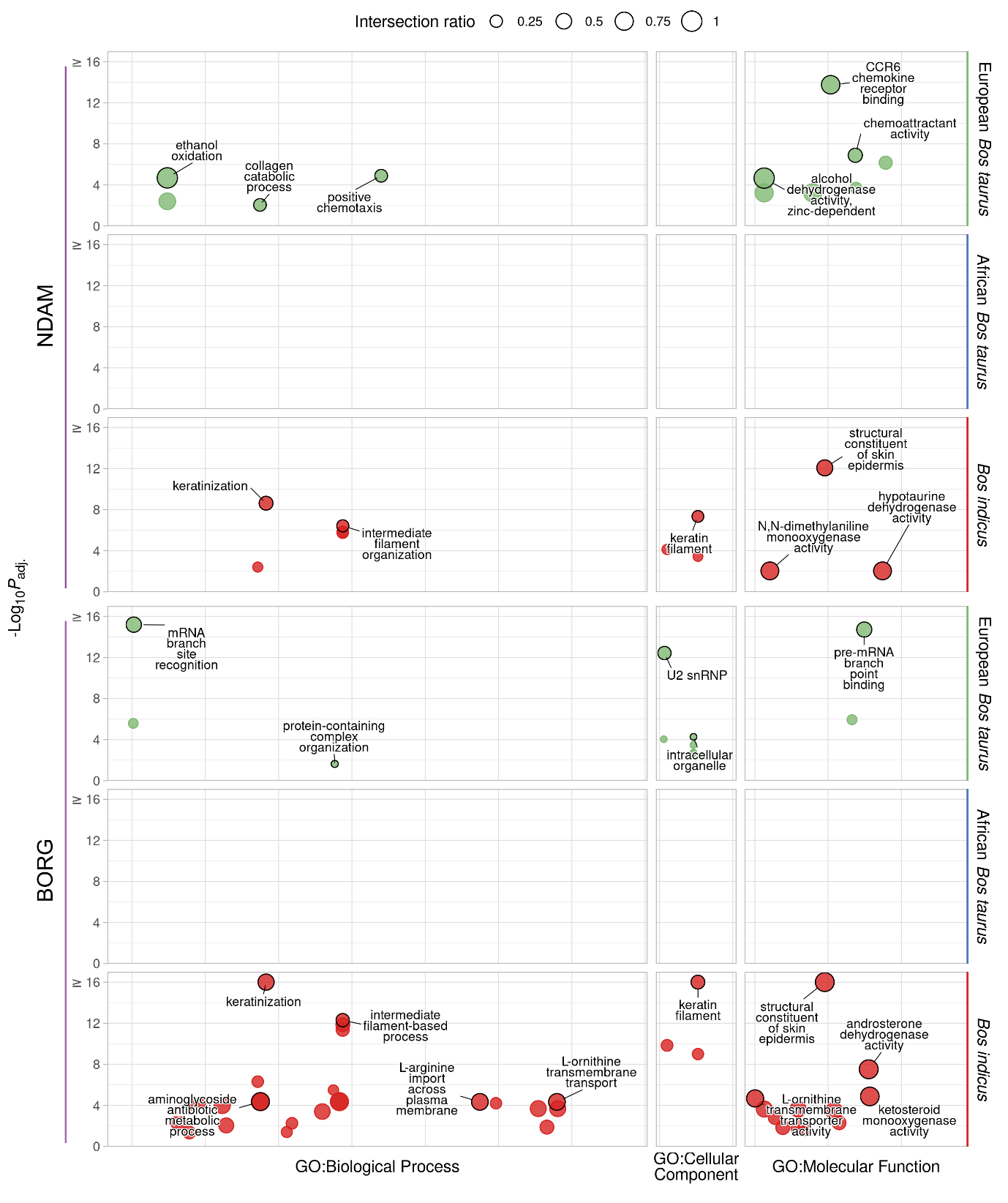

**Fig S11.** g:Profiler functional enrichment of introgressed regions in the trypanotolerant African hybrid cattle populations with gene expression data available according to local ancestry analysis of SNP data. Each dot represents a significantly enriched GO term with the size indicating the ratio of the intersection between the term and the introgressed genes. The *y*-axis shows the -log_10_*P*_adj._ value up to a maximum of 16 and the panels along the *y*-axis and colours indicate the ancestry component. The panels along the *x*-axis indicate the source of the term and the position within the panels groups terms from the same GO subtree. The top driver GO terms up to a maximum of ten are indicated with a black outline and label.

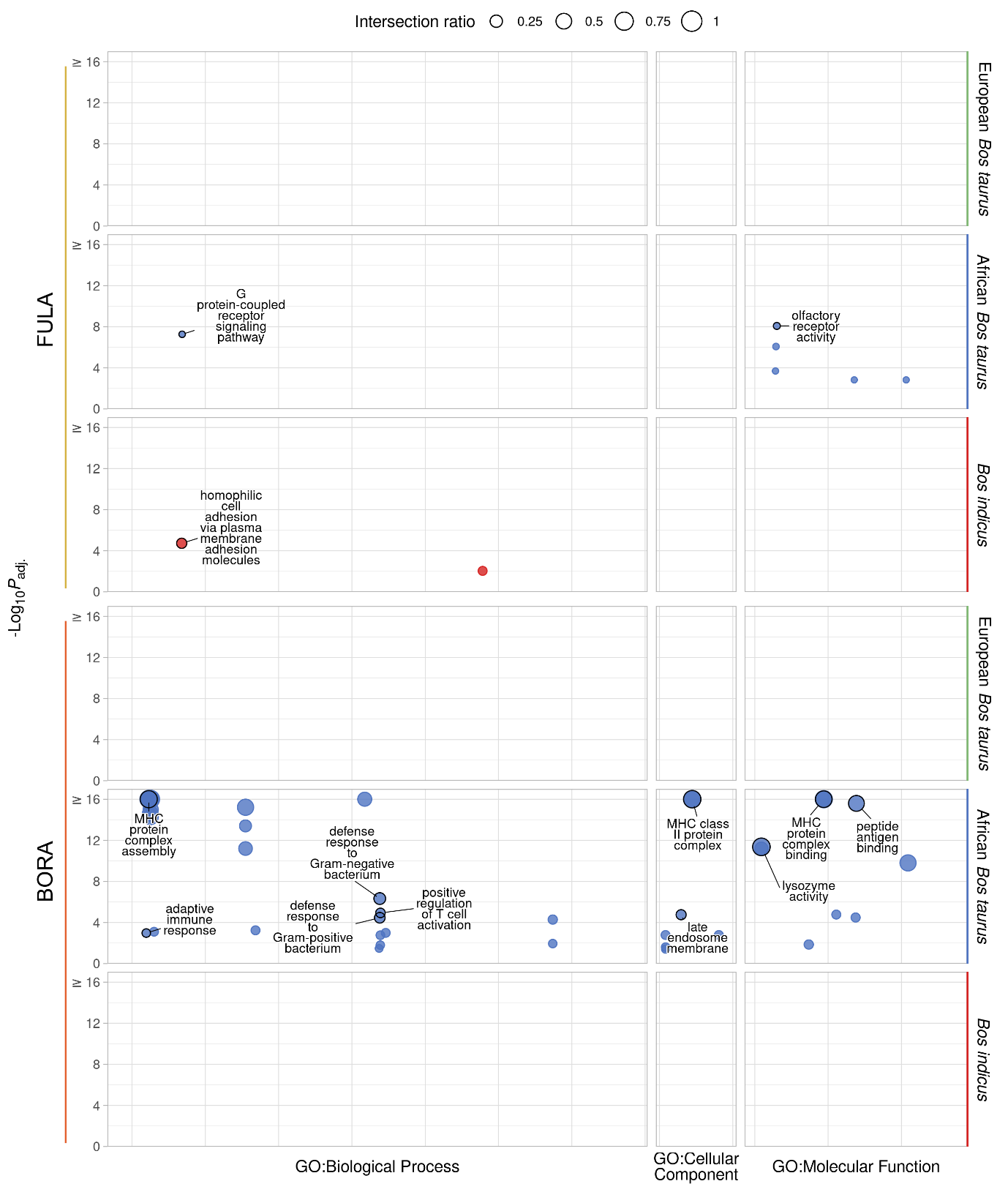

**Fig S12.** g:Profiler functional enrichment of introgressed regions in the trypanosusceptible African hybrid cattle populations with gene expression data available according to local ancestry analysis of SNP data. Each dot represents a significantly enriched GO term with the size indicating the ratio of the intersection between the term and the introgressed genes. The *y*-axis shows the -log_10_*P*_adj._ value up to a maximum of 16 and the panels along the *y*-axis and colours indicate the ancestry component. The panels along the *x*-axis indicate the source of the term and the position within the panels groups terms from the same GO subtree. The top driver GO terms up to a maximum of ten are indicated with a black outline and label.

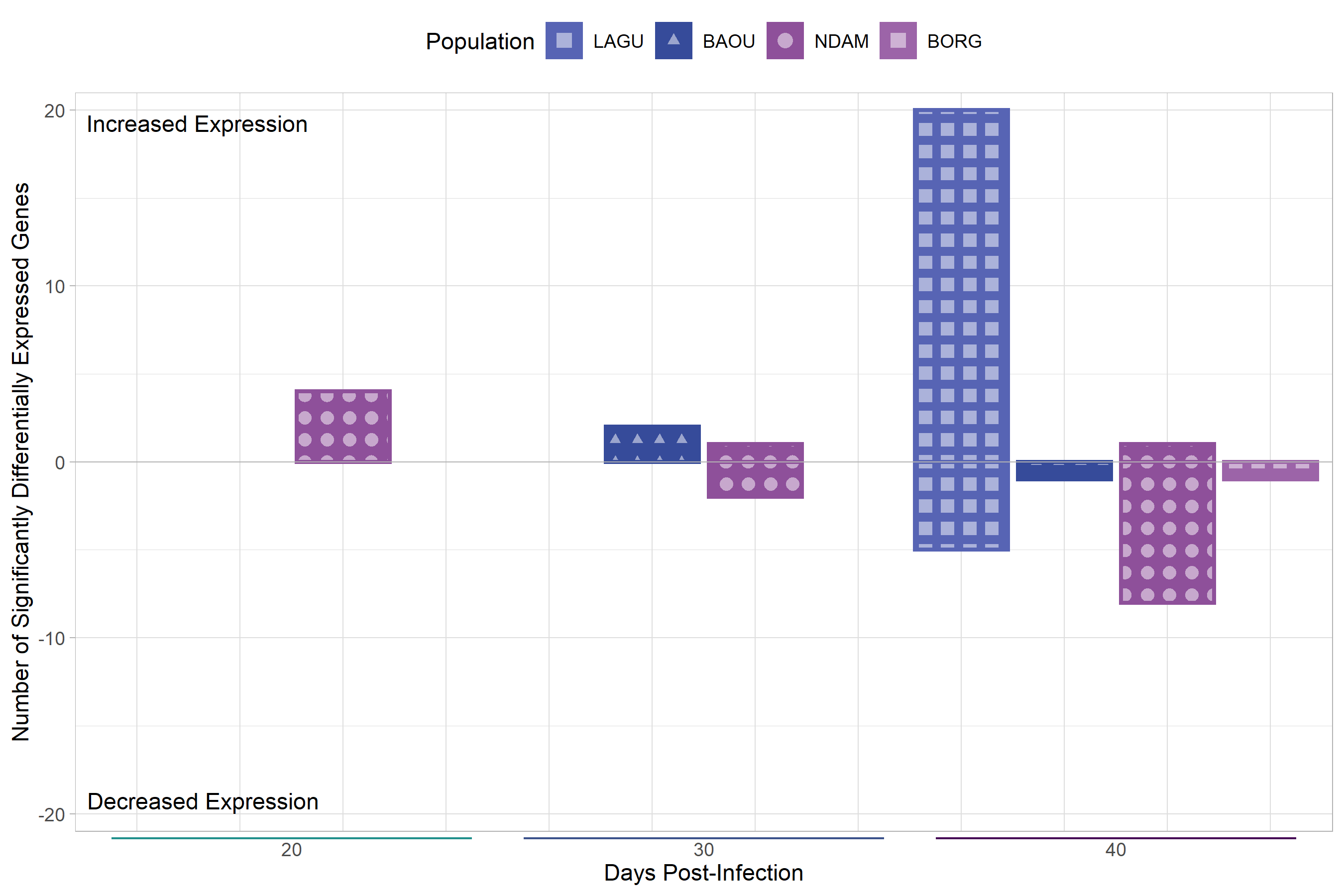

**Fig S13.** Bar chart showing the numbers of significantly differentially expressed genes for the response contrasts of the RNA-seq data. The extent of the bar above and below 0 on the *y*-axis indicates the numbers of significantly differentially expressed genes with increased and decreased expression respectively. The position on the *x*-axis indicates the number of days post-infection and the colour and shapes within the bars represent the population.

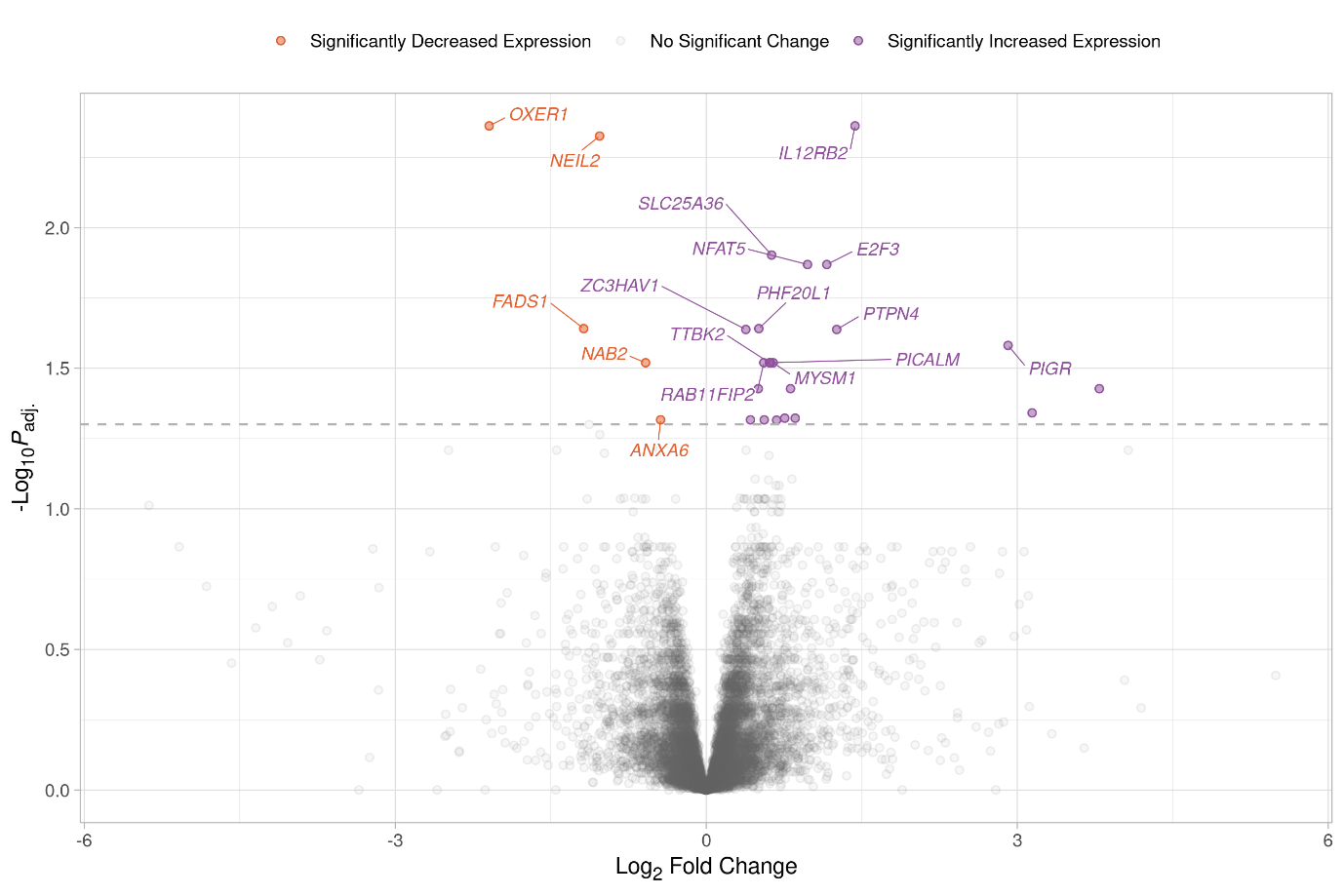

**Fig S14.** Volcano plot showing the results of the response contrast for the RNA-seq data from the LAGU population at 40 days post-infection. Each dot represents a gene with the position on the *x*- and *y*-axes indicating the log_2_ fold change and -log_10_*P*_adj._, respectively. Genes above the horizontal dashed line are significantly differentially expressed with the colours representing the change in expression. The top 10 most significant genes for increased and decreased expression with gene symbols are labelled.

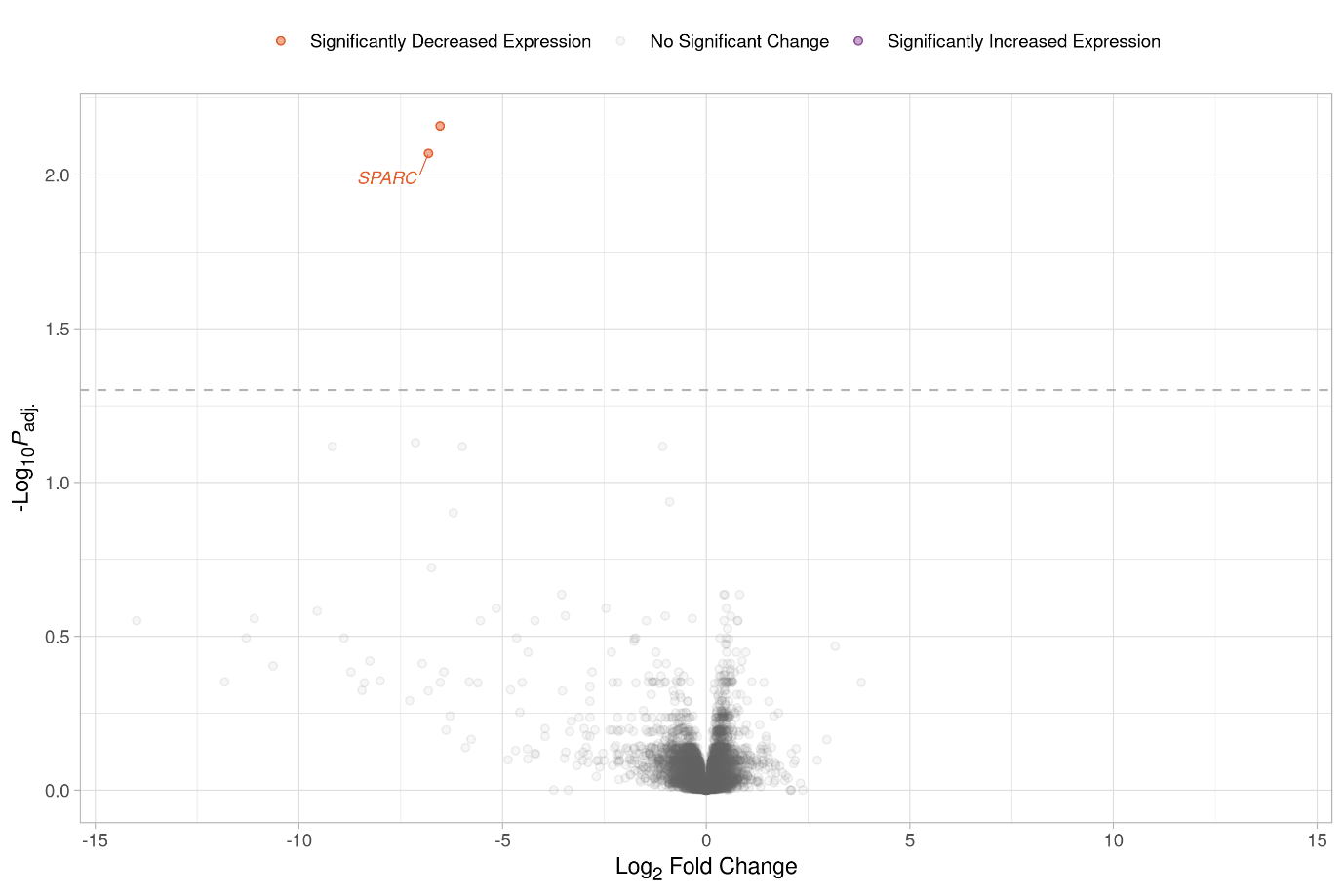

**Fig S15.** Volcano plot showing the results of the response contrast for the RNA-seq data from the BAOU population at 40 days post-infection. Each dot represents a gene with the position on the *x*- and *y*-axes indicating the log_2_ fold change and -log_10_*P*_adj._, respectively. Genes above the horizontal dashed line are significantly differentially expressed with the colours representing the change in expression. The top 10 most significant genes for increased and decreased expression with gene symbols are labelled.

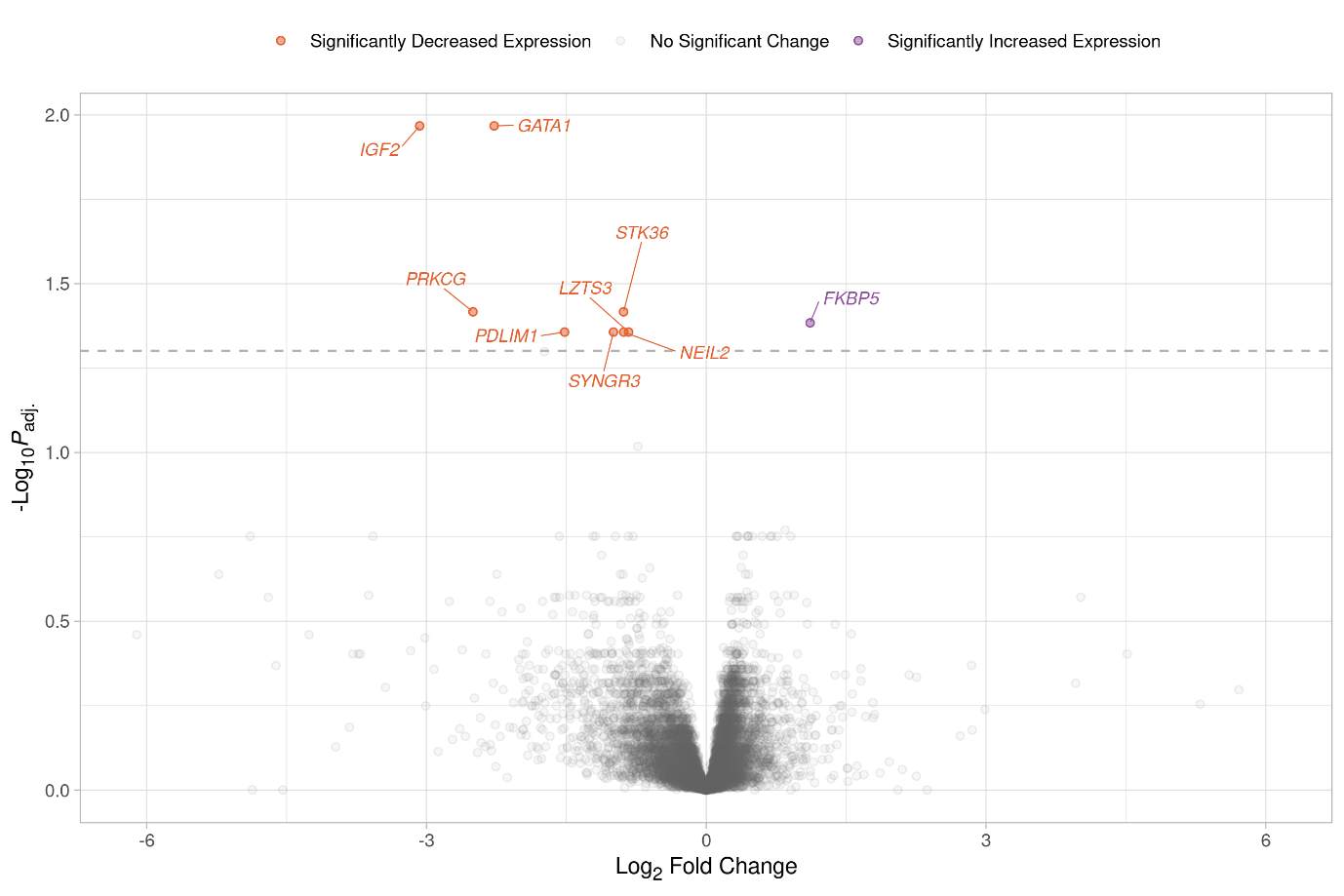

**Fig S16.** Volcano plot showing the results of the response contrast for the RNA-seq data from the NDAM population at 40 days post-infection. Each dot represents a gene with the position on the *x*- and *y*-axes indicating the log_2_ fold change and -log_10_*P*_adj._, respectively. Genes above the horizontal dashed line are significantly differentially expressed with the colours representing the change in expression. The top 10 most significant genes for increased and decreased expression with gene symbols are labelled.

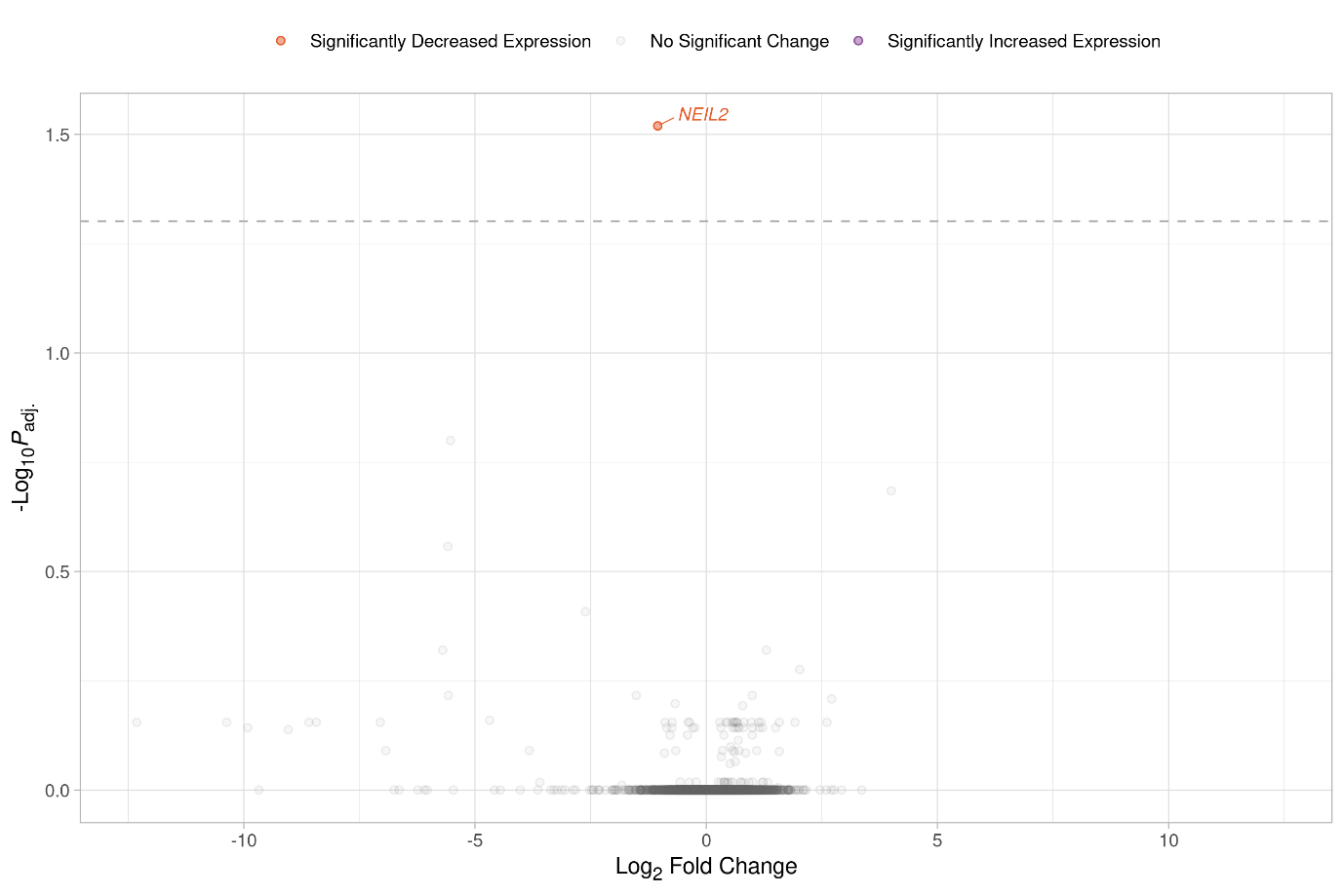

**Fig S17.** Volcano plot showing the results of the response contrast for the RNA-seq data from the BORG population at 40 days post-infection. Each dot represents a gene with the position on the *x*- and *y*-axes indicating the log_2_ fold change and -log_10_*P*_adj._, respectively. Genes above the horizontal dashed line are significantly differentially expressed with the colours representing the change in expression. The top 10 most significant genes for increased and decreased expression with gene symbols are labelled.

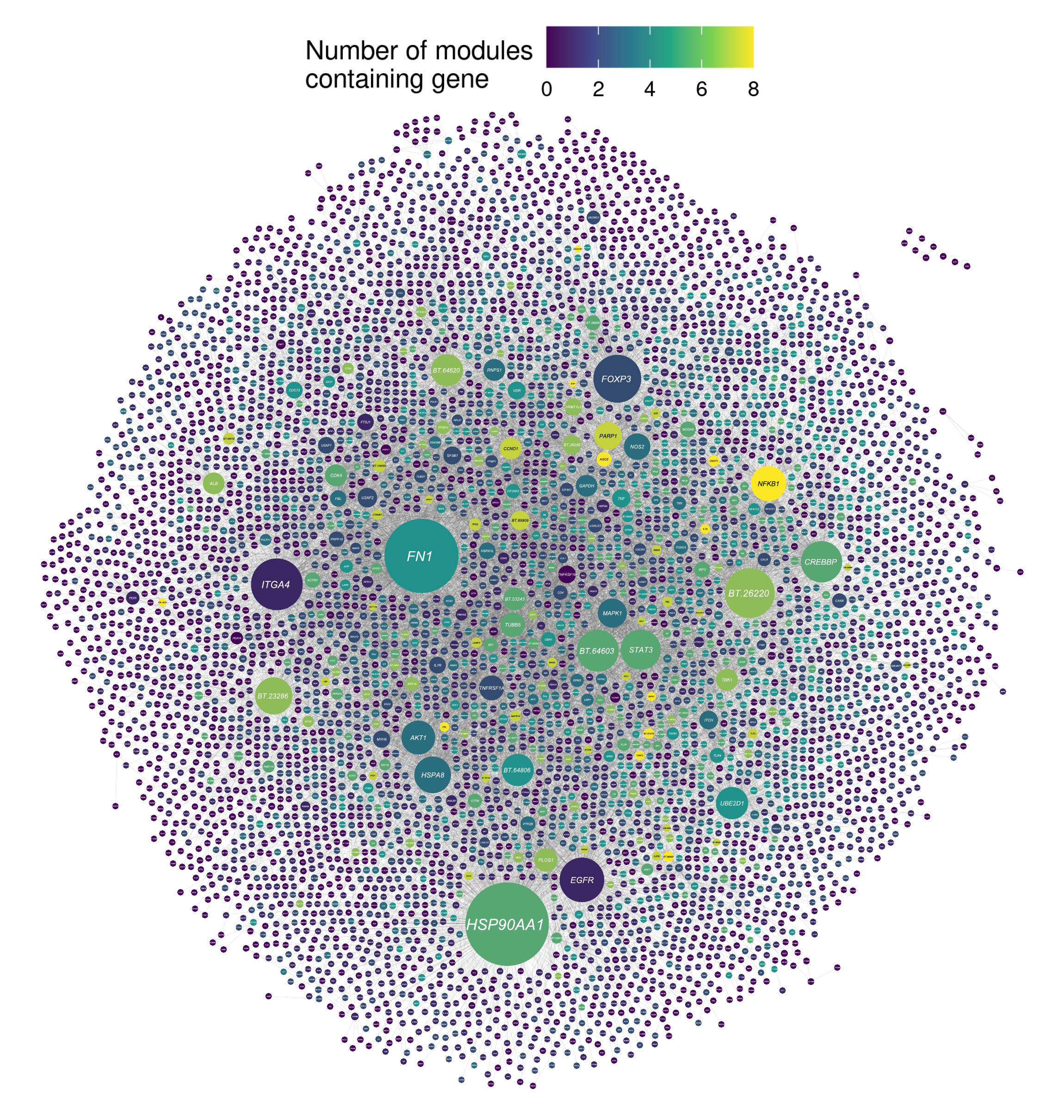

**Fig S18.** Base network generated using InnateDB with the top results of a search of the GeneCards^®^ for genes relating to the term “trypano*”. Each node in the network represents a gene coloured according to the number of functional modules that contain that gene, while the edges connecting the nodes represent gene interactions. The nodes are sized according to their degree or number of interactions.

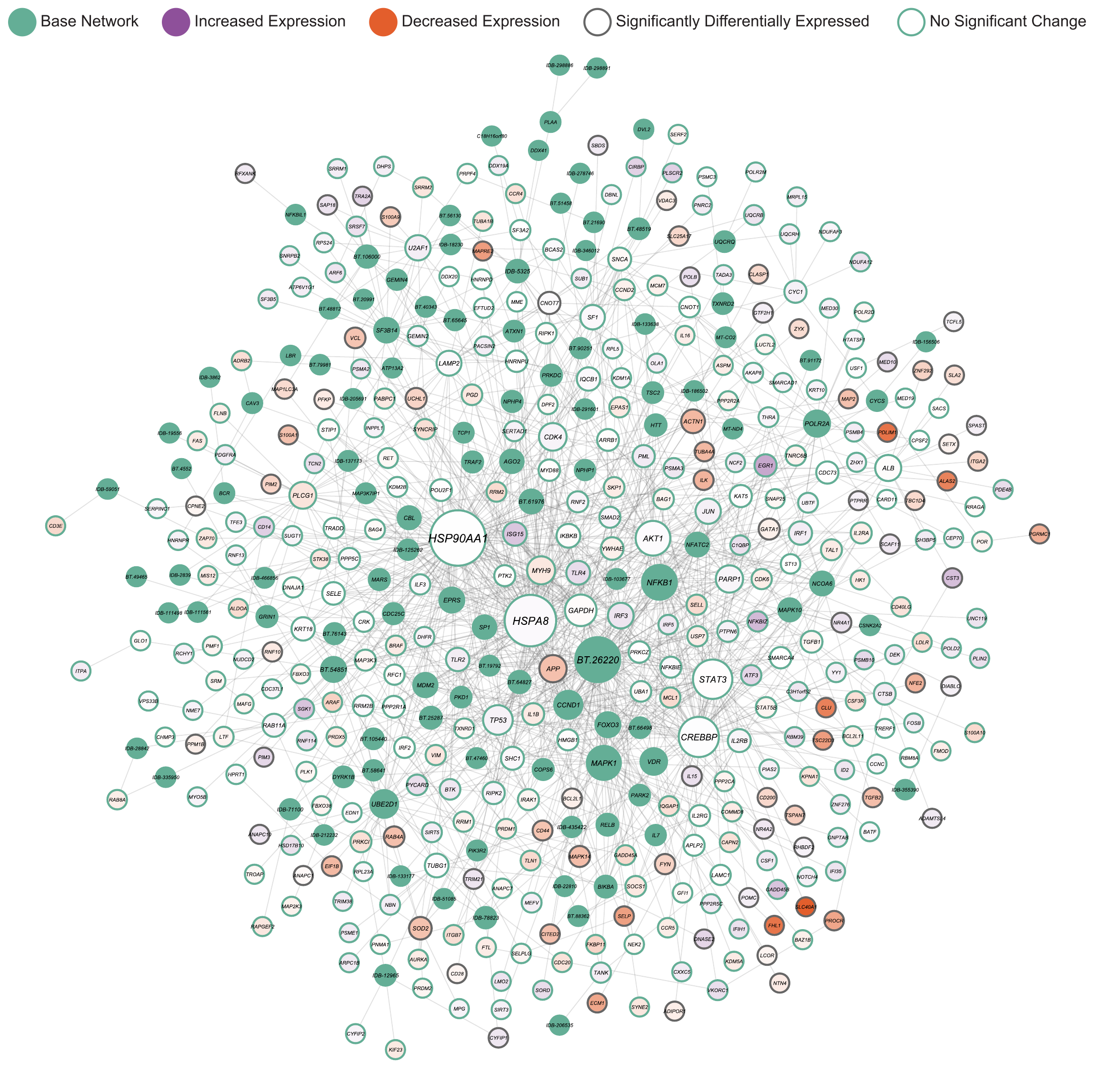

**Fig S19.** Functional module identified using jActiveModules and differential expression results for the MICRO BL 34 contrast. Each node in the network represents a gene coloured according to expression with significant differential expression indicated by the outline. The edges connecting the nodes represent gene interactions and the nodes are sized according to their number of interactions or degree.

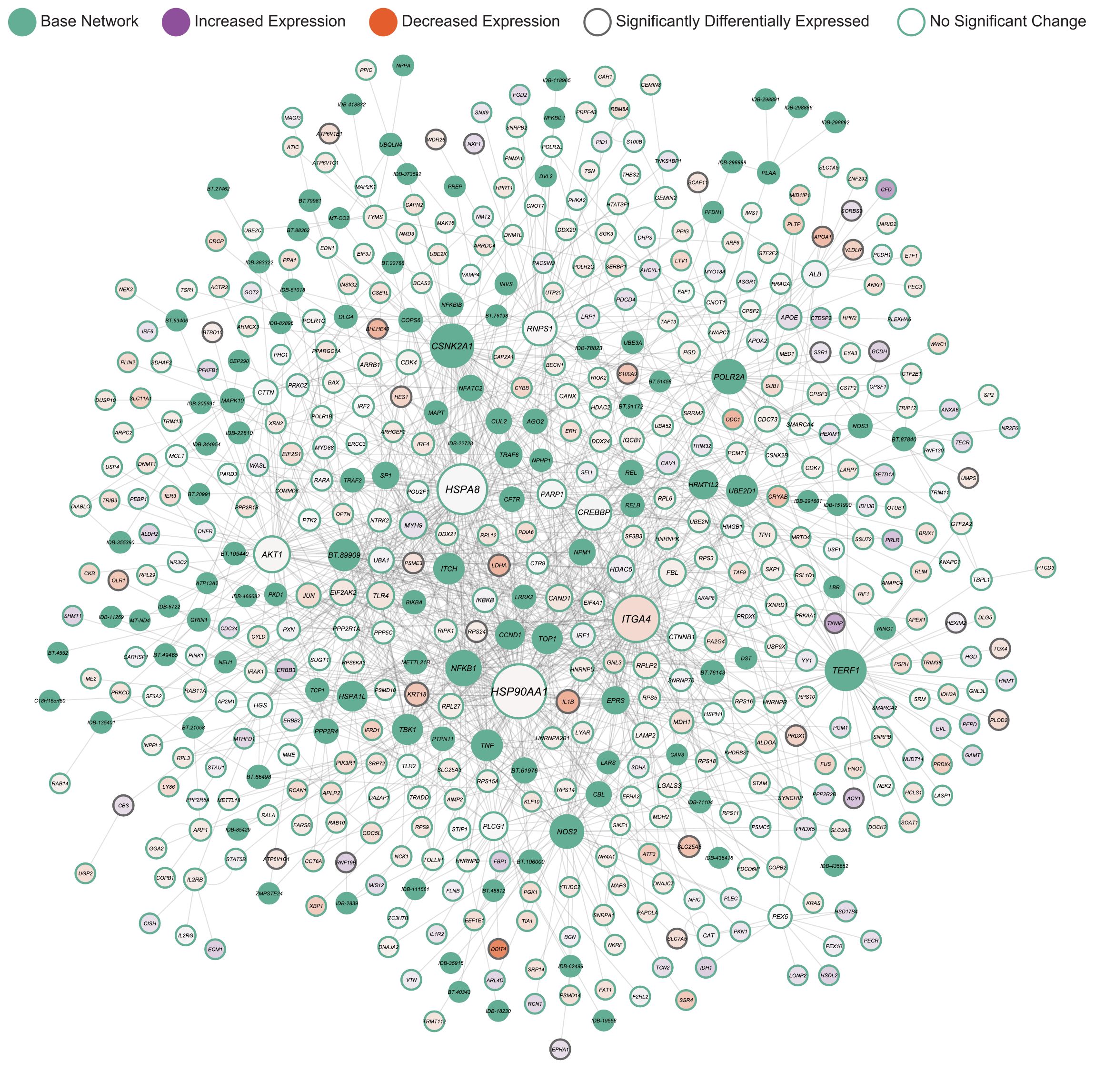

**Fig S20.** Functional module identified using jActiveModules and differential expression results for the MICRO LI 35 contrast. Each node in the network represents a gene coloured according to expression with significant differential expression indicated by the outline. The edges connecting the nodes represent gene interactions and the nodes are sized according to their number of interactions or degree.

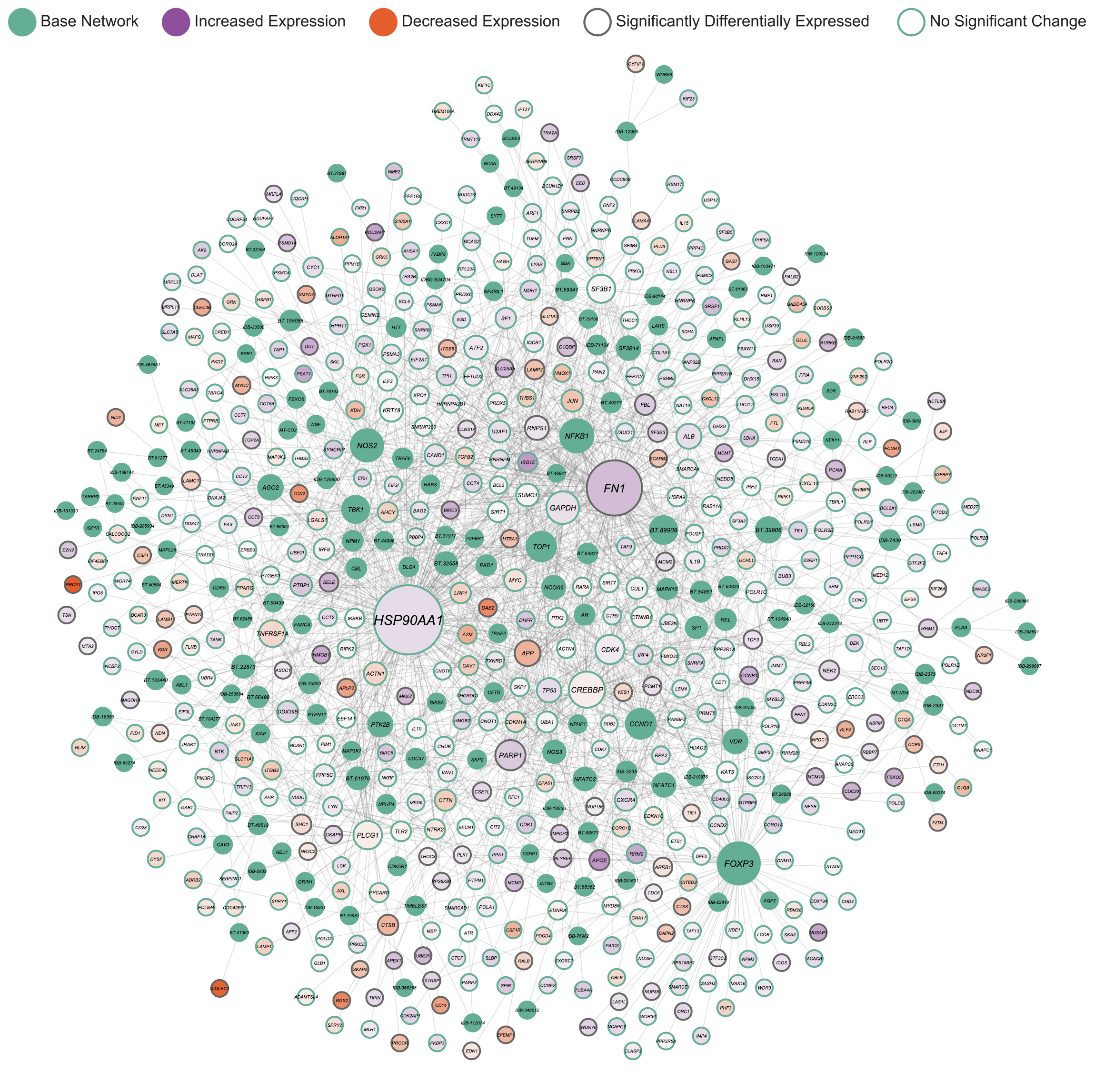

**Fig S21.** Functional module identified using jActiveModules and differential expression results for the MICRO LN 35 contrast. Each node in the network represents a gene coloured according to expression with significant differential expression indicated by the outline. The edges connecting the nodes represent gene interactions and the nodes are sized according to their number of interactions or degree.

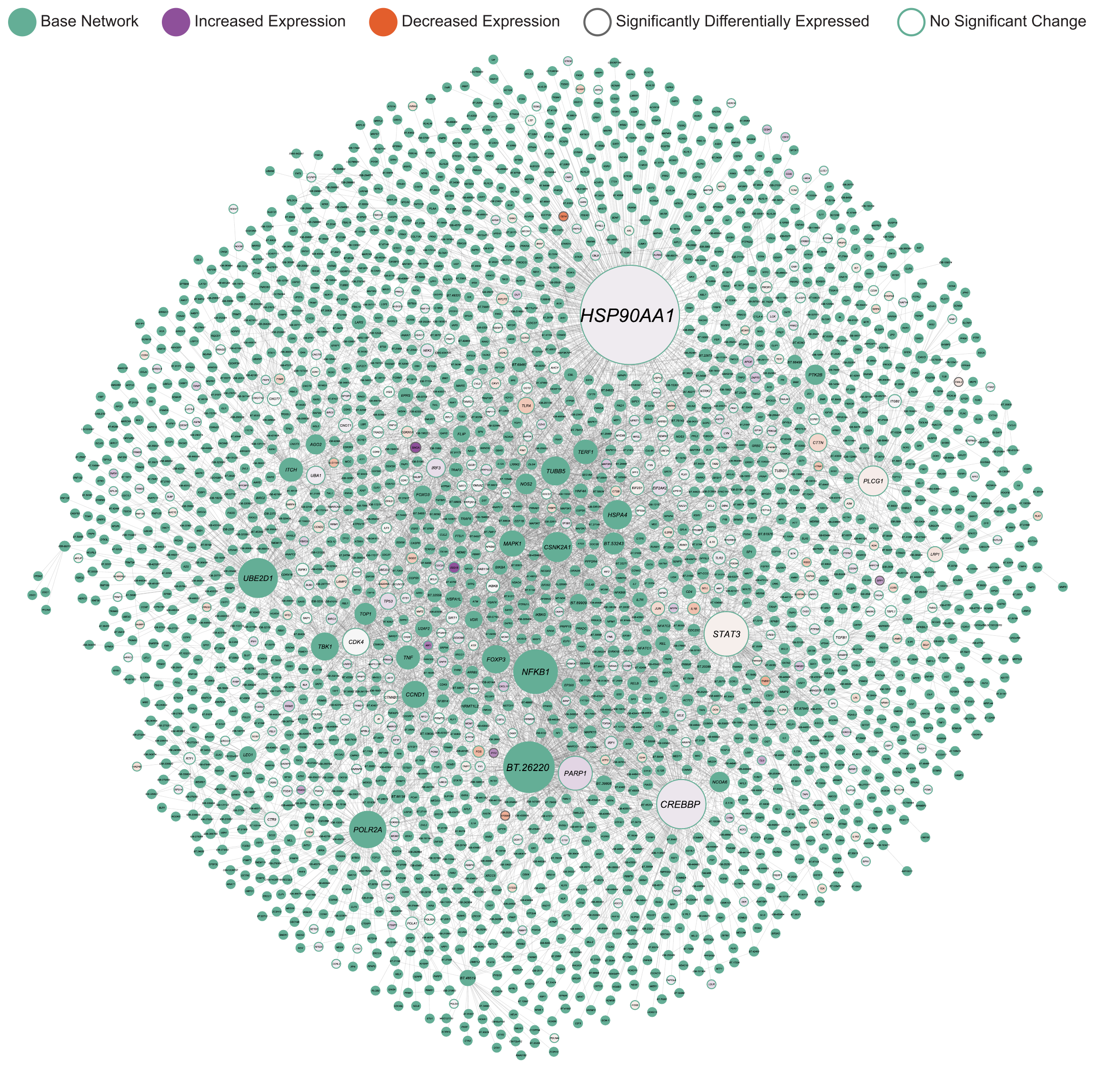

**Fig S22.** Functional module identified using jActiveModules and differential expression results for the MICRO SP 35 contrast. Each node in the network represents a gene coloured according to expression with significant differential expression indicated by the outline. The edges connecting the nodes represent gene interactions and the nodes are sized according to their number of interactions or degree.

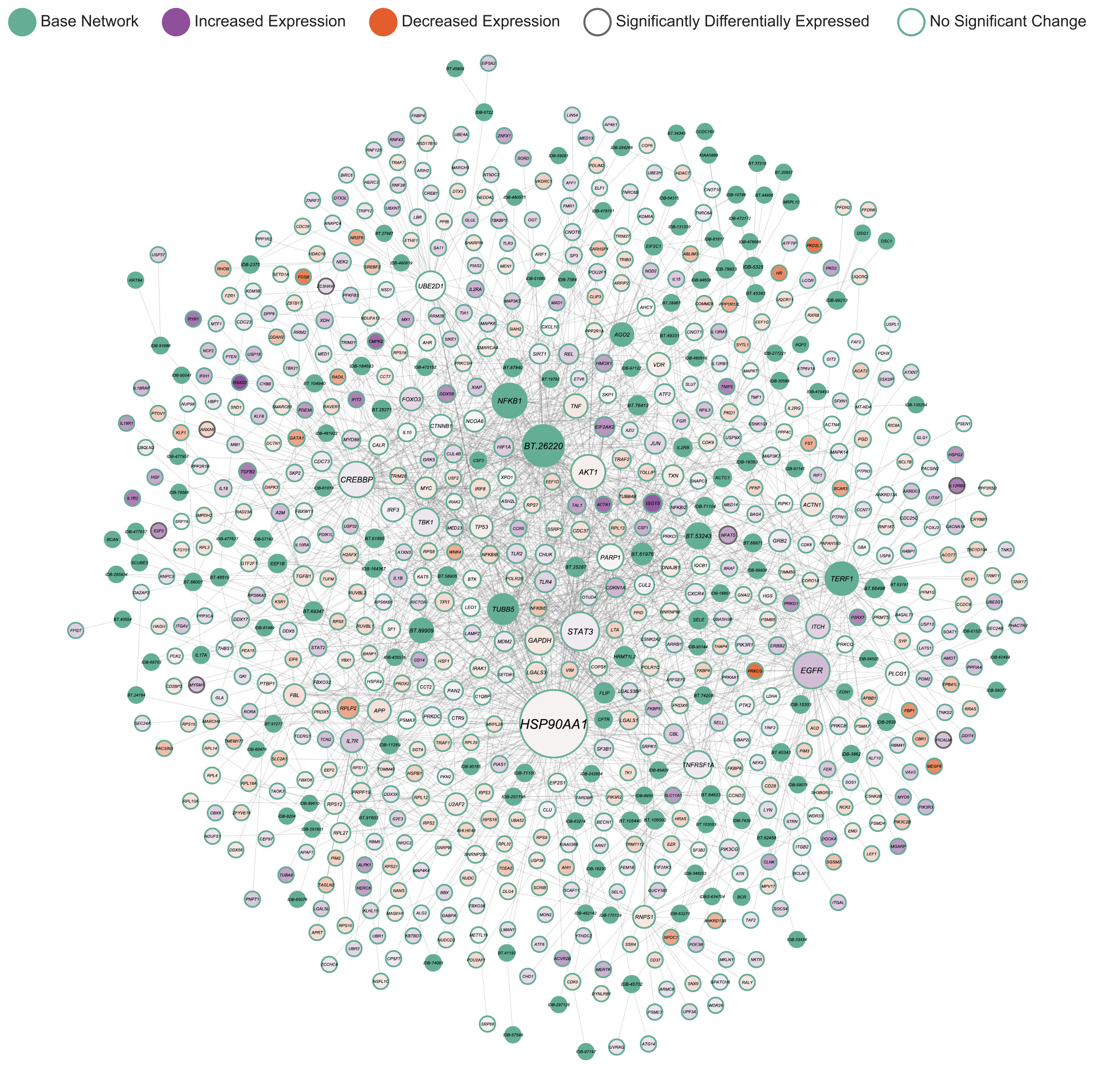

**Fig S23.** Functional module identified using jActiveModules and differential expression results for the RNA LAGU 40 contrast. Each node in the network represents a gene coloured according to expression with significant differential expression indicated by the outline. The edges connecting the nodes represent gene interactions and the nodes are sized according to their number of interactions or degree.

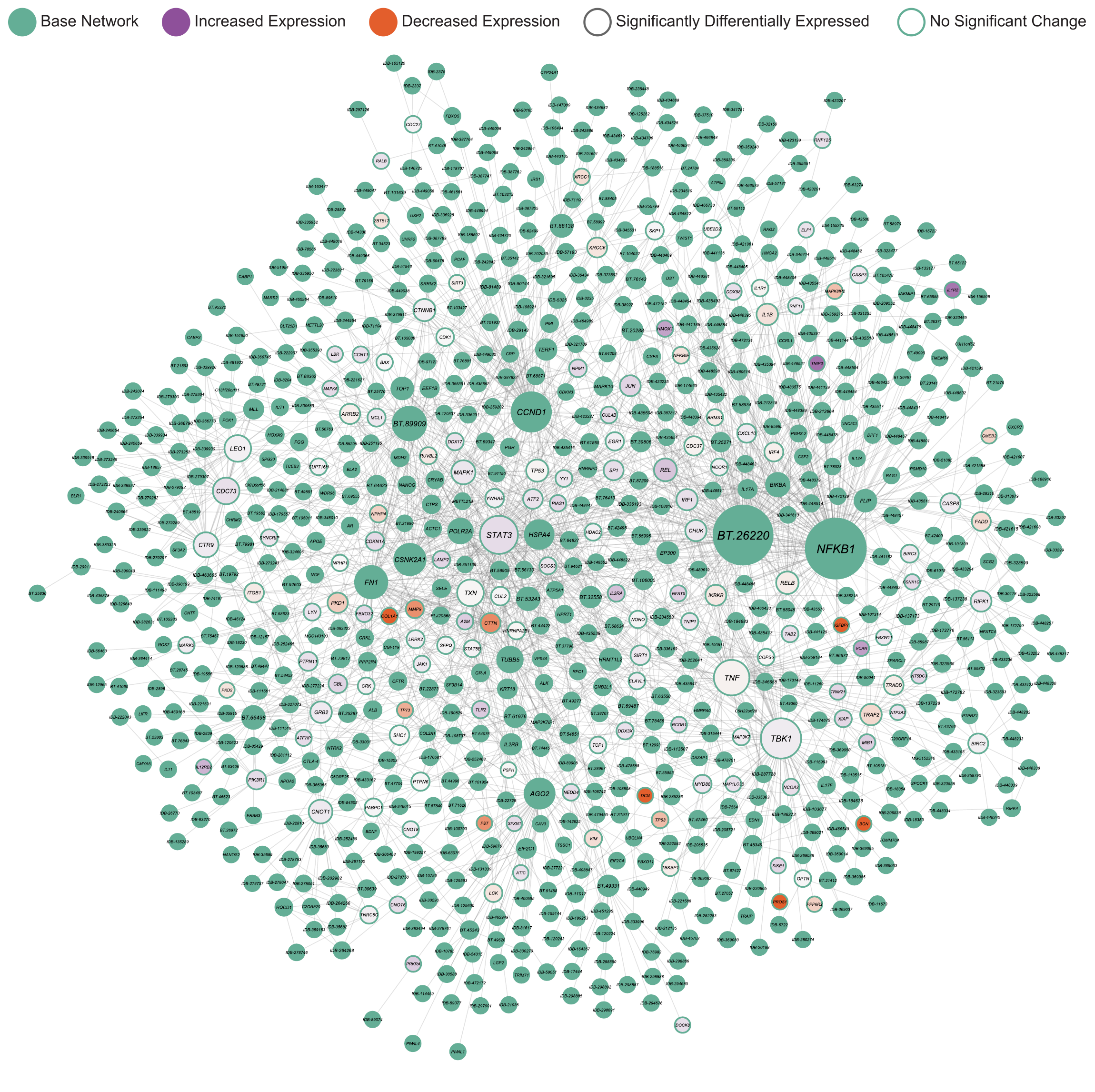

**Fig S24.** Functional module identified using jActiveModules and differential expression results for the RNA BAOU 40 contrast. Each node in the network represents a gene coloured according to expression with significant differential expression indicated by the outline. The edges connecting the nodes represent gene interactions and the nodes are sized according to their number of interactions or degree.

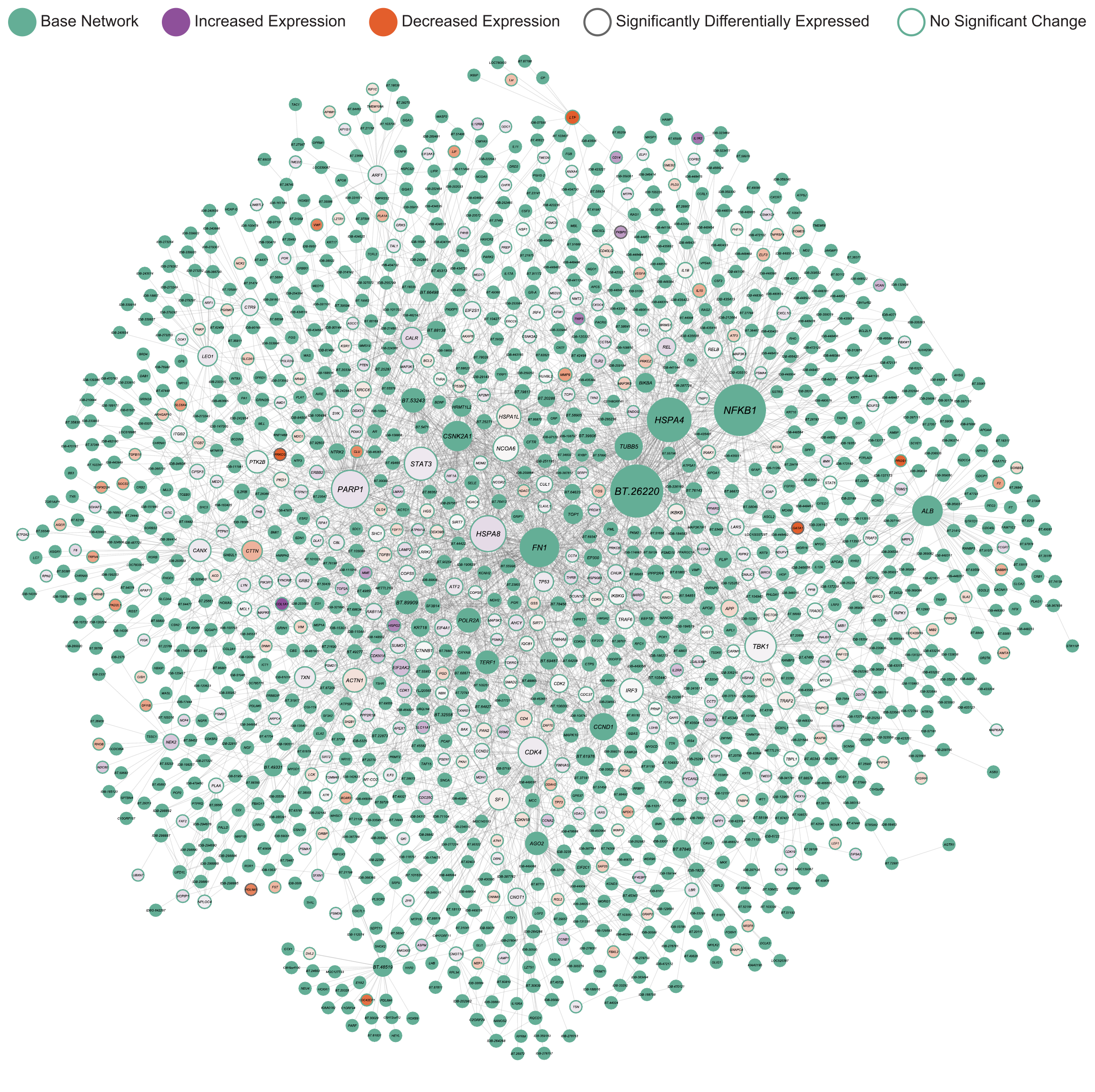

**Fig S25.** Functional module identified using jActiveModules and differential expression results for the RNA NDAM 40 contrast. Each node in the network represents a gene coloured according to expression with significant differential expression indicated by the outline. The edges connecting the nodes represent gene interactions and the nodes are sized according to their number of interactions or degree.

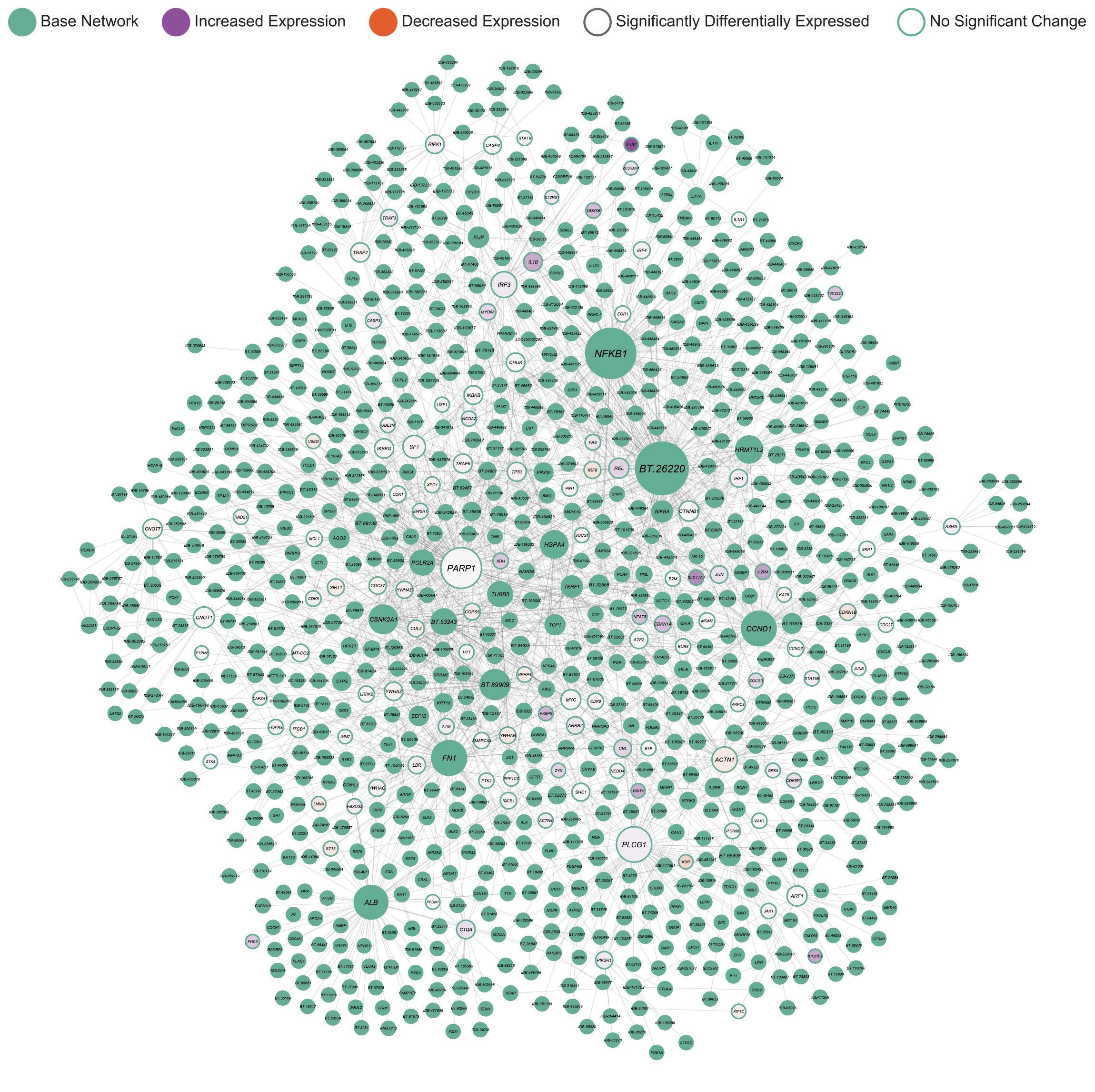

**Fig S26.** Functional module identified using jActiveModules and differential expression results for the RNA BORG 40 contrast. Each node in the network represents a gene coloured according to expression with significant differential expression indicated by the outline. The edges connecting the nodes represent gene interactions and the nodes are sized according to their number of interactions or degree.

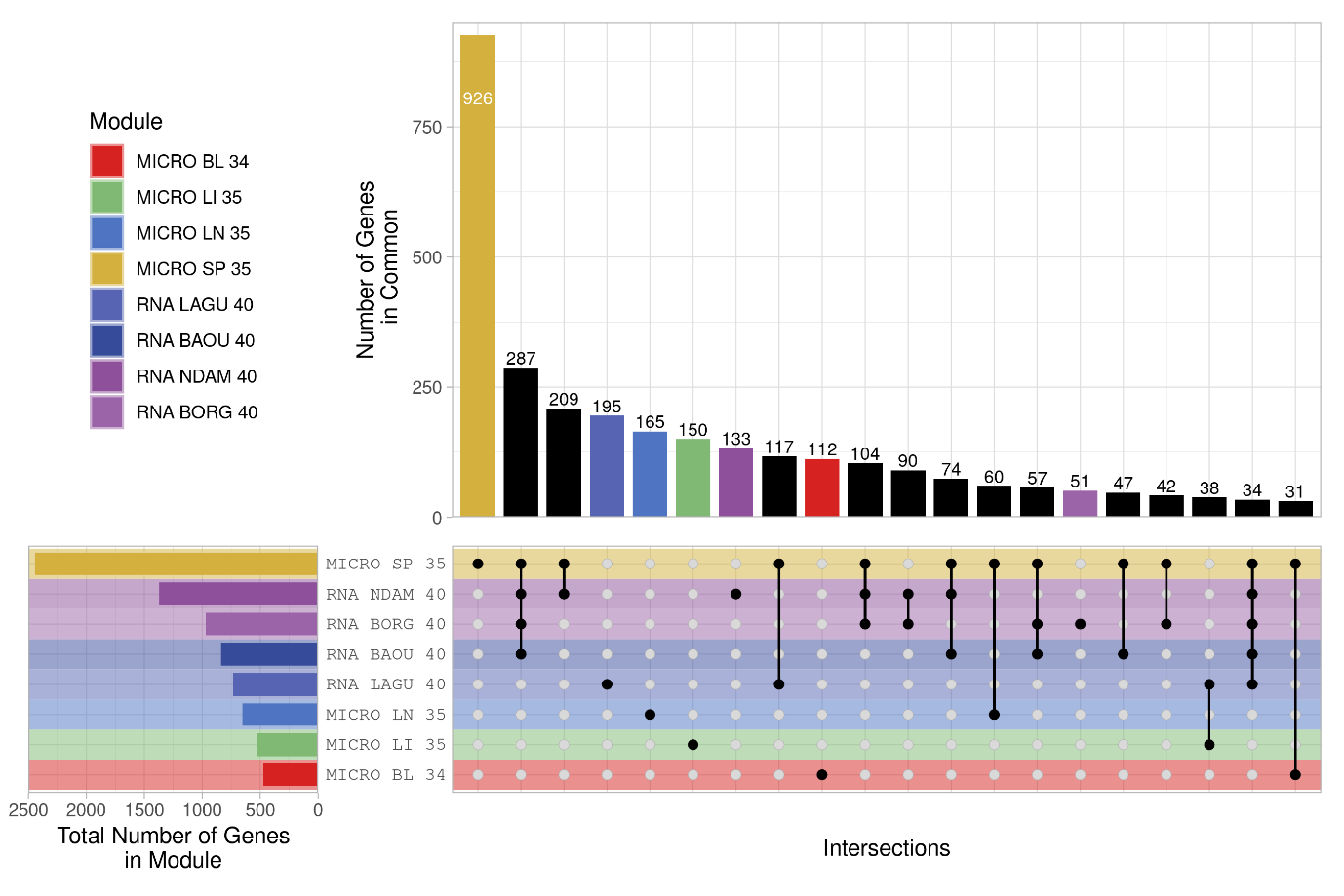

**Fig S27.** Upset plot showing the top 20 intersections between the genes in the functional modules identified using jActiveModules and differential expression results for each of the final response contrasts for the microarray and RNA-seq data. The horizontal bars indicate the total number of genes in each module while the vertical bars indicate the number of genes in common between the modules annotated with black dots connected by lines in the intersection matrix. The background colour of the stripes in the intersection matrix and colour of the horizontal and vertical bars represent the module. Black bars indicate an overlap between different modules.

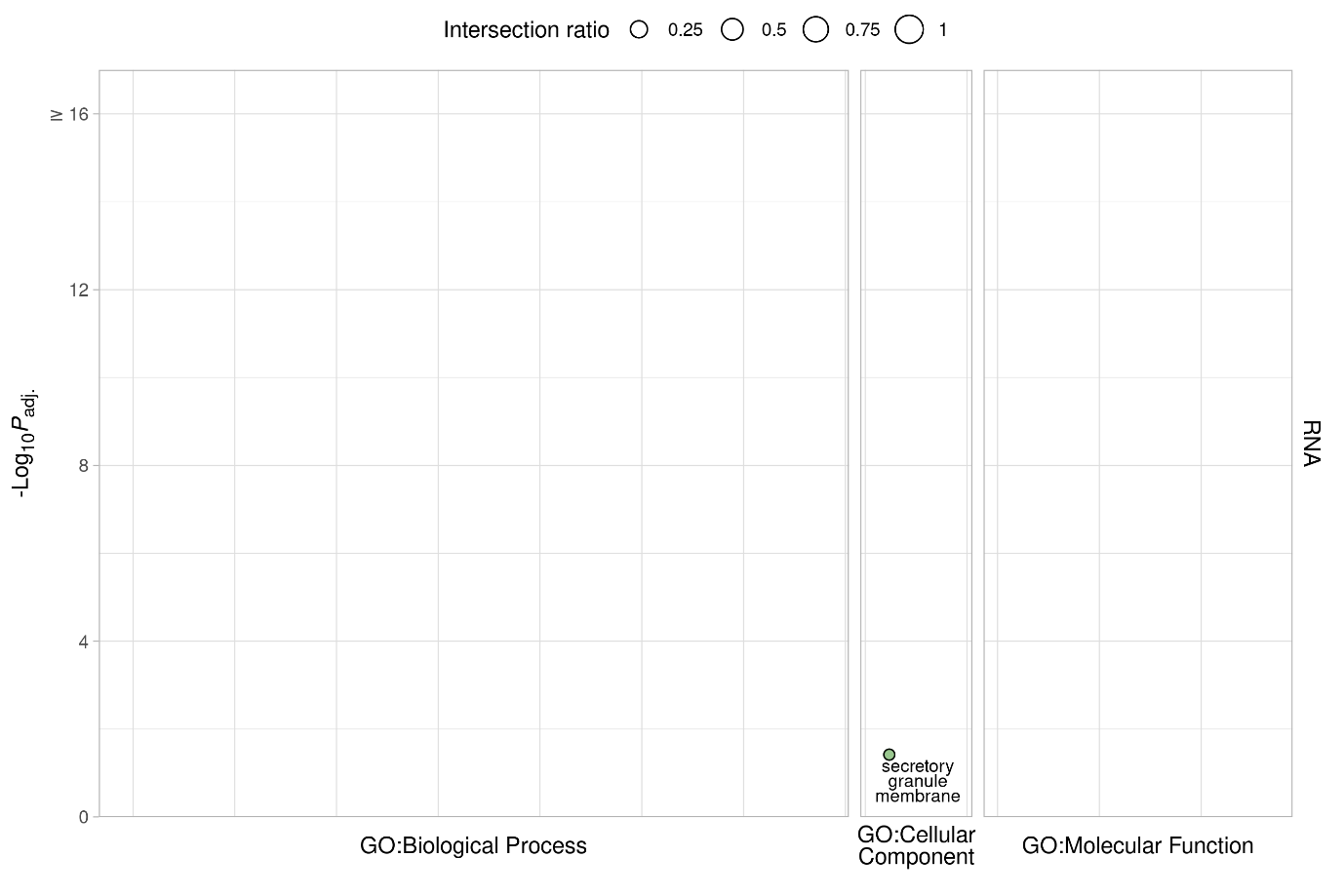

**Fig S28.** g:Profiler functional enrichment of significantly differentially expressed genes in the RNA-seq response contrasts with no background data set specified. Each dot represents a significantly enriched GO term with the size indicating the ratio of the intersection between the term and the introgressed genes. The *y*-axis shows the -log_10_*P*_adj._ value up to a maximum of 16 and the panels along the *y*-axis and colours indicate the module. The panels along the *x*-axis indicate the source of the term and the position within the panels groups terms from the same GO subtree. The top driver GO terms up to a maximum of ten are indicated with a black outline and label.

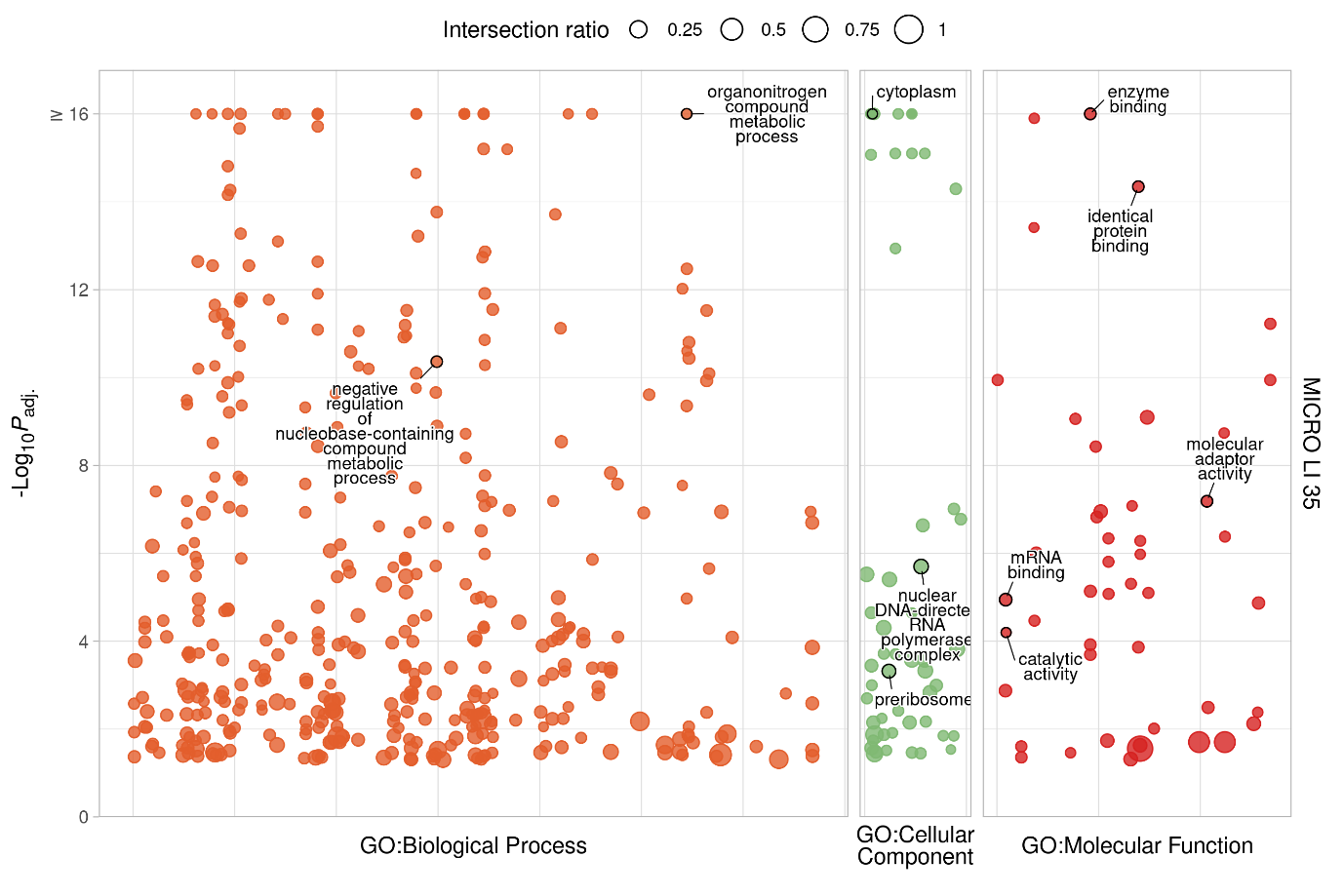

**Fig S29.** g:Profiler functional enrichment of the genes in the MICRO LI 35 functional module with no background data set specified. Each dot represents a significantly enriched GO term with the size indicating the ratio of the intersection between the term and the introgressed genes. The *y*-axis shows the -log_10_*P*_adj._ value up to a maximum of 16 and the panels along the *y*-axis and colours indicate the module. The panels along the *x*-axis indicate the source of the term and the position within the panels groups terms from the same GO subtree. The top driver GO terms up to a maximum of ten are indicated with a black outline and label.

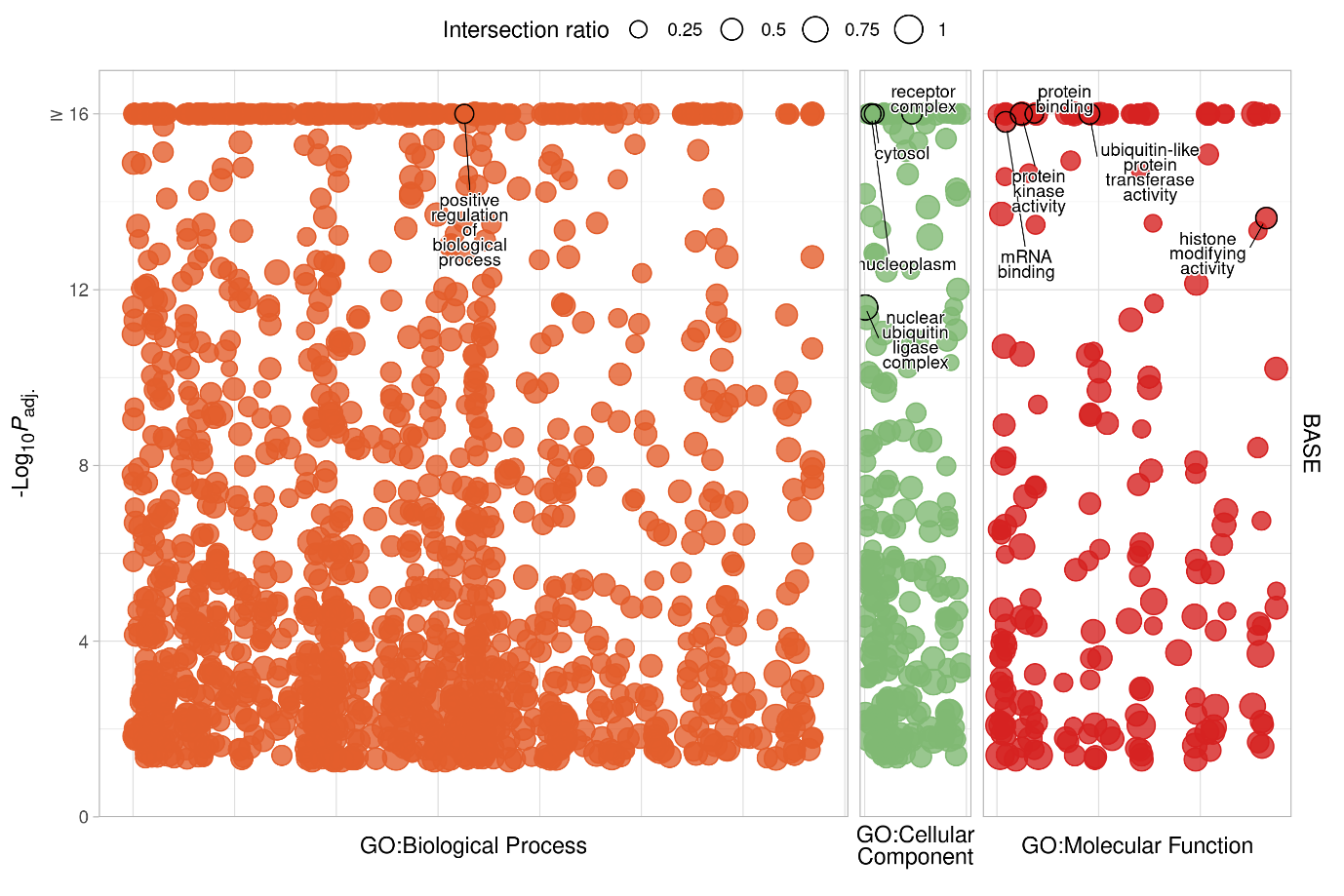

**Fig S30.** g:Profiler functional enrichment of the genes in the base network with no background data set specified. Each dot represents a significantly enriched GO term with the size indicating the ratio of the intersection between the term and the introgressed genes. The *y*-axis shows the -log_10_*P*_adj._ value up to a maximum of 16 and the panels along the *y*-axis and colours indicate the module. The panels along the *x*-axis indicate the source of the term and the position within the panels groups terms from the same GO subtree. The top driver GO terms up to a maximum of ten are indicated with a black outline and label.
